# The floral illusion: A parasitic beetle mimics the scent of flowers to attract bees

**DOI:** 10.64898/2026.01.15.699641

**Authors:** Ryan M. Alam, Danny Kessler, Heiko Vogel, Katrin Luck, Anja David, Maritta Kunert, Martin Kaltenpoth, Sarah E. O’Connor, Tobias G. Köllner

## Abstract

Animals are not known to biosynthesize floral signals to manipulate pollinators, although such mimicry could profoundly shape plant-pollinator interactions. Larvae of the poisonous European blister beetle *Meloe proscarabaeus* parasitize multiple solitary bee species, yet the mechanism enabling host attraction has remained unresolved. Here we show that these larvae lure bees by emitting a bouquet of volatile compounds that closely resembles floral scent. Chemical analyses reveal a complex blend of monoterpenoids derived from (*S*)-linalool, a ubiquitous floral volatile. Behavioral assays demonstrate that these compounds function as floral cues, eliciting attraction in bees. Transcriptomic and functional analyses identify cytochrome P450 enzymes that oxidize (*S*)-linalool, demonstrating that larvae biosynthesize these plant-like volatiles *de novo*. Together, these findings broaden the scope of interkingdom chemical mimicry and uncover a striking form of sensory deception in which an insect chemically assumes the signal identity of a flower, revealing that animals can evolve biosynthetic pathways to exploit plant–pollinator communication.

## Main

Mimicry, broadly defined as deceptive resemblance, is a widespread evolutionary strategy utilized across the tree of life^1^. Since Bates’ seminal observations of visual mimicry among Amazonian butterflies in 1862^2^, the concept has broadened to include deception across all major sensory modalities, including visual^3–5^, tactile^6^, acoustic^7^, and chemical^8–12^. Although many studies focus on mimicry in predator-prey or host-parasite interactions^13–15^, deceptive resemblance also plays key roles in other interspecific relationships. One striking example is phoresy, where one organism (the phoront) uses another (the host) for transport to essential resources^16^. Phoronts frequently evolve morphological, behavioral, and/or chemical traits that imitate benign or mutualistic species, deceiving hosts into providing transport^8, 17–19^. Moreover, phoresy has been proposed as an evolutionary precursor to parasitism in certain systems^16^, and complete dependence on a host for dispersal can further drive the emergence of obligate parasitic relationships^8^.

Blister beetles (Coleoptera: Meloidae) exemplify the intersection of mimicry, phoresy, and parasitism. Most species parasitize solitary bees^20^. In the subfamily Meloinae, gravid females deposit large egg clutches underground; the eclosed larvae (triungula) are highly mobile and either climb nearby vegetation to await hosts (**Fig. 1A**)^8, 21, 22^ or, in non-phoretic species, actively search for bee nests on the soil surface^23^. These larvae commonly target solitary bees, rapidly attaching to them, and are then transferred to the bee nest. Once in the nest, triungula dismount, consume the bee egg and provisions, and later complete several distinct developmental stages before emerging as adults the following year^24^.

**Fig. 1.**
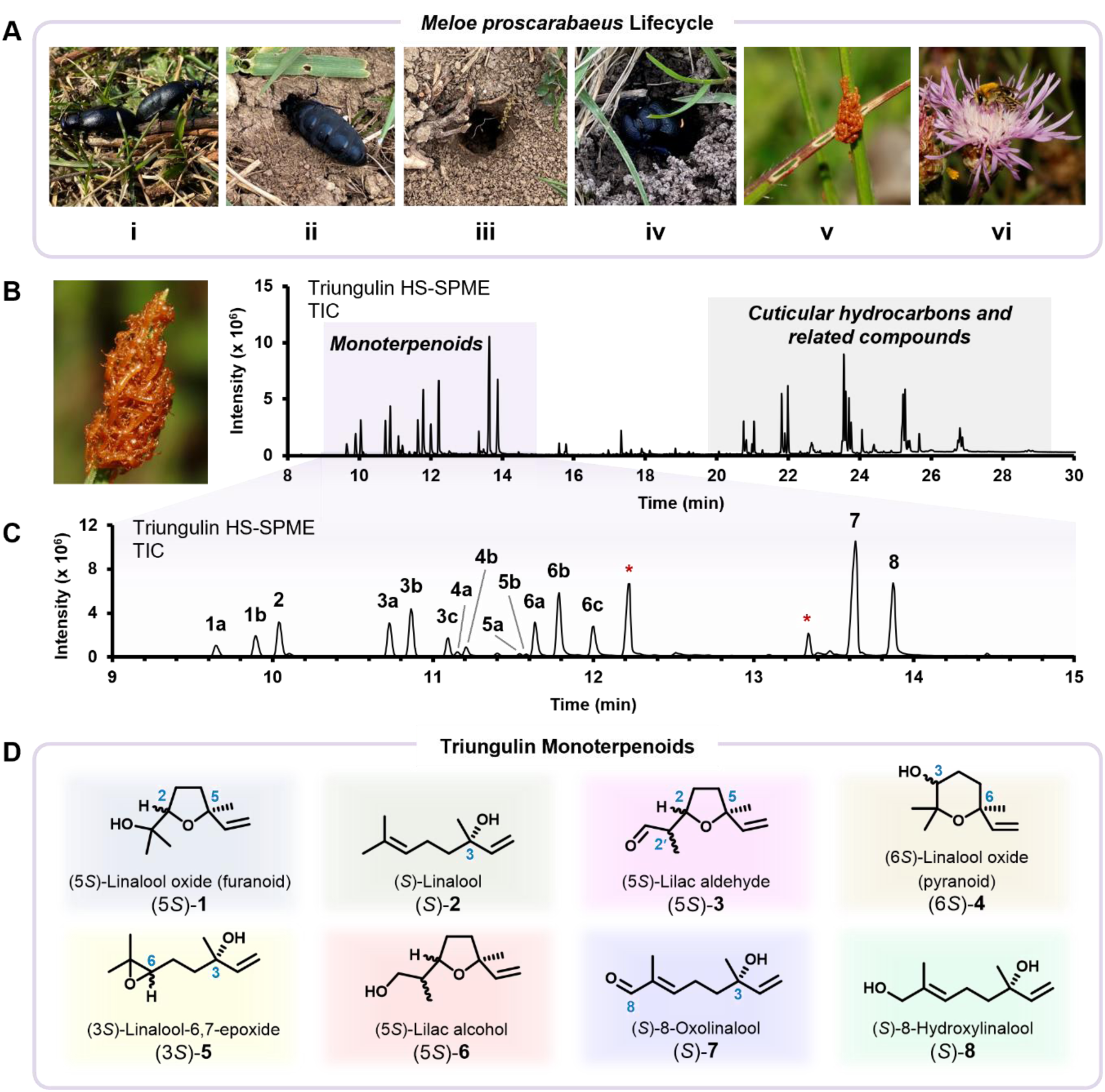
Lifecycle and volatile emissions of *Meloe proscarabaeus* triungula. (**A**) Lifecycle of the toxic European Black oil beetle *M*. *proscarabaeus*. After emerging and mating in the spring (i), gravid females dig underground chambers (ii–iii) and oviposit thousands of yellow eggs (iv). Triungula hatch after several weeks and form conspicuous orange aggregations on vegetation (v), where they await contact with a solitary bee host that inadvertently carries them to their nest to complete their development to adulthood (vi). (**B**) Total ion current (TIC) chromatogram of triungulin volatile profile, obtained *via* HS-SPME collection and achiral GC-EI-MS analysis. (**C**) Alongside long-chain hydrocarbons (t_r_ 20.95–29.22 min), *M*. *proscarabaeus* triungula emit a complex bouquet of floral-scent monoterpenoids (t_r_ 9.83–14.05 min, **1**–**8**). Peak identification: **1a** and **1b**) (5*S*)-linalool oxide (furanoid)^‡^, **2**) (*S*)-linalool, **3a**–**c**) (5*S*)-lilac aldehyde^‡^, **4a** and **4b**) (6*S*)-linalool oxide (pyranoid)^‡^, **5a** and **5b**) (3*S*)-linalool-6,7-epoxide^‡^, **6a**–**c**) (5*S*)-lilac alcohol^‡^, **7**) (*S*)-8-oxolinalool, **8**) (*S*)-8-hydroxylinalool. *Background; ^‡^Mixture of stereoisomers. (**D**) Structural and stereochemical assignments of identified monoterpene volatiles were confirmed by comparison with synthetic standards using achiral and chiral GC-EI-MS.

Host-attraction strategies in kleptoparasitic phoretic *Meloe* species vary and often exploit host-specific sensory cues. For instance, *Meloe franciscanus* triungula in North America aggregate on vegetation and emit a blend of female bee sex pheromones to specifically attract male *Habropoda* spp.^8, 22^. Alternatively, *M. strigulosus* larvae station themselves on flowers, where they intercept foraging pollinators^21^. In contrast, triungula of the European Black oil beetle *M*. *proscarabaeus* typically form conspicuous orange aggregations on grasses and exhibit little host specificity, suggesting a generalist phoretic strategy^25^. This behavioral divergence led us to hypothesize that *M*. *proscarabaeus* triungula use an alternative volatile cue to attract a broad range of pollinator hosts. Here, we demonstrate that *M*. *proscarabaeus* larvae emit a complex bouquet of floral-scent monoterpenoids to deceive foraging bees. This discovery reveals a previously undescribed form of interkingdom chemical mimicry and expands the conceptual framework of sensory deception and phoretic host attraction.

## Larvae emit floral monoterpenoids

To characterize volatile emissions from *M*. *proscarabaeus* triungula, adult beetles were collected in early spring (February–April 2024–25) from Jena, Thuringia, Germany, and maintained under controlled conditions for mating and oviposition (**Supplementary Fig. 1**). After approximately three weeks, egg clutches hatched synchronously, and aggregating triungula were harvested for volatile analysis (**Supplementary Fig. 2**). Headspace solid-phase microextraction (HS-SPME) coupled with achiral gas chromatography-electron ionization-mass spectrometry (GC-EI-MS) revealed a complex volatile profile, comprising more than thirty C_19_–C_27_ saturated and unsaturated long-chain hydrocarbons (t_r_ 20.95–29.22 min; **Fig. 1B**; **Supplementary Fig. 3**), similar to compounds previously reported in *M*. *franciscanus* larvae^22^. However, in addition to these hydrocarbons, we detected several structurally distinct monoterpenoids (t_r_ 9.83–14.05 min; **Figs. 1C** and **D**), including *cis*– and *trans*-diastereoisomers of linalool oxide (furanoid) (**1**), linalool (**2**), multiple stereoisomers of lilac aldehyde (**3**), linalool oxide (pyranoid) (**4**), linalool-6,7-epoxide (**5**), and lilac alcohol (**6**), and 8-oxolinalool (**7**) and 8-hydroxylinalool (**8**). Compound identities (**1**–**8**) were confirmed by comparison with synthesized reference standards (**Supplementary Fig. 4**).

Chiral GC-EI-MS analysis revealed that *M*. *proscarabaeus* triungula exclusively emit the (*S*)-enantiomer of linalool ((*S*)-**2**) (**Supplementary Figs. 5** and **6**) along with a range of (*S*)-**2**-derived metabolites bearing conserved stereochemistry at the C-3 position (**Fig. 1D**, **Supplementary Figs. S7**–**S20**). Absolute configurations were verified by comparison with synthetic reference standards derived from *rac*-, (*R*)-, and (*S*)-**2**. In total, triungula emitted 17 structurally distinct monoterpenoids (**1**–**8**), all of which are known floral volatiles widespread among angiosperms^26, 27^ such as *Berberis vulgaris*, *Prunus* spp., and *Salix* spp.^28–31^, which serve as the main food source for pollinating bees in the spring, when *M. proscarabaeus* triungula also emerge. However, except for (*S*)-**2**^32^ and linalool oxide **1**^33^, none of these monoterpenoids have been previously reported to be produced in an insect.

## Floral-scent mimicry attracts bees

Several monoterpenoids emitted by *M*. *proscarabaeus* triungula—including (*S*)-linalool ((*S*)-**2**) and its derivatives (**1** and **3**–**8**)—are common floral volatiles known to attract pollinators, including solitary wild bees^28–31, 34^. We hypothesized that by production and emission of these volatiles, triungula closely mimic a floral scent that, in turn, can lure a broad range of phoretic hosts. Although *M*. *proscarabaeus* larvae primarily parasitize ground-nesting solitary bees^20^, triungula have also been recovered from nests of social bee species^20, 35^, suggesting that volatile emission by these larvae may also mediate phoretic interactions with these unsuitable social insect hosts that can eject the larvae from the nest after transmission^36^. Notably, the observation of *M. proscarabaeus* triungula on both suitable and unsuitable bee hosts is consistent with a general attraction strategy.

To test whether *M*. *proscarabaeus* triungulin monoterpenoids attract solitary bees, we first conducted dual-choice olfactometer assays with polylectic *Osmia bicornis,* a commercially available wild bee species (**Fig. 2A**; **Supplementary Fig. 21**). Live triungula, emitting a complex blend of monoterpenoids and long-chain hydrocarbons attracted both male and female *O*. *bicornis* (**Fig. 2B**). Similarly, volatile extracts obtained *via* dynamic headspace volatile (DHV) collection—which contained markedly reduced amounts of long-chain hydrocarbons—also attracted both sexes of *O*. *bicornis* (**Fig. 2C**). Notably, DHV collection captured the full triungulin monoterpenoid profile, whereas parallel solvent-based extractions failed to recover these volatiles (**Supplementary Fig. 22**).

**Fig. 2.**
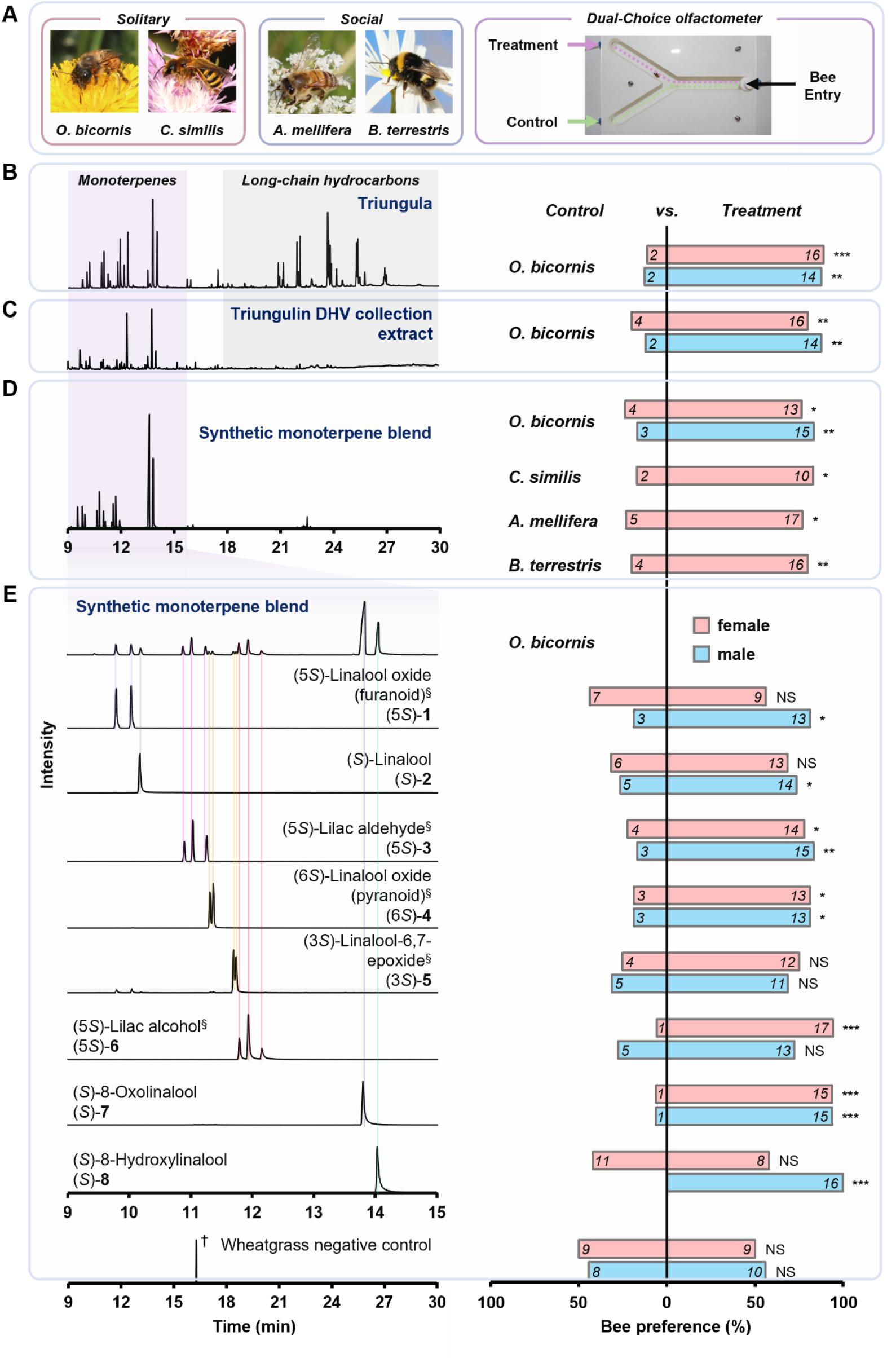
Triungulin-derived monoterpenoids attract solitary and social bees. (**A**) Dual-choice Y-tube behavioral assays were performed with solitary (*Osmia bicornis* and *Colletes similis*) and social (*Apis mellifera* and *Bombus terrestris*) bee species. (**B**) Both male and female *O*. *bicornis* were attracted to live triungula. (**C**) Volatile extracts from aggregating triungula, obtained *via* dynamic headspace volatile (DHV) collection, also attracted male and female *O*. *bicornis*. (**D**) A synthetic blend of (*S*)-linalool ((*S*)-**2**)-derived monoterpenoids (**1**–**8**) elicited significant attraction from female *C. similis*, *A*. *mellifera*, *B*. *terrestris*, and both sexes of *O*. *bicornis*. (**E**) Behavioral assays with individual monoterpenoids showed sex-specific responses in *O*. *bicornis*. Control assays with wheat in both Y-tube arms confirmed no side bias. All bees, except *C*. *similis*, were flower-naïve. Numbers in bars indicate absolute choices. χ² test: \*\*\**p* < 0.001, \*\**p* < 0.01, \**p* < 0.05, NS = not significant. ^§^Mixture of stereoisomers; ^†^Geranylacetone; tentatively identified using the NIST MS-Library v. 3.0 (2023).

As previously noted, *M*. *franciscanus* larvae attract male solitary bees by emitting a blend of long-chain hydrocarbons that mimic female bee sex pheromones^22^. To determine whether triungulin-emitted monoterpenoids can independently elicit attraction—without the influence of co-occurring long-chain hydrocarbons—we exposed male and female *O*. *bicornis* to a synthetic blend of (*S*)-linalool derivatives (compounds **1**–**8**; **Fig. 2C**; **Supplementary Fig. 23A**) lacking these hydrocarbons. Both sexes were significantly attracted to this blend (χ² test, *p* < 0.05). In contrast, only males, but not females, responded to an analogous blend of synthetic (*R*)-linalool-derived monoterpenoids (**Supplementary Fig. 23B**), highlighting the importance of the natural stereoisomeric composition of the triungulin volatile bouquet.

Considering that *O*. *bicornis* is likely an unsuitable host for *M*. *proscarabaeus* triungula—as it typically nests in pre-existing cavities in wood, hollow stems, loess, clay, or masonry^37^—we next tested the synthetic (*S*)-**2-**derived blend in dual-choice assays with wild-caught females of *Colletes similis*, a ground-nesting solitary bee, and with workers of the eusocial generalists *Bombus terrestris* and *Apis mellifera* (**Fig. 2D**). These assays confirmed the attraction of *C. similis* and additionally revealed that triungula can also attract unsuitable social bee hosts, a capacity that may enhance their phoretic dispersal. For instance, *M*. *strigulosus* triungula have been observed to detach from unsuitable hosts during inter-floral movements^21^. Such a “hop-on, hop-off” strategy likely increases dispersal efficiency and may similarly occur in *M. proscarabaeus*.

While the sex pheromones of *A*. *mellifera* and *B*. *terrestris* have been described previously as long-chain hydrocarbon derivatives^38, 39^ and do not include monoterpenoids, certain colletid bees^40^, such as *Colletes cunicularius*^32^, use (*S*)-**2** as a mate attractant. However, we could not detect monoterpenoids in either sex of *C. similis* or *O. bicornis* (**Supplementary Fig. 24**). Together with the observation that triungulin monoterpenoids attract several distantly related bee species, this supports the conclusion that bee attraction to triungulin monoterpenoids is based on floral scent mimicry rather than the imitation of bee sex pheromones. In addition, the bright orange coloration and shape of the larval aggregates may also visually mimic floral signals, a possibility further supported by the contrasting appearances of *M*. *franciscanus* and closely related *M*. *violaceus*^41^ triungula, which are markedly darker^8, 20^. Indeed, increasing evidence indicates that mimics exploit multiple sensory channels simultaneously, enhancing the effectiveness of the deception^42^.

We next investigated each compound (**1**–**8**) individually in dual-choice assays using naïve male and female *O*. *bicornis* (**Fig. 2D**). All monoterpenoids except (3*S*)-linalool-6,7-epoxide ((3*S*)-**5**) and (5*S*)-lilac alcohol ((5*S*)-**6**) elicited significant attraction in males (χ² test, 0.001 > *p* < 0.05). Females were not attracted to (3*S*)-**5** and, unlike males, showed no preference for (*S*)-**2**, (5*S*)-linalool oxide (furanoid) ((5*S*)-**1**), or (*S*)-8-hydroxylinalool ((*S*)-**8**) (*p* > 0.05). However, (5*S*)-lilac aldehyde ((5*S*)-**3**), (6*S*)-linalool oxide (pyranoid) ((6*S*-**4**), (5*S*-**6**), and (*S*)**-8-**oxolinalool ((*S*)-**7**) were significantly attractive to females (0.01 > *p* < 0.05). The observed attraction of female bees to the monoterpenoid blend as well as several of the individual components support the hypothesis that mimicry of floral scent in *M*. *proscarabaeus* larvae can bypass male bees and directly lure female hosts, unlike triungula of *M*. *franciscanus* who only attract intermediary male *Habropoda* hosts and then have to move to females during copulation in order to reach the nest. The direct attraction of females by *M*. *proscarabaeus* larvae likely enhances their chances of nest entry and successful development.

Considering that *M. proscarabaeus* triungula usually aggregate on vegetation but sometimes climb to reach flowers, we hypothesized that larval monoterpenoids have a dual role: facilitating both host attraction *and* larval aggregation. Indeed, olfactometric assays showed that triungula were significantly attracted to a synthetic blend of (*S*)-linalool-derived monoterpenoids (**1**–**8**), supporting a dual role for these volatiles in bee attraction and larval clustering (**Supplementary Fig. 25**, **Supplementary Table 1**). This result suggests an intriguing evolutionary scenario in which triungula likely originally relied on floral volatiles as cues to locate flowers, where they could await pollinating insect hosts. The emission of such scents by the larvae may have initially merely amplified the natural floral signal. However, ultimately, this also led to the aggregation of triungula, allowing clustered larvae to generate a substantially stronger bouquet than solitary individuals and thereby reduce their dependence on actual flowers. In fact, *M*. *proscarabaeus* larvae typically emerge in the spring near bee nests in areas that have few ground-flowering plants and where early-flowering shrubs and trees, which act as main food source for pollinating bees, are often several hundred meters away. Thus, flower mimicry may have helped *M. proscarabaeus* to occupy new ecological niches. Remarkably, this host-attraction strategy also targets a fundamental behavioral mechanism essential for the survival of pollinating insects, limiting their potential for rapid coevolutionary escape.

## Triungula biosynthesize monoterpenoids

Following eclosion, larvae of *M*. *proscarabaeus* consistently emit floral-scent monoterpenoids **1**–**8**. However, it remained unclear whether these volatiles were sequestered, maternally transferred, or biosynthesized *de novo*. However, as larvae do not feed on plants following eclosion^20^, the sequestration of floral monoterpenoids seems rather unlikely. To determine if larval monoterpenoids **1**–**8** were maternally derived, we analyzed the volatile profiles of adult females and their eggs, using HS-SPME GC-EI-MS (**Supplementary Figs. 26** and **27**). However, apart from the emission of (*S*)-linalool ((*S*)-**2**) by eggs (**Supplementary Fig. 28**), we could detect no other floral monoterpenoids in eggs or adults. This result, together with the conserved (*S*) stereochemistry at the linalool C-3 position among compounds **1** and **3**–**8**, led us to hypothesize that triungula biosynthesize these monoterpenoids from (*S*)-**2** (**Fig. 3A**).

**Fig. 3.**
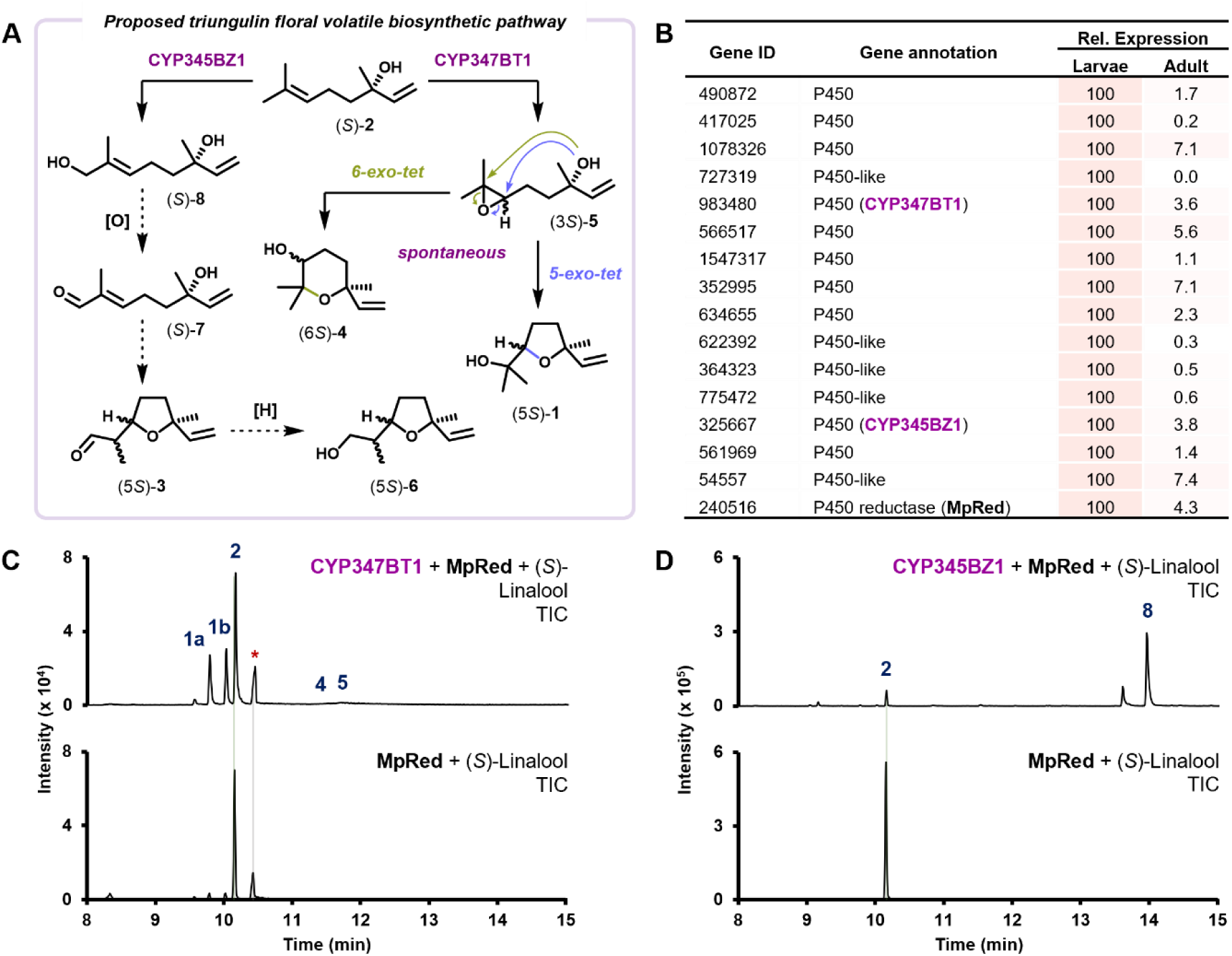
Triungula biosynthesize a range of (*S*)-linalool-derived floral monoterpenoids. (**A**) Proposed biosynthetic pathway of floral-scent volatiles in triungula showing early P450-mediated oxidation of central precursor (*S*)-linalool ((*S*)-**2**) to epoxide (3*S*)-**5** and diol (*S*)-**8**. Spontaneous ring-closure of labile epoxide (3*S*)-**5** subsequently affords linalool oxides (5*S*)-**1** and (6*S*)-**4**, while further oxidation of (*S*)-**8** furnishes intermediate (*S*)-**7** that cyclizes to lilac aldehyde (5*S*)-**3**. Enzymatic reduction of aldehyde (5*S*)-**3** generates alcohol (5*S*)-**6**. (**B**) Differential gene expression analysis identified a cytochrome P450 reductase (MpRed) and P450-encoding transcripts upregulated in larvae relative to adult females. Relative (Rel.) expression values are based on reads per kilobase of transcript per million reads mapped (RPKM) values obtained by RNA-seq (for absolute RPKM values, see **Supplementary Table 4**). (**C**) Functional characterization of CYP347BT1 in *S*. *cerevisiae* expressing either CYP347BT1 and MpRed or MpRed alone in the presence of (*S*)-**2**. Reaction products were analyzed using GC-EI-MS. Peak identification: **1a** and **1b**) (5*S*)-linalool oxide (furanoid), **2**) (*S*)-linalool, **4**) (6*S*)-linalool oxide (pyranoid), **5**) (3*S*)-linalool-6,7-epoxide. Note: Only traces of (6*S*)-**4** and (3*S*)-**5** were detected. *Background. (**D**) Functional characterization of CYP345BZ1 in *S*. *cerevisiae* expressing either CYP345BZ1 and MpRed or MpRed alone in the presence of (*S*)-**2**. Reaction products were analyzed using GC-EI-MS. Peak identification: **8**) (*S*)-8-hydroxylinalool.

As compounds **2** and **5**, and **3**, **7**, and **8** are known intermediates in the biosynthesis of linalool oxides (**1** and **4**) and lilac alcohol (**6**), respectively, we hypothesized that two cytochrome P450 (P450) enzymes generate structural diversity within the triungulin monoterpenoid bouquet by oxidizing (*S*)-linalool ((*S*)-**2**) to (*S*)-**8** and (3*S*)-**5**. To test this, we extracted RNA from aggregating triungula and adult females, and generated a *de novo* transcriptome assembly. Differential gene expression analysis identified 15 transcripts annotated as P450 or P450-like, with elevated expression in triungula but not in adult females, along with a putative P450 reductase (*MpRed*) (**Fig. 3B**).

Each candidate P450 was individually co-expressed with *MpRed* in *Saccharomyces cerevisiae* and screened for biosynthetic activity *in vivo* in the presence of (*S*)-**2**. One candidate, designated CYP347BT1 according to P450 nomenclature, epoxidized (*S*)-**2** to primarily yield linalool oxide (furanoid) stereoisomers, (2*S*,5*S*)– and (2*R*,5*S*)-**1**, along with trace amounts of (3*S*)-linalool-6,7-epoxide ((3*S*)-**5**) and linalool oxide (pyranoid) diastereomers (6*S*)-**4** (**Fig. 3C**). Another candidate designated CYP345BZ1, catalyzed the terminal allylic C-H hydroxylation of (*S*)-**2** to produce (*S*)-8-hydroxylinalool (*S*)-**8** (**Fig. 3D**). Together, these results confirm that *M*. *proscarabaeus* triungula independently biosynthesize floral-scent monoterpenoids.

Consistent with *in vivo* assays, HS-SPME and DHV collection extracts of triungula contained relatively low amounts of (3*S*)-**5** and (6*S*)-**4**, compared to linalool oxide (5*S*)-**1** (**Fig. 1B** and **C**). Furthermore, GC-EI-MS analysis indicated that (3*S*)-**5** undergoes both 5– and 6-exo-tet intramolecular cyclization to form (5*S*)-**1** and (6*S*)-**4**, respectively, during analysis (**Supplementary Fig. 29**), congruent with the spontaneous degradation of synthetic (5*S*)-**5** to (5*S*)-**1** and (6*S*)-**4** over time (**Supplementary Fig. 30**). The formation of both (2*R*,5*S*)– and (2*S*,5*S*)-**1** stereoisomers from (*S*)-**2** in the presence of CYP347BT1 and MpRed further suggests that CYP347BT1-mediated epoxidation proceeds non-stereoselectively, initially generating labile intermediates (3*S*,6*S*)-*and* (3*S*,6*R*)-**5** from (*S*)-**2**.

## Conclusion

Our results reveal a hitherto undescribed form of interkingdom aggressive chemical mimicry in which *M*. *proscarabaeus* triungula collectively emit a complex bouquet of floral-scent monoterpenoids to lure solitary and social bees. Behavioral assays show that these volatiles act as floral-scent mimics rather than pheromone analogs, enabling attraction of male and female solitary bees, promoting successful phoretic dispersal, and coordinating larval aggregation. We identify two P450 enzymes, CYP347BT1 and CYP345BZ1, that generate committed biosynthetic precursors ((3*S*)-**5** and (*S*)-**8**) responsible for the suite of (*S*)-linalool-derived monoterpenoids (**1**, **3**, **4**, **6**, and **7**). Together, these findings broaden the evolutionary framework of mimicry by demonstrating how an insect can independently produce plant-like signals to manipulate inter– and intraspecific interactions.

## Data availability

All data required to evaluate the conclusions proposed herein are present in the Manuscript or associated Supplementary Information. Genes characterized in this study are deposited in the NCBI GenBank with the following accession numbers: MpRed (PX569140), CYP347BT1 (PX569141), and CYP345BZ1 (PX569142) and sequencing data are available at https://doi.org/10.17617/3.7TXB4J.

## Acknowledgements

We thank Daniel Veit and Angela Lehmann (MPI-ICE) for the design and construction of volatile collection and behavioral bioassay apparatuses, Sarah Heinicke (MPI-ICE) for assistance with GC-EI-MS data acquisition, Prof. David Nelson (UTHSC) for kindly annotating P450 genes, *CYP345BZ1* and *CYP347BT1*, and Drs Klaus Gase, Mohamed Omar Kamileen, Maite Colinas, Song Wu (MPI-ICE), and Hannah M. Rowland (MPI-ICE/UOL) for helpful discussion.

## Funding

This work was supported by funding from the Max Planck Society.

## Author contributions

Conceptualization: R.M.A., M.Ka., S.E.OC., and T.G.K.; Data curation: R.M.A., H.V., and T.G.K.; Formal analysis: R.M.A., D.K., H.V., and T.G.K.; Funding acquisition: S.E.OC.; Investigation: R.M.A., D.K., H.V., K.L., and A.D.; Methodology: R.M.A., D.K., M.Ku., and T.G.K.; Project administration: R.M.A. and T.G.K.; Resources: S.E.OC.; Supervision: S.E.OC. and T.G.K.; Validation: R.M.A., D.K., K.L., and T.G.K.; Visualization: R.M.A. and T.G.K.; Writing – original draft: R.M.A. and T.G.K.; Writing – review and editing: R.M.A., D.K., H.V., M.Ka., S.E.OC., and T.G.K.

## Ethics declarations

Competing interest:

The authors declare that they have no competing interests.

## Notes

### Competing Interest Statement

The authors have declared no competing interest.

