## Supplementary Information for "The floral illusion: A parasitic beetle mimics the scent of flowers to attract bees"

**This PDF file includes:**

Materials and Methods
Supplementary Figs. 1–30
Supplementary Tables 1–4
Supplementary References (43–56)

### Materials and Methods

#### *Meloe proscarabaeus* triungulin cultivation and collection

Adult male and female specimens of *Meloe proscarabaeus* were collected in the field during the Spring of 2024 and 2025 in Jena, Thuringia, Germany, with permission from Untere Naturschutzbehörde Jena (permit UNB-AS-BE-AG-2023-01) (**Supplementary Fig. 1A**). Adult beetles were maintained on a diet of fresh *Vicia faba, Trifolium pratense,* and *Triticum aestivum*, and housed together in tanks that were kept in a ventilated greenhouse out of direct sunlight (temperature: daytime, *ca*. 10–15 °C; night, ≮ 5 °C). Following oviposition (**Supplementary Fig. 1B**) and subsequent eclosion (**Supplementary Fig. 1C**), live triungula were collected for analysis when required.

#### Triungulin volatile collection and analysis

Preliminary sampling of *M. proscarabaeus* triungulin volatiles was performed using a solid-phase microextraction (SPME) fiber (100 μm poly(dimethylsiloxane) [PDMS], Supelco®, Darmstadt, Germany) that had been conditioned at 250 °C for 10 min prior to volatile collection. Briefly, *ca*. 50 live triungula were collected in a 1.5 mL septum-sealed amber glass vial that had previously been heated to 200 °C for 30 min and subsequently cooled (prior to use) and then a pre-conditioned SPME fiber (heated to 250 °C for 10 min, prior to use) was inserted *via* the septum of the vial into the headspace (**Supplementary Fig. 2A**) and left *in situ* for 3 h, before being directly submitted for GC-EI-MS analysis (*vide infra*). For dynamic headspace volatile (DHV) collection, live triungula (*ca*. 250) were collected in a 25 mL glass bottle (previously heated to 200 °C for 30 min and subsequently cooled, prior to use). The bottle was connected to an in-house closed loop sampling apparatus (**Supplementary Fig. 2B**) which comprised a lid fitted with an airtight inlet and outlet containing a removable Porapak™ P cartridge (**Supplementary Fig. 2C**) (connected *via* an electronic air circulation pump). Following 3 h of volatile collection the Porapak™ cartridge was removed and flushed with $CH_2Cl_2$ (100 μL x 3). For liquid solvent extractions, approximately 50 triungula were immersed in hexane, MeOH, or $CH_2Cl_2$. After standing for 1 h at room temperature, extracts were taken off with a glass pipette. All volatile extracts were carefully stored at −20 °C, until required.

**Gas chromatography-electron ionization-mass spectrometry (GC-EI-MS)**

Gas chromatography electron ionization mass spectrometry (GC-EI-MS) analyses were performed using an Agilent 8890 Series gas chromatograph coupled with an Agilent 5977B single quadrupole mass selective detector (MSD) (Agilent Technologies, CA 95051, United States). Unless stated otherwise, samples were prepared using $CH_2Cl_2$ as sample solvent to give a final concentration of *ca*. 10 µg/mL. Chromatographic analyses were performed using helium as the carrier gas applied at a constant flow rate with an injection volume of 1 µL in splitless mode and MS data acquisition was carried out in EI mode using an electron ionization energy of 70 eV. HS-SPME analyses were carried out by inserting the SPME fiber directly into the injection port of the GC.

Achiral chromatographic separations were performed using the following method:

Separation method: A

Achiral chromatographic separations were performed using a Zebron ZB-5™ column (5% phenyl, 95% dimethylpolysiloxane; 30 m length, 0.25 mm i.d., 0.25 µm $d_f$, 10 m pre-column, Phenomenex, Aschaffenburg, Germany). Carrier gas was applied at a constant flow rate of 1 mL/min. The inlet and transfer line temperatures were 250 °C and 280 °C, respectively. For chromatographic separation, an initial column oven temperature of 40 °C was maintained for 2 min and then increased by 10 °C/min to 280 °C and then held for 2 min. Solvent delay was set to 5 min. The ion source temperature was 230 °C. MS data acquisition was carried out in scan mode (mass range, 50–450 m/z).

Chiral GC-EI-MS analyses were carried out using the following separation methods:

Separation method: B

Employing a β-DEX™ 225 column (25% 2,3-di-*O*-acetyl-6-*O*-TBDMS-β-cyclodextrin in SPB-20 poly(20% phenyl/80% dimethylsiloxane); 30 m length, 0.25 mm i.d., 0.25 $d_f$; Supelco), both the inlet and transfer line temperatures were set to 220 °C. Carrier gas was applied at a constant flow rate of 1.2 mL/min. For chromatographic separation, an initial column temperature of 40 °C was held for 2 min and then increased by 2 °C/min to 200 °C. After holding at 200 °C for 5 min, the temperature was then decreased by 7 °C/min to 40 °C. The solvent delay was set to 12 min and MS data acquisition was carried out in scan mode (mass range, 50–500 m/z).

Separation method: C

Using a Hydrodex β-6TBDM column (Heptakis(2,3-di-*O*-methyl-6-*O*-tert-butyldimethylsilyl)-β-cyclodextrin; 25 m length, 0.25 mm i.d.; Macherey-Nagel, Düren, Germany), both the inlet and transfer line temperatures were set to 220 °C. The carrier gas was applied at a constant flow rate of 1.5 mL/min and the solvent delay was set to 15 min. For chromatographic separation, the column temperature was initially held at 60 °C for 2 min and then increased by 2 °C/min to 120 °C. After holding the temperature at 120 °C for 10 min, the column temperature was increased by 5 °C/min to 180 °C, held for 5 min, and then decreased by 10 °C/min to 60 °C. MS data acquisition was carried out in scan mode (mass range, 50–350 m/z).

Separation method: D

Employing a β-DEX™ 110 column (10% permethylated β-cyclodextrin in SPB-35 poly(35% diphenyl/65% dimethylsiloxane; 30 m length, 0.25 mm i.d., 0.25 $d_f$, Supelco), both the inlet and transfer line temperatures were set to 220 °C with a constant carrier gas flow rate of 1.2 mL/min. Chromatographic separation was performed using an initial column temperature of 50 °C that was held for 2 min and then increased by 2 °C/min to 150 °C. After holding at 150 °C for 10 min, the temperature was further increased by 4 °C/min to 180 °C, held for 5 min, and then decreased by 7 °C/min to 50 °C. The solvent delay was set to 15 min and MS data acquisition was carried out in scan mode (mass range, 50–350 m/z).

Separation method: E

Using a β-DEX™ 225 column (25% 2,3-di-*O*-acetyl-6-*O*-TBDMS-β-cyclodextrin in SPB-20 poly(20% phenyl/80% dimethylsiloxane); 30 m length, 0.25 mm i.d., 0.25 $d_f$; Supelco), both the inlet and transfer line temperatures were set to 220 °C. Carrier gas was applied at a constant flow rate of 1.2 mL/min. For chromatographic separation, an initial column temperature of 40 °C was held for 2 min and then increased by 8 °C/min to 120 °C. After holding at 120 °C for 60 min, the temperature was increased by 2 °C/min to 160 °C and then further increased by 8 °C/min to 200

°C, held for 5 min, and subsequently decreased by 8 °C/min to 40 °C. The solvent delay was set to 12 min and MS data acquisition was carried out in scan mode (mass range, 50–500 m/z).

Separation method: F

Utilizing a β-DEX™ 225 column (25% 2,3-di-*O*-acetyl-6-*O*-TBDMS-β-cyclodextrin in SPB-20 poly(20% phenyl/80% dimethylsiloxane); 30 m length, 0.25 mm i.d., 0.25 $d_f$; Supelco), both the inlet and transfer line temperatures were set to 220 °C. Carrier gas was applied at a constant flow rate of 1.2 mL/min. For chromatographic separation, an initial column temperature of 40 °C was held for 2 min and then increased by 8 °C/min to 130 °C. After holding at 130 °C for 90 min, the temperature was increased by 2 °C/min to 180 °C and then further increased by 8 °C/min to 200 °C, held for 5 min, and subsequently decreased by 8 °C/min to 40 °C. The solvent delay was set to 12 min and MS data acquisition was carried out in scan mode (mass range, 50–500 m/z).

### Chemical synthesis of monoterpenoid volatiles

#### General note

All reagents were obtained from commercial sources and used without any further purification, unless otherwise specified. Powdered 4 Å molecular sieves were dried (200 °C for 48 h), prior to use. Room temperature ranged between 18–22 °C. Thin layer chromatography (TLC) was carried out on aluminum-backed silica gel 60 $F_{254}$ plates (Merck) that were visualized with Hanessian's stain and/or using $UV_{254\ nm}$ light detection. Flash column chromatography was performed using Merck silica gel 60 (particle size 0.040–0.063 mm, density 0.8 $g/cm^3$).

### Nuclear magnetic resonance (NMR) spectroscopy

Nuclear magnetic resonance (NMR) spectra were recorded on a Bruker Avance® III HD 400 MHz spectrometer (Bruker Biospin GmbH, Rheinstetten, Germany) at room temperature (*ca*. 293 K), using $CDCl_3$ as sample solvent. Chemical shift values ($\delta_H$) are reported in parts per million (ppm) relative to residual solvent ($\delta_H$ 7.26 [$C\textit{H}Cl_3$]) and coupling constants (*J*) are expressed in Hertz (Hz), in the following format; chemical shift value (multiplicity, coupling constant, integration). $^1H$ NMR spectral data are described, using the following abbreviations; brs (broad singlet), s (singlet), t (triplet), q (quartet), dd (doublet of doublets), ddd (doublet of doublets of doublets), appd (apparent doublet), and m (multiplet).

**Synthesis of (*S*)-linalool ((*S*)-2)**

Following a reported route [43, 44], (*S*)-**2** was prepared over three steps from geraniol (**S1**).

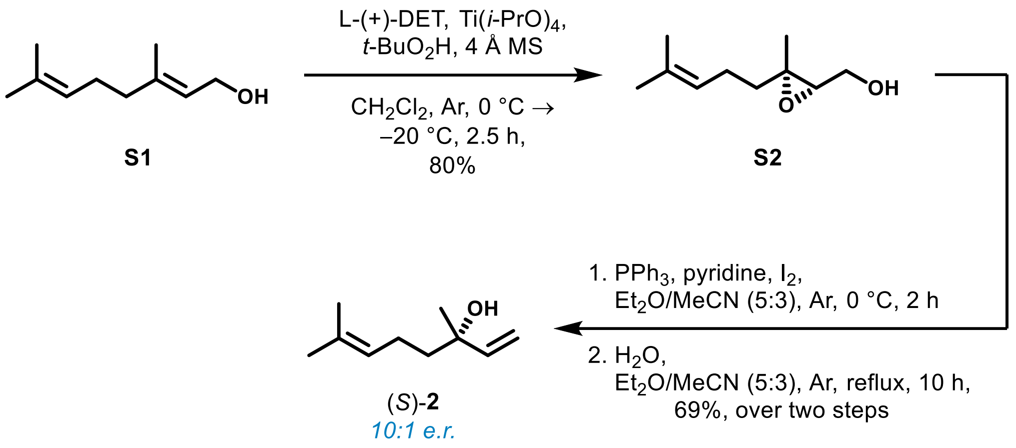


[(2*S*,3*S*)-3-Methyl-3-(4-methylpent-3-en-1-yl)oxiran-2-yl]methanol (**S2**)

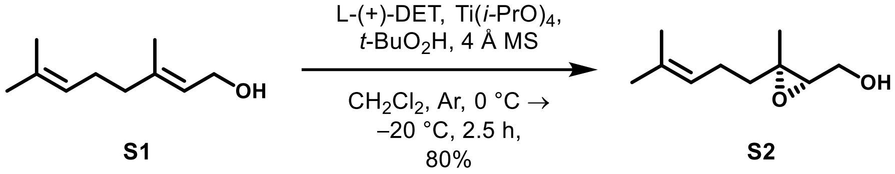


To a dry 250 mL two-neck round bottom flask fitted with a dropping funnel under an Ar atmosphere was added powdered 4 Å molecular sieves (1.8 g) and anhydrous $CH_2Cl_2$ (50 mL). Upon cooling to −20 °C, the suspension was sequentially charged with L-(+)-diethyl tartrate (1.66 mL, 9.72 mmol), Ti(*i*-PrO)$_4$ (1.92 mL, 6.48 mmol), and *t*-butyl hydroperoxide (5.5 M in *n*-decane, 17.68 mL, 97.25 mmol). After stirring at −20 °C for 30 min, to the mixture was then slowly added geraniol (**S1**, 3.16 mL, 18.21 mmol) *via* dropping funnel. Reaction monitoring using TLC (1:3, EtOAc/ petroleum ether [b.p. 40–60 °C]) confirmed complete consumption of allylic alcohol **S1**, after stirring at −20 °C for a further 2.5 h. The reaction mass was gradually warmed to 0 °C and then charged with 30% (w/v) NaOH in sat. aq. $NaHCO_3$ (20 mL). After stirring at 0 °C for 30 min, the mixture was warmed to room temperature and further diluted with $H_2O$ (20 mL). The resulting biphasic suspension was passed through a short pad of Celite®. Following phase separation, the aqueous layer was repeatedly extracted with $CH_2Cl_2$ (20 mL x 3). The combined organic layers were dried over anhydrous $MgSO_4$ and carefully concentrated under reduced pressure gave a residue that was subsequently subjected to short path vacuum distillation to afford an oil, which was then purified by gradient flash column chromatography (0–25% EtOAc in petroleum ether [b.p. 40–60 °C]) to yield title compound **S2** as a colorless oil (2.480 g, 80%): $R_f$ 0.40 (1:3, EtOAc/petroleum ether [b.p. 40–60 °C]); $^1$H NMR ($CDCl_3$, 400 MHz) $\delta$ 5.09–5.06 (m, 1H), 3.83–3.79 (m, 1H), 3.70–3.65 (m, 1H), 2.97 (dd, J = 6.7, 4.3 Hz, 1H), 2.08 (q, J = 7.7 Hz, 2H), 1.87 (brs, 1H), 1.71–1.63 (overlapping m, 1H), 1.68 (s, 3H), 1.60 (s, 3H), 1.50–1.43 (m, 1H), 1.29 (s, 3H). Physical and spectral data agreed with those reported previously [45].

(*S*)-Linalool ((*S*)-**2**)

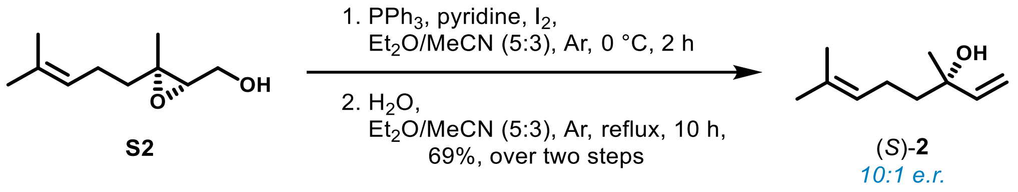


To a dry 250 mL round bottom flask, containing an ice-cold solution of **S2** (2.465 g, 14.48 mmol) in a mixture of anhydrous $Et_2O$ (50 mL) and MeCN (30 mL) under an Ar atmosphere, was sequentially added triphenylphosphine (11.394 g, 43.44 mmol), anhydrous pyridine (4.67 mL, 57.92 mmol), and $I_2$ (5.513 g, 21.72 mmol). After stirring at 0 °C for 2 h, the reaction mass was charged with $H_2O$ (0.26 mL) and heated to reflux (45 °C, oil bath) for 10 h. Upon cooling to room temperature, the resulting mixture was treated with 20% (w/v) aq. $Na_2S_2O_3$ (8 mL) and sat. aq. $NaHCO_3$ (8 mL). Following separation, the aqueous layer was extracted with $Et_2O$ (10 mL x 3) and the combined organic phases were washed with 1 M aq. HCl (10 mL), sat. aq. $NaHCO_3$ (10 mL), $H_2O$ (10 mL), and brine (10 mL). Careful concentration of the organic phase gave an oily residue that was further purified by gradient flash column chromatography (0–10% $Et_2O$ in pentane) to afford title compound (*S*)-**2** (10:1 e.r.) as a colorless oil (1.541 g, 69% [over two steps from **S2**]): $R_f$ 0.50 (1:4, $Et_2O$/pentane); $^1$H NMR ($CDCl_3$, 400 MHz) $\delta$ 5.91 (dd, J = 17.3, 10.8 Hz, 1H), 5.21 (dd, J = 17.3, 1.2 Hz, 1H), 5.14–5.10 (m, 1H), 5.06 (dd, J = 10.8, 1.2 Hz, 1H), 2.10 (m, 2H), 1.68 (s, 3H), 1.60 (overlapping s, 3H), 1.60–1.56 (overlapping m, 2H), 1.28 (s, 3H); GC-MS (EI) *m*/*z* (%) 71 (100), 93 (85), 55 (50), 69 (42), 80 (35), 121 (27), 83 (19), 67 (19), 92 (17), 79 (13). Physical and spectral data agreed with those reported previously [44].

Chiral GC-EI-MS extracted ion chromatograms (EIC) for (**A**) synthetic (*S*)-**2** (10:1 *e.r.*;), (**B**) (*R*)-**2**, and (**C**) *rac*-**2**. Separation method: B.

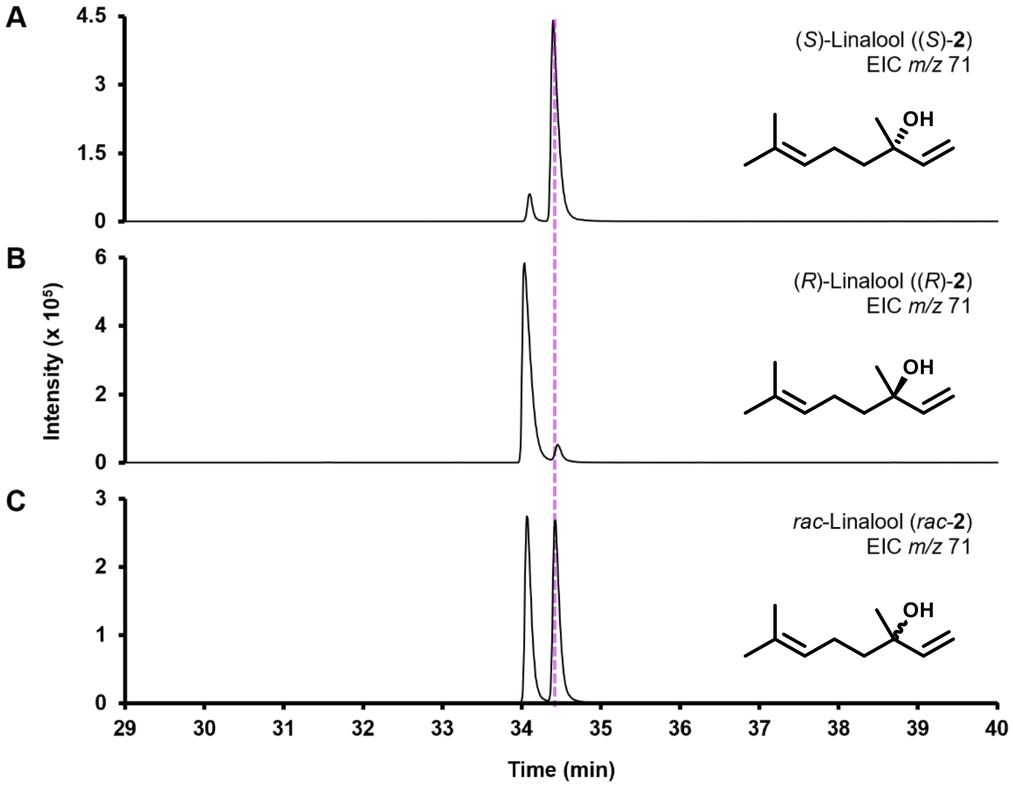

**Synthesis of linalool oxide (furanoid) (1) and (pyranoid) (4)**

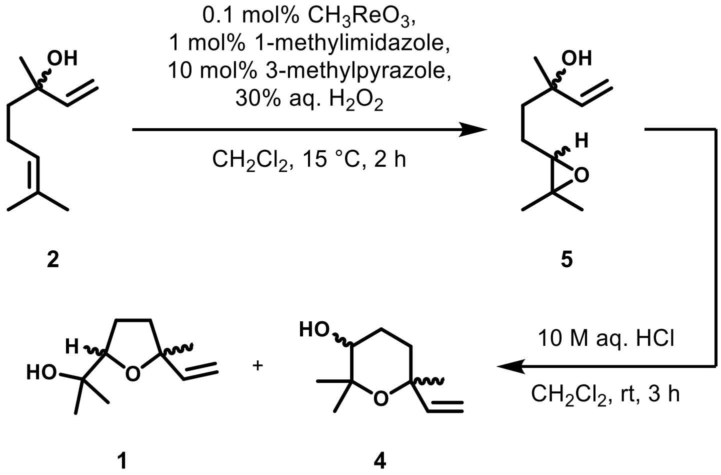


Linalool-6,7-epoxide (**5**, (3*S*)-**5**, and (3*R*)-**5**)

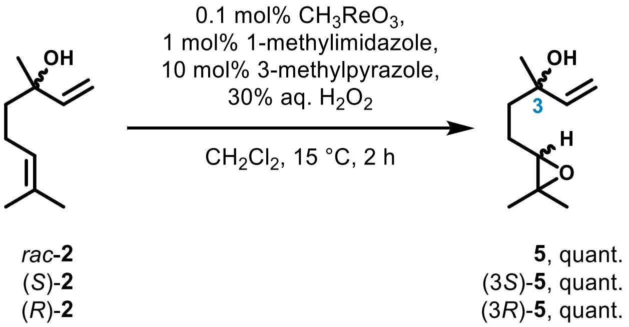


Briefly following a previously reported procedure [46], to a 10 mL round bottom flask containing *rac*-linalool (**2**, 1.80 mL, 10 mmol) in $CH_2Cl_2$ (5 mL) cooled to 15 °C, was added 3-methyl-1*H*-pyrazole (80 µL, 1 mmol), 1-methyl-1*H*-imidazole (8 µL, 0.1 mmol), and methyltrioxorhenium (2.5 mg, 0.01 mmol). Following the addition of 30% (w/w) aq. $H_2O_2$ (1.2 mL, 10.6 mmol), the resulting biphasic mixture was vigorously stirred for 2 h at 15 °C. The reaction mass was charged with brine (10 mL) and extracted with $CH_2Cl_2$ (5 mL x 3). The combined organic layers were washed with sat. aq. $Na_2S_2O_3$ (5 mL) and then dried over anhydrous $Na_2SO_4$. Careful concentration *in vacuo* gave title compound **5** as a colorless oil (1.701 g, quant.): $^1$H NMR ($CDCl_3$, 400 MHz) mixture of diastereoisomers* $\delta$ 5.93–5.84 (m, 1H), 5.24–5.19 (m, 1H), 5.08–5.04 (m, 1H), 2.74–2.71 (m, 1H), 1.79–1.49 (m, 4H), 1.30–1.29 (m, 3H), 1.26 (s, 3H). Physical and spectral data agreed with those reported previously [46]. *Both diastereoisomers of **5** are described.

Note: Following this procedure, (3*R*)-**5** and (3*S*)-**5** were analogously prepared from (*R*)- and (*S*)-linalool, respectively.

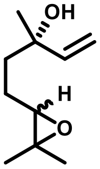

(3*S*)-**5**, quant.

(3*S*)-Linalool-6,7-epoxide ((3*S*)-**5**) was isolated as a colorless oil (1.701 g [quant.] from 1.543 g [10 mmol] of (*S*)-**2**): GC-MS (EI) *m*/*z* (%) *2*) 71 (100), 68 (77), 59 (75), 67 (54), 72 (51), 55 (38), 97 (36), 79 (30), 69 (28), 85 (25), *4*) 71 (100), 68 (78), 59 (75), 67 (54), 72 (49), 97 (37), 55 (36), 79 (31), 69 (28), 85 (25). Physical and spectral data agreed with those reported previously [46].

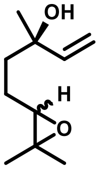

(3*R*)-**5**, quant.

(3*R*)-Linalool-6,7-epoxide ((3*R*)-**5**) was isolated as a colorless oil (1.699 g [quant.] from 1.543 g [10 mmol] of (*R*)-**2**): Physical and spectral data agreed with those reported previously [46].

Chiral GC-EI-MS EIC chromatograms for (**A**) synthetic linalool-6,7-epoxide **5** (peaks 1–4), (**B**) (3*S*)-**5** (peaks 2 and 4), and (**C**) (3*R*)-**5** (peaks 1 and 3). Separation method: C. *Mixture of stereoisomers.

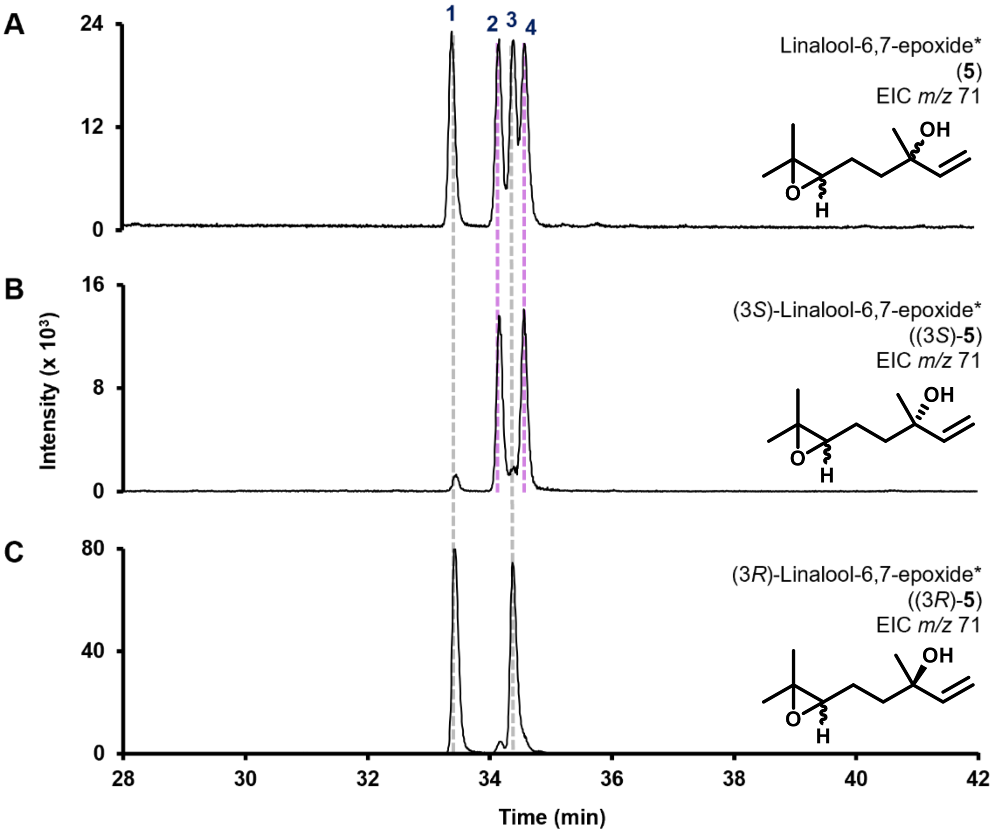

Linalool oxide (furanoid) ((5*S*)-**1** and (5*R*)-**1**) and linalool oxide (pyranoid) ((6*S*)-**4** and (6*R*)-**4**)

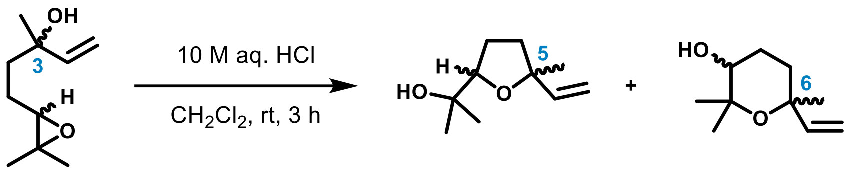


(3*S*)-**5**
(3*R*)-**5**

(5*S*)-**1**, 79%
(5*R*)-**1**, 75%

(6*S*)-**4**,10%
(6*R*)-**4**, 8%

To a 50 mL round bottom flask containing a solution of epoxide (3*S*)-**5** (1.7 g, 10 mmol) dissolved in $CH_2Cl_2$ (5 mL) was added 10 M aq. HCl (50 µL, 0.5 mmol). After stirring at room temperature for 3 h (complete consumption of starting material **5** was confirmed by TLC [$Et_2O$/pentane, 3:1]), the reaction mass was partitioned between $CH_2Cl_2$ (10 mL) and $H_2O$ (20 mL). The resulting aqueous layer was repeatedly extracted with $CH_2Cl_2$ (10 mL x 3) and the combined organic phases were then dried over anhydrous $Na_2SO_4$ and carefully concentrated under reduced pressure to afford a colorless oil. Purification of the residue, using gradient flash column chromatography (0–20% $Et_2O$ in pentane), afforded title compounds **1** and **4**.

Note: Following this procedure, (5*S*)-linalool oxide (furanoid) ((5*S*)-**1**) and (6*S*)-linalool oxide (pyranoid) ((6*S*)-**4**) were analogously prepared from (3*S*)-**5**. Similarly, (5*R*)-**1** and (6*R*)-**4** were synthesized from epoxide (3*R*)-**5**.

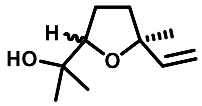


(5*S*)-**1**, 79%

(5*S*)-Linalool oxide (furanoid) ((5*S*)-**1**) was isolated as a colorless oil (1.342 g [79%] from 1.7 g [10 mmol] of (3*S*)-**5**): $R_f$ 0.23 ($Et_2O$/pentane, 1:4); $^1$H NMR ($CDCl_3$, 400 MHz) mixture of diastereoisomers* $\delta$ 5.97 (dd, $J$ = 17.4, 10.8 Hz, 0.5H), 5.87 (dd, $J$ = 17.3, 10.6 Hz, 0.5H), 5.21–5.16 (m, 1H), 5.02–4.98 (m, 1H), 3.86 (t, $J$ = 6.9 Hz, 0.5H), 3.80 (t, $J$ = 7.1 Hz, 0.5H), 2.00 (brs, 1H), 1.96–1.68 (m, 4H), 1.31 (s, 3H), 1.232 (s, 1.5H), 1.226 (s, 1.5H), 1.131 (s, 1.5H), 1.126 (s, 1.5H); GC-MS (EI) *m/z* (%) *2*) 59 (100), 94 (74), 111 (56), 93 (56), 68 (37), 55 (36), 67 (33), 81 (26), 83 (21), 79 (18), *4*) 59 (100), 94 (75), 93 (53), 111 (52), 68 (38), 55 (37), 67 (32), 81 (25), 83 (23), 79 (19). Physical and spectral data agreed with those reported previously [47]. *Both diastereoisomers of (5*S*)-**1** are described.

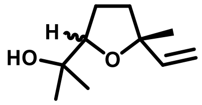

(5*R*)-**1**, 75%

(5*R*)-Linalool oxide (furanoid) ((5*R*)-**1**) was isolated as a colorless oil (1.273 mg [75%] from 1.697 g [9.97 mmol] of (3*R*)-**5**). Physical and spectral data agreed with those reported previously [47].

Chiral GC-EI-MS EIC chromatograms for (**A**) linalool oxide (furanoid) **1**, (**B**) (5*S*)-**1**, and (**C**) (5*R*)-**1**. Elution order of **1** stereoisomers: *1*) (2*R*,5*R*)-**1**, *2*) (2*S*,5*S*)-**1**, *3*) (2*S*,5*R*)-**1**, and *4*) (2*R*,5*S*)-**1** [48]. Separation method: D. *Mixture of stereoisomers.

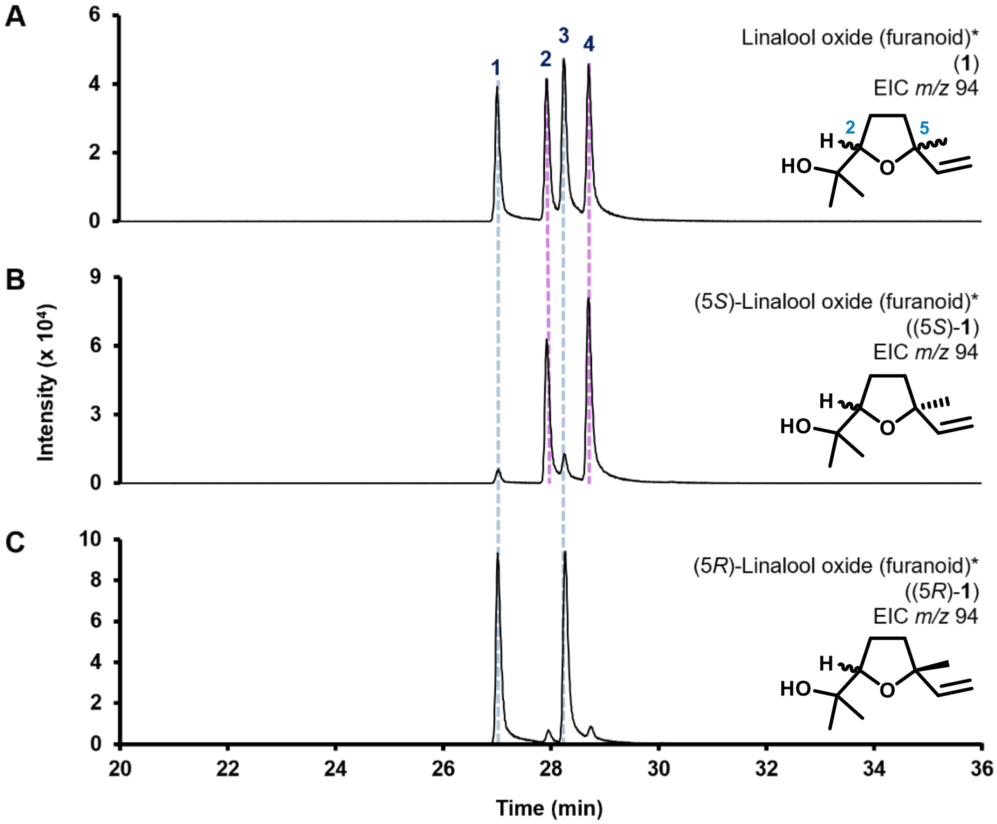

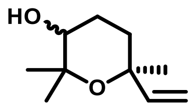

(6*S*)-**4**, 10%

(6*S*)-Linalool oxide (pyranoid) ((6*S*)-**4**) was isolated as a colorless wax (167 mg [10%] from 1.700 g [10 mmol] of (3*S*)-**5**): $R_f$ 0.13 ($Et_2O$/pentane, 1:4); $^{1}H$ NMR ($CDCl_3$, 400 MHz) mixture of diastereoisomers* $\delta$ 6.00–5.90 (m, 1H), 5.04–4.95 (m, 2H), 3.45–3.39 (m, 1H), 2.12 (dt, *J* = 13.6, 3.8 Hz, 0.5H), 2.02–1.94 (m, 0.5H), 1.84–1.53 (m, 4H), 1.25 (overlapping s, 1.5H), 1.24 (overlapping s, 1.5H), 1.22 (s, 3H), 1.169 (overlapping s, 1.5H), 1.161 (overlapping s, 1.5H); GC-MS (EI) *m*/*z* (%) *1*) 68 (100), 94 (75), 59 (61), 67 (57), 79 (12), 53 (12), 83 (12), 69 (12), 55 (11), 79 (11), *3*) 68 (100), 94 (75), 59 (60), 67 (52), 79 (23), 69 (14), 84 (13), 53 (12), 81 (12), 55 (11). Physical and spectral data agreed with those reported previously [49]. *Both diastereoisomers of **4** are described.

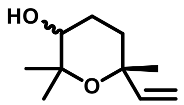

(6*R*)-**4**, 8%

(6*R*)-Linalool oxide (pyranoid) ((6*R*)-**4**) was isolated as a colorless wax (136 mg [8%] from 1.697 g [9.97 mmol] of (3*R*)-**5**). Physical and spectral data agreed with those reported previously [49].

Chiral GC-EI-MS EIC chromatograms for (**A**) linalool oxide (pyranoid) **4**, (**B**) (6*S*)-**4**, and (**C**) (6*R*)-**4**. Elution order of **4** stereoisomers: *1*) (3*R*,6*S*)-**4**, *2*) (3*S*,6*R*)-**4**, *3*) (3*S*,6*S*)-**4**, and *4*) (3*R*,6*R*)-**4** [48]. Separation method: D. *Mixture of stereoisomers.

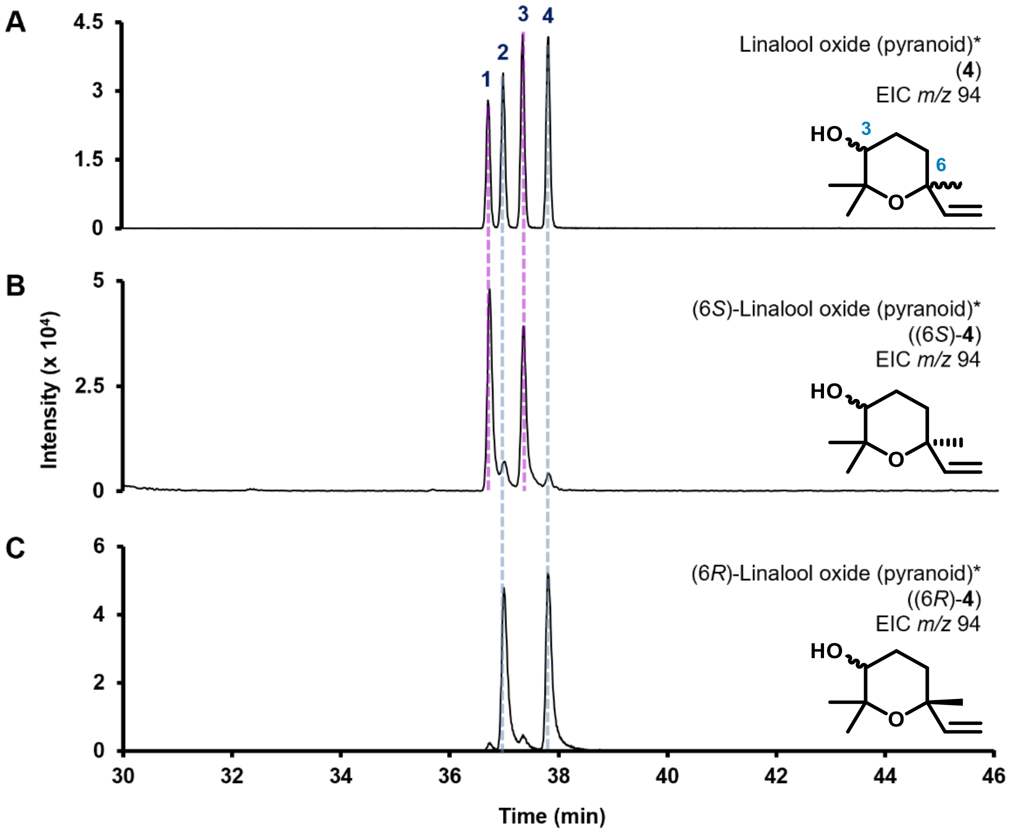

**Synthesis of 8-Oxolinalool (7)**

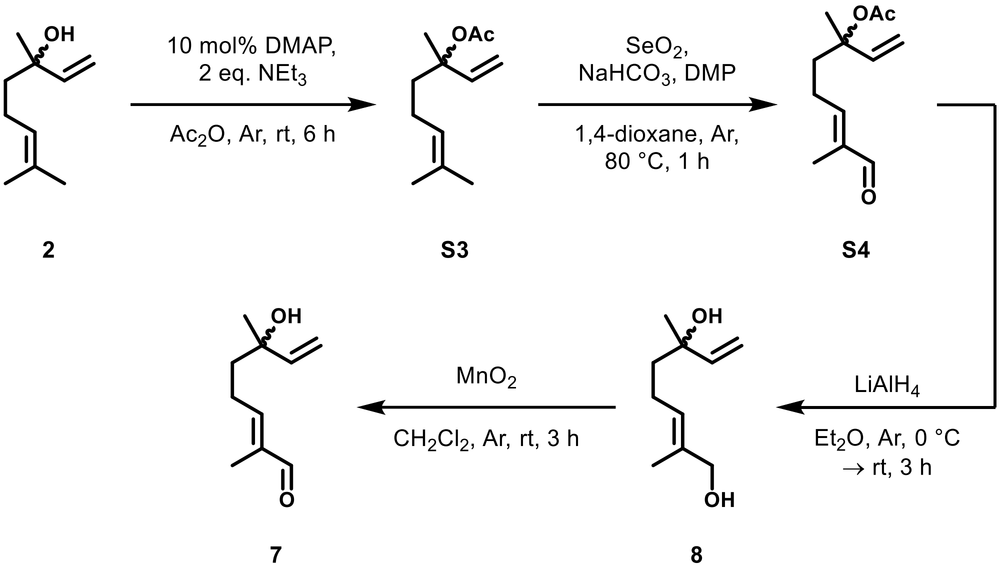


Linalyl acetate (**S3**)

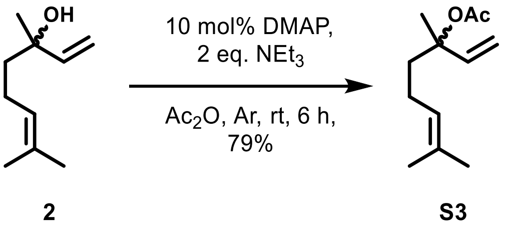


To a dry 100 mL round bottom flask under an Ar atmosphere containing *rac*-**2** (5 mL, 27.94 mmol) and 4-dimethylaminopyridine (341 mg, 2.79 mmol) was added acetic anhydride (5.28 mL, 55.88 mmol) and triethylamine (7.79 mL, 55.88 mmol). After stirring at room temperature for 6 h, the resulting deep amber solution was cooled to 0 °C, slowly charged with sat. aq. $NaHCO_3$ (100 mL), and allowed to stir for a further 30 min whilst warming to room temperature. The mixture was extracted with $Et_2O$ (25 mL x 3) and the combined organic layers were sequentially washed with sat aq. $NaHCO_3$ (25 mL x 3), 0.1 M aq. HCl (25 mL x 2), $H_2O$ (25 mL), and brine (25 mL). The organic phase was dried over anhydrous $Na_2SO_4$ and carefully concentrated under reduced pressure to afford a residue that was purified by gradient flash column chromatography (0–5% $Et_2O$ in hexane) to furnish acetate *rac*-**S3** as a colorless oil (4.332 g, 79%): $R_f$ 0.23 ($Et_2O$/hexane, 1:19); $^{1}$H NMR ($CDCl_3$, 400 MHz) $\delta$ 5.97 (dd, $J$ = 17.5, 11.0 Hz, 1H), 5.15 (dd, $J$ = 17.4, 0.9 Hz, 1H), 5.12 (dd, $J$ = 11.0, 0.9 Hz, 1H), 5.11–5.06 (overlapping m, 1H), 2.00 (s, 3H), 1.98–1.94 (m, 2H), 1.91–1.72 (m, 2H), 1.67 (appd, $J$ = 0.8 Hz, 3H), 1.59 (s, 3H), 1.54 (s, 3H); GC-MS (EI) *m/z*

(%) 93 (100), 121 (36), 80 (35), 69 (21), 92 (17), 71 (14), 91 (14), 79 (13), 94 (13), 136 (12). Physical and spectral data agreed with those reported previously [50].

Note: Following this procedure, (*S*)-**S3** and (*R*)-**S3** were analogously prepared from (*S*)- and (*R*)-**2**, respectively.

Achiral GC-EI-MS data for *rac*-**S3**, including (**A**) EIC chromatogram and (**B**) EI-MS spectrum. Separation method: A

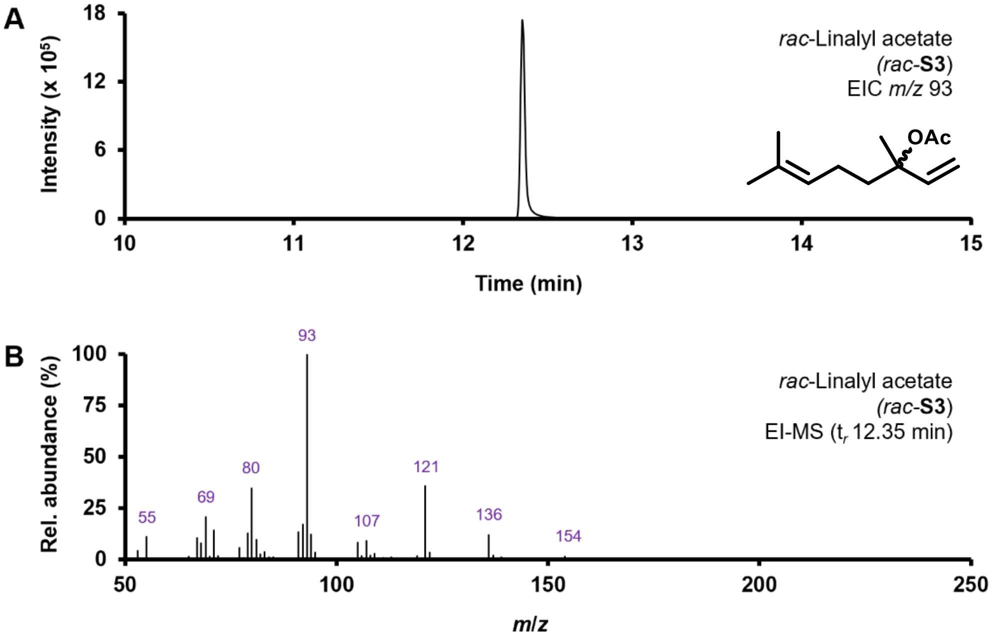

3,7-Dimethyl-8-oxoocta-1,6-dien-3-yl acetate (**S4**)

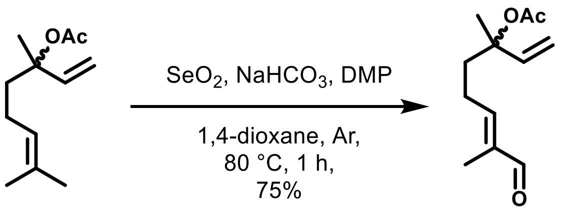


**S3** **S4**

To a dry 250 mL round bottom flask under an Ar atmosphere, containing acetate **S3** (4.044 g, 20.60 mmol) and anhydrous 1,4-dioxane (41.2 mL), was sequentially added $SeO_2$ (4.572 g, 41.20 mmol), $NaHCO_3$ (3.461 g, 41.20 mmol), and Dess-Martin periodinane (17.475 g, 41.20 mmol). The beige suspension was heated to 80 °C (oil bath) for 1 h and then cooled to room temperature. The resulting burgundy suspension was passed through a short pad of Celite® and the wet cake was repeatedly washed with portions of $Et_2O$ (20 mL x 5). Upon cooling to 0 °C, the combined filtrate was slowly poured onto a rapidly stirring ice-cold mixture of sat. aq. $NaHCO_3$ (50 mL) and sat. aq. $Na_2S_2O_3$ (50 mL). After warming to room temperature and stirring for 15 min, the mixture was extracted with $Et_2O$ (50 mL x 3). The combined organic layers were washed with brine (20 mL) and dried over anhydrous $Na_2SO_4$. Careful concentration under reduced pressure afforded a residue that was distilled *in vacuo* (b.p. 147 °C/19 mbar) to afford α,β-unsaturated aldehyde *rac*-**S4** as a bright yellow oil (3.248 g, 75%): $R_f$ 0.27 ($Et_2O$/pentane, 1:4); $^1$H NMR ($CDCl_3$, 400 MHz) $\delta$ 9.38 (s, 1H), 6.48–6.44 (m, 1H), 5.95 (dd, $J$ = 17.5, 11.0 Hz, 1H), 5.21–5.15 (m, 2H), 2.36 (q, $J$ = 7.8 Hz, 2H), 2.12–2.04 (m, 1H), 2.01 (s, 3H), 1.94–1.86 (m, 1H), 1.73 (m, 3H), 1.58 (m, 3H); GC-MS (EI) $m/z$ (%) 71 (100), 82 (98), 84 (67), 150 (66), 93 (63), 121 (63), 95 (60), 80 (52), 107 (49), 108 (46). Physical and spectral data agreed with those reported previously [51].

Note: Following this procedure, (3*S*)-**S4** and (3*R*)-**S4** were analogously prepared from (3*S*)- and (3*R*)-**S3**, respectively.

Achiral GC-EI-MS data for *rac*-**S4**, including (**A**) EIC chromatogram and (**B**) EI-MS spectrum. Separation method: A

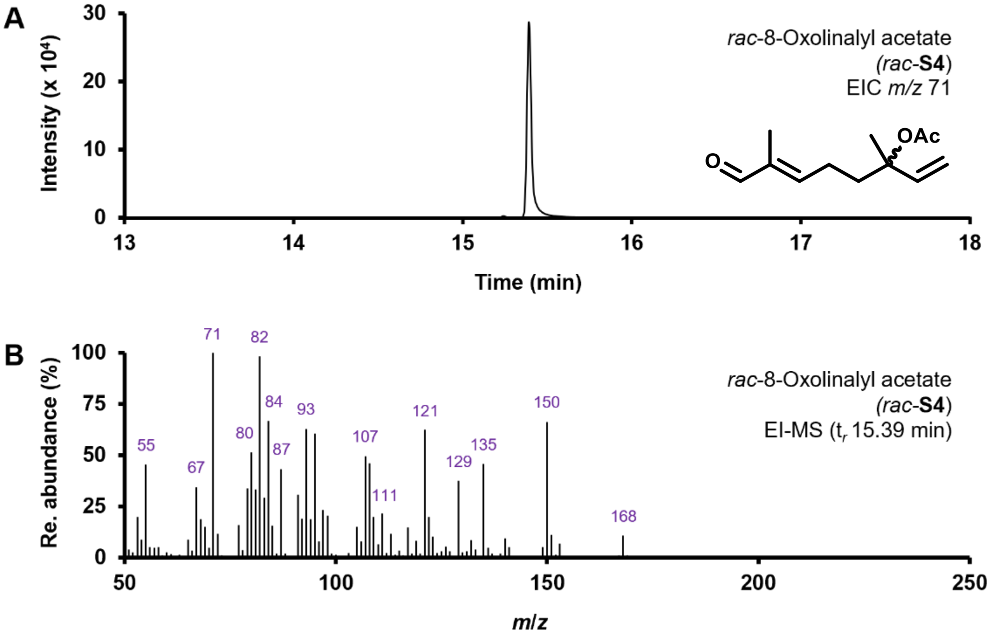

8-Hydroxylinalool (**8**)

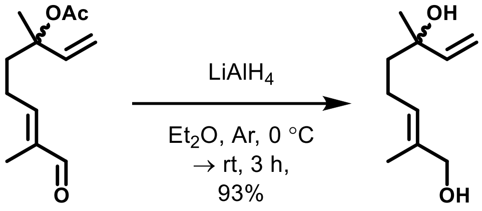


**S4** **8**

To a dry 25 mL round bottom flask under an Ar atmosphere, containing a suspension of $LiAlH_4$ (783 mg, 20.64 mmol) in anhydrous $Et_2O$ (5 mL) cooled to 0 °C, was added dropwise a solution of aldehyde **S4** (868 mg, 4.13 mmol) in anhydrous $Et_2O$ (5 mL). After warming to room temperature and stirring for 3 h (complete consumption of starting material **S4** was confirmed by TLC [$Et_2O$/pentane, 1:1]), the reaction mass was cooled to 0 °C and then cautiously treated dropwise with $H_2O$ (0.78 mL), 15% (w/v) aq. NaOH (0.78 mL), and $H_2O$ (2.34 mL). The resulting grey suspension was further diluted with $Et_2O$ (10 mL) and stirred at room temperature for 15 min. Filtration of the colorless suspension through a short pad of Celite® gave a colorless filtrate that was dried over anhydrous $MgSO_4$ and carefully concentrated *in vacuo* to afford a light yellow residue which was then purified by gradient flash column chromatography (0–100% $Et_2O$ in pentane) to yield diol **8** as a colorless oil (654 mg, 93%): $R_f$ 0.34 ($Et_2O$); $^1$H NMR ($CDCl_3$, 400 MHz) $\delta$ 5.91 (dd, $J$ = 17.3, 10.7 Hz, 1H), 5.43–5.39 (m, 1H), 5.21 (dd, $J$ = 17.4, 1.2 Hz, 1H), 5.07 (dd, $J$ = 10.8, 1.2 Hz, 1H), 3.98 (s, 2H), 2.15–2.01 (m, 2H), 1.65 (s, 3H), 1.61–1.55 (m, 4H), 1.29 (s, 3H); GC-MS (EI) *m/z* (%) 71 (100), 67 (81), 55 (39), 68 (39), 82 (35), 93 (29), 79 (28), 81 (26), 84 (25), 110 (23). Physical and spectral data agreed with those reported previously [52].

Note: Following this procedure, (*S*)-**8** and (*R*)-**8** were analogously prepared from (*S*)- and (*R*)-**S4**, respectively.

Chiral GC-EI-MS EIC chromatograms for (**A**) *rac*-8-hydroxylinalool (*rac*-**8**), (**B**) (*S*)-**8**, and (**C**) (*R*)-**8**. Separation method: E.

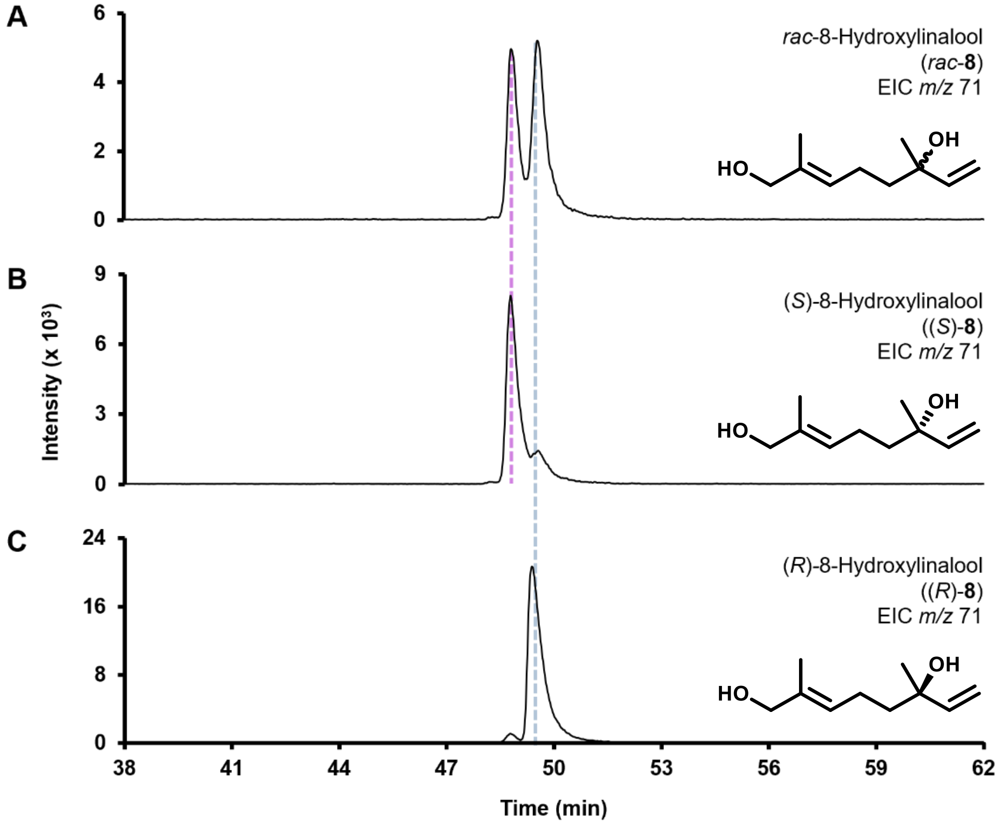

8-Oxolinalool (**7**)

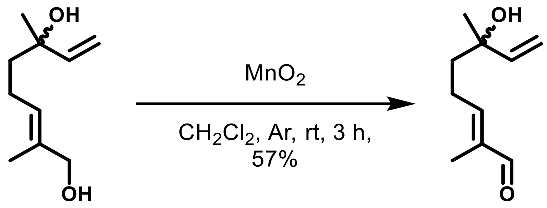


**8** **7**

To a dry 25 mL round bottom flask under an Ar atmosphere, containing diol **8** (386 mg, 2.27 mmol) in anhydrous $CH_2Cl_2$ (11.5 mL), was added $MnO_2$ (2.960 g, 34.05 mmol). After stirring in darkness at room temperature for 3 h, the resulting black suspension was eluted through a short plug of silica with $Et_2O$ (10 mL x 10). Careful concentration of the eluate under reduced pressure gave a colorless oil that was further purified by gradient flash column chromatography (0–50% $Et_2O$ in hexane) to afford title compound **7** as a colorless oil (218 mg, 57%): $R_f$ 0.28 ($Et_2O$/pentane, 1:1); $^1H$ NMR ($CDCl_3$, 400 MHz) $\delta$ 9.38 (d, $J$ = 1.8 Hz, 1H), 6.51–6.47 (m, 1H), 5.92 (ddd, $J$ = 17.3, 10.8, 1.2 Hz, 1H), 5.27–5.23 (m, 1H), 5.13–5.10 (m, 1H), 2.48–2.33 (m, 2H), 1.73 (s, 3H), 1.72–1.65 (m, 2H), 1.57 (brs, 1H), 1.33 (s, 3H); GC-MS (EI) $m/z$ (%) 71 (100), 87 (34), 82 (30), 55 (28), 83 (26), 98 (20), 97 (15), 95 (14), 67 (14), 84 (13). Physical and spectral data agreed with those reported previously [52].

Note: Following this procedure, (*S*)-**7** and (*R*)-**7** were analogously prepared from (*S*)- and (*R*)-**8**, respectively.

Chiral GC-EI-MS EIC chromatograms for (**A**) *rac*-8-oxolinalool (*rac*-**7**), (**B**) (*S*)-**7**, and (**C**) (*R*)-**7**. Separation method: F.

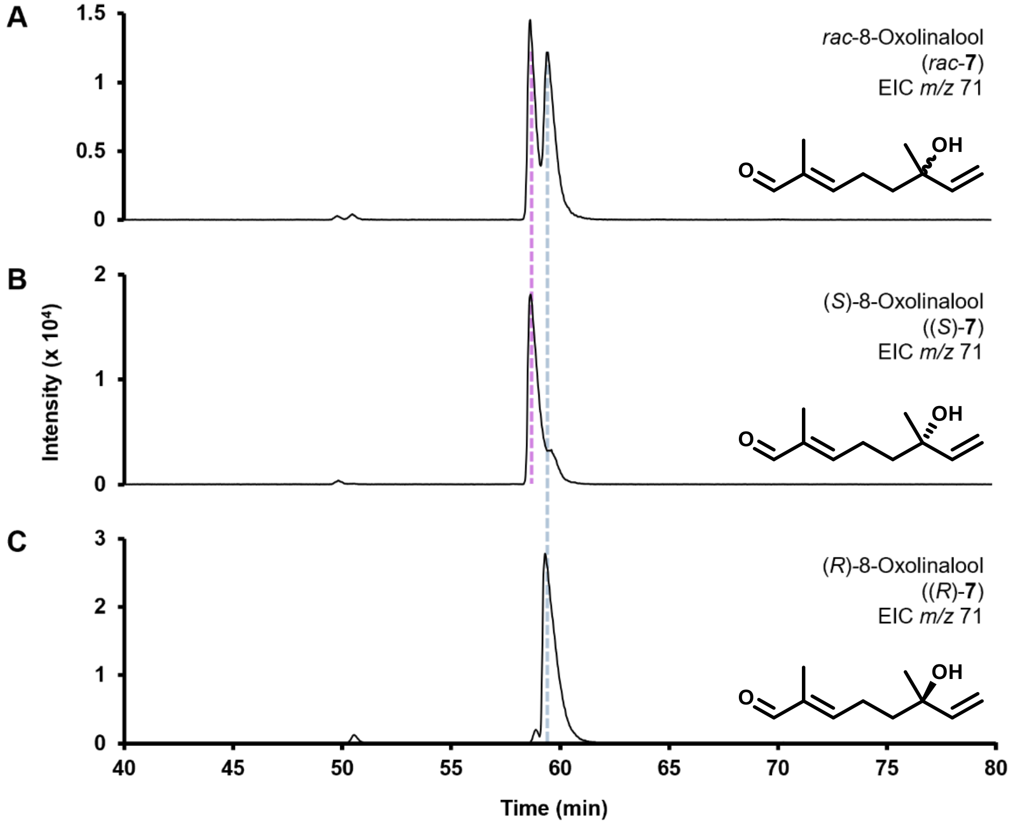

**Synthesis of lilac aldehyde (3) and lilac alcohol (6)**

Lilac alcohol **6** was prepared over two steps from **S4**, following Ohno's route [53].

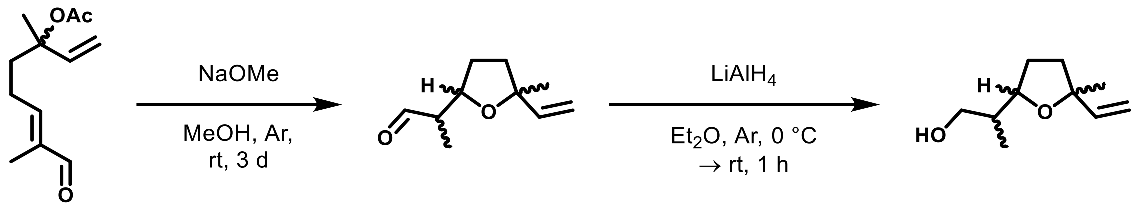


**S4** **3** **6**

Lilac aldehyde (**3**)

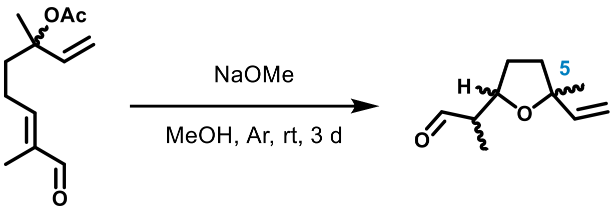


*rac*-**S4** → **3**, 15%
(3*S*)-**S4** → (5*S*)-**3**, 12%
(3*R*)-**S4** → (5*R*)-**3**, 18%

To a dry 250 mL round bottom flask under an Ar atmosphere, containing acetate *rac*-**S4** (3.233 g, 15.38 mmol) dissolved in anhydrous MeOH (154 mL), was added 5.4 M NaOMe in MeOH (1.14 mL, 6.15 mmol). After stirring in darkness at room temperature for 3 days, the resulting amber solution was adjusted to pH 7 using Dowex$^{TM}$ 50W X8 resin and filtered. The filtrate was dried over anhydrous $Na_2SO_4$ and carefully concentrated under reduced pressure to afford a residue that was then fractionally distilled *in vacuo* (b.p. 99–101 °C/31 mbar) to give a colorless oil that was further purified by gradient flash column chromatography (0–25% $Et_2O$ in petroleum ether [b.p. 40–60 °C]) to yield title compound **3** as a colorless oil† (388 mg, 15%): $R_f$ 0.23 and 0.20 ($Et_2O$/pentane, 1:9); $^1$H NMR ($CDCl_3$, 400 MHz) mixture of stereoisomers* $\delta$ 9.81 (d, $J$ = 2.5 Hz, 0.2H), 9.80 (d, $J$ = 1.3 Hz, 0.2H), 9.79 (d, $J$ = 2.4 Hz, 0.4H), 9.77 (d, $J$ = 1.6 Hz, 0.2H), 5.94–5.81 (m, 1H), 5.22–5.15 (m, 1H), 5.03–4.97 (m, 1H), 4.29–4.11 (m, 1H), 2.64–2.46 (m, 1H), 2.11–2.00 (m, 1H), 1.96–1.86 (m, 1H), 1.82–1.66 (m, 2H), 1.31 (d, $J$ = 3.0 Hz, 3H), 1.16 (d, $J$ = 7.0 Hz, 0.6H), 1.12 (d, $J$ = 7.0 Hz, 0.7H), 1.07 (d, $J$ = 7.0 Hz, 1H), 1.04 (d, $J$ = 7.0 Hz, 0.8H). Physical and spectral data agreed with those reported previously [53]. *All four diastereoisomers of **3** are described; †Lilac aldehyde **3** is highly volatile and also degrades upon standing at room temperature. Following isolation, **3** should be immediately stored at −20 °C under Ar.

Note: Following this procedure, (5*R*)-**3** and (5*S*)-**3** were analogously prepared from (*R*)- and (*S*)-**S4**, respectively.

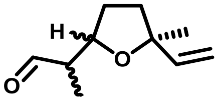


(5*S*)-**3**, 12%

(5*S*)-lilac aldehyde ((5*S*)-**3**) was isolated as a colorless oil (249 mg [12%] from 2.595 g [12.34 mmol] of (*S*)-**3**): GC-MS (EI) *m*/*z* (%) *1*) 55 (100), 93 (68), 111 (65), 67 (52), 71 (43), 69 (36), 110 (34), 81 (32), 68 (30), 153 (27), *2*) 55 (100), 93 (64), 71 (55), 111 (53), 67 (48), 69 (36), 81 (31), 68 (31), 153 (28), 110 (27), *7*) 55 (100), 111 (86), 93 (70), 67 (42), 69 (38), 71 (29), 81 (28), 110 (27), 153 (26), 68 (24), *8*) 55 (100), 93 (60), 111 (58), 71 (43), 67 (38), 69 (35), 153 (30), 81 (26), 68 (24), 110 (20). Physical and spectral data agreed with those reported previously [53].

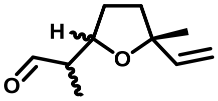


(5*R*)-**3**, 18%

(5*R*)-lilac aldehyde ((5*R*)-**3**) was isolated as a colorless oil (454 mg [18%] from 3.154 g [15 mmol] of (*R*)-**3**). Physical and spectral data agreed with those reported previously [53].

Chiral GC-EI-MS EIC chromatograms for (**A**) lilac aldehyde **3**, (**B**) (5*S*)-**3**, and (**C**) (5*R*)-**3**. Elution order of **3** stereoisomers: *1*) (2´*S*,2*S*,5*S*)-**3**, *2*) (2´*R*,2*S*,5*S*)-**3**, *3*) (2´*R*,2*R*,5*R*)-**3**, *4*) (2´*S*,2*R*,5*R*)-**3**, *5*) (2´*S*,2*S*,5*R*)-**3**, *6*) (2´*R*,2*S*,5*R*)-**3**, *7*) (2´*S*,2*R*,5*S*)-**3**, and *8*) (2´*R*,2*R*,5*S*)-**3** [54]. Separation method: C. *Mixture of stereoisomers.

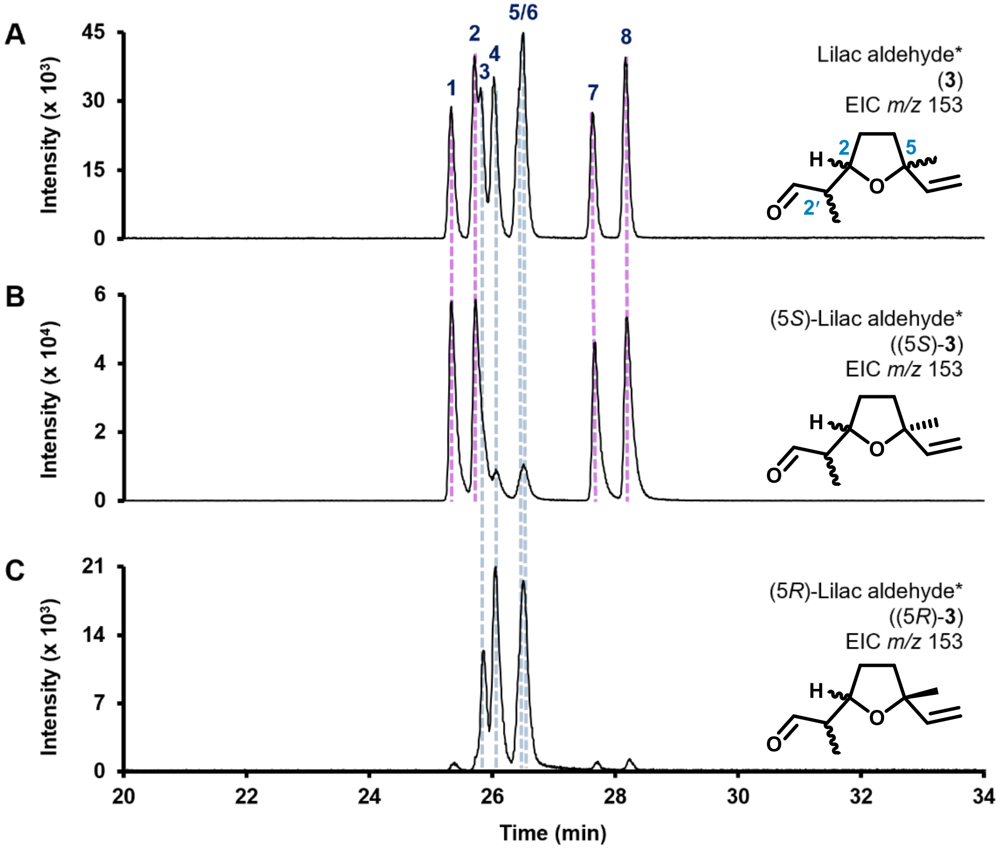

Lilac alcohol (**6**)

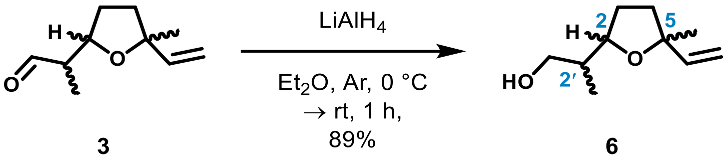


To a dry 8 mL vial under an Ar atmosphere, containing $LiAlH_4$ (54 mg, 1.42 mmol) suspended in anhydrous $Et_2O$ (0.5 mL) at 0 °C, was added dropwise a solution of aldehyde **3** (100 mg, 0.59 mmol) in anhydrous $Et_2O$ (0.6 mL). After warming to room temperature and stirring for 1 h the resulting suspension was cooled to 0 °C and cautiously treated stepwise with $H_2O$ (0.05 mL), 15% (w/v) aq. NaOH (0.05 mL), $H_2O$ (1.5 mL), and $Et_2O$ (2 mL). Upon stirring at room temperature for 30 min, the grey mixture was filtered through a short pad of neutral $Al_2O_3$ and the resulting filtrate carefully concentrated under reduced pressure to afford title compound **6** as a colorless oil (89 mg, 89%): $R_f$ 0.63 ($Et_2O$); $^{1}H$ NMR ($CDCl_3$, 400 MHz) mixture of stereoisomers* $\delta$ 5.97–5.81 (m, 1H), 5.23–5.14 (m, 1H), 5.01–4.97 (m, 1H), 4.16–4.10 (m, 0.4H), 3.86–3.77 (m, 0.6H), 3.72–3.55 (m, 2H), 2.07–1.94 (m, 1H), 1.93–1.55 (m, 5H), 1.31 (s, 3H), 0.94 (dd, $J$ = 7.0 Hz, 0.3H), 0.90 (dd, $J$ = 7.0 Hz, 0.9H), 0.80 (dd, $J$ = 6.8 Hz, 0.8H), 0.78 (dd, $J$ = 6.5 Hz, 1H). Physical and spectral data agreed with those reported previously.[53] *All four diastereoisomers of **6** are described.

Note: Following this procedure, (5*R*)-**6** and (5*S*)-**6** were analogously prepared from (5*R*)- and (5*S*)-**3**, respectively.

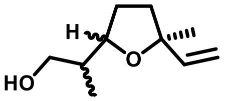

(5*S*)-**6**, 87%

(5*S*)-lilac alcohol ((5*S*)-**6**) was isolated as a colorless oil (132 mg [87%] from 150 mg [0.89 mmol] of (5*S*)-**3**): GC-MS (EI) *m*/*z* (%) *1*) 111 (100), 93 (77), 55 (60), 67 (33), 71 (25), 81 (21), 69 (19), 155 (18), 83 (12), 91 (11), *4*) 111 (100), 93 (78), 55 (63), 67 (30), 71 (24), 81 (20), 155 (19), 69 (19), 83 (12), 91 (11), *5*) 111 (100), 93 (79), 55 (62), 67 (37), 71 (22), 81 (22), 69 (19), 155 (16), 83 (13), 91 (11), *8*) 111 (100), 93 (82), 55 (70), 67 (35), 71 (27), 81 (24), 155 (22), 69 (19), 59 (18), 83 (13). Physical and spectral data agreed with those reported previously [53].

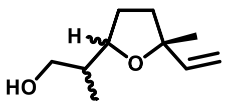

(5*R*)-**6**, 79%

(5*R*)-lilac alcohol ((5*R*)-**6**) was isolated as a colorless oil (120 mg [79%] from 150 mg [0.89 mmol] of (5*R*)-**3**). Physical and spectral data agreed with those reported previously [53].

Chiral GC-EI-MS EIC chromatograms for (**A**) lilac alcohol **6**, (**B**) (5*S*)-**6**, and (**C**) (5*R*)-**6**. Elution order of **6** stereoisomers: *1*) (2´*S*,2*S*,5*S*)-**6**, *2*) (2´*R*,2*R*,5*R*)-**6**, *3*) (2´*R*,2*S*,5*R*)-**6**, *4*) (2´*S*,2*R*,5*S*)-**6**, *5*) (2´*R*,2*S*,5*S*)-**6**, *6*) (2´*S*,2*R*,5*R*)-**6**, *7*) (2´*S*,2*S*,5*R*)-**6**, and *8*) (2´*R*,2*R*,5*S*)-**6** [54]. Separation method: C. *Mixture of stereoisomers.

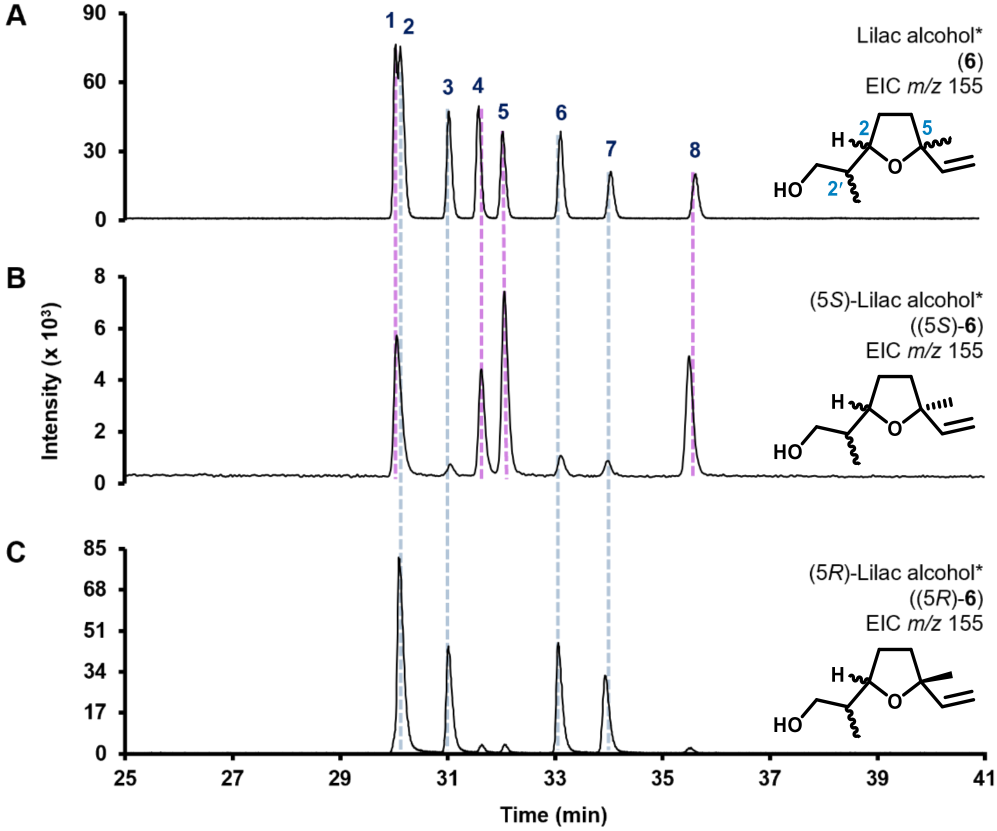

**NMR Data**

$^{1}H$ NMR ($CDCl_3$, 400 MHz) **5**

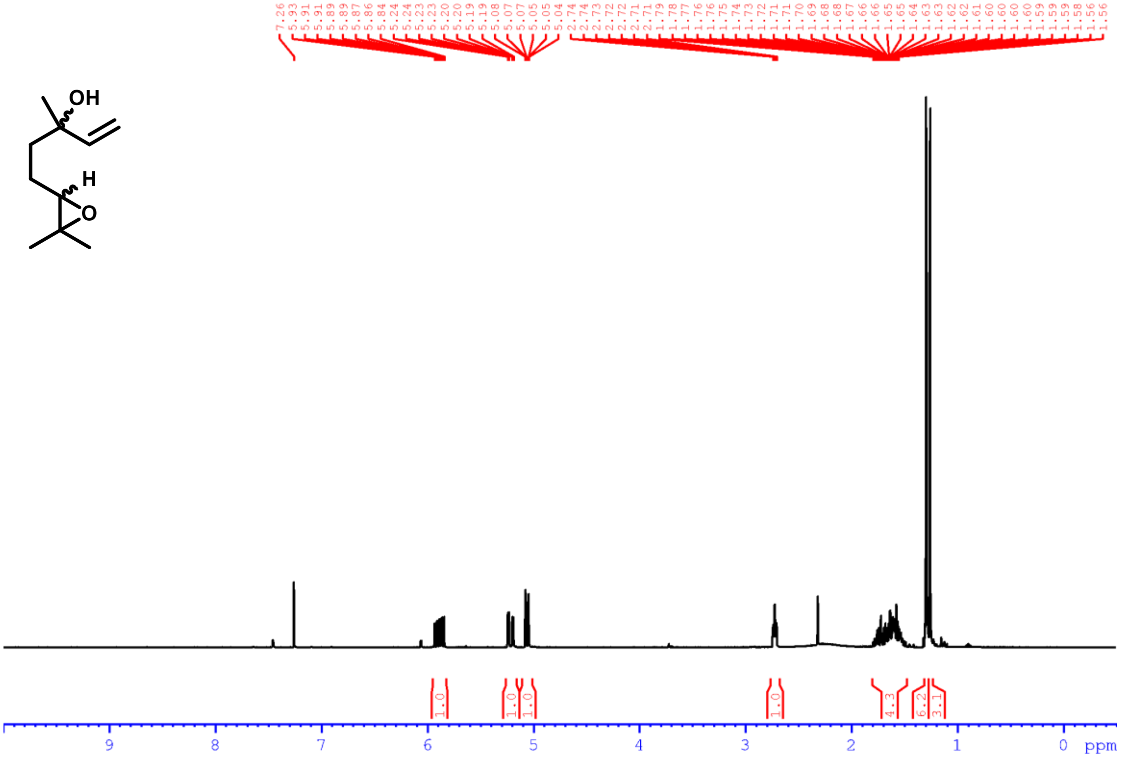

$^{1}H$ NMR ($CDCl_3$, 400 MHz) (5*S*)-**1**

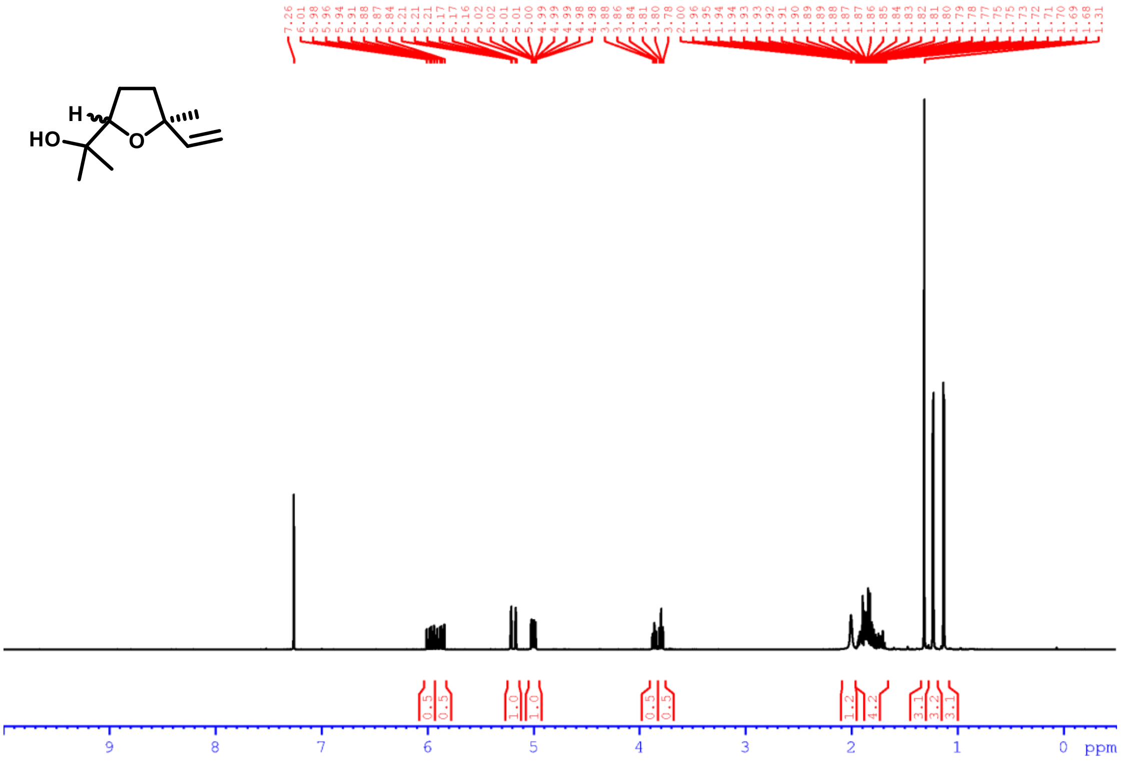

$^{1}$H NMR ($CDCl_3$, 400 MHz) (6*S*)-**4**

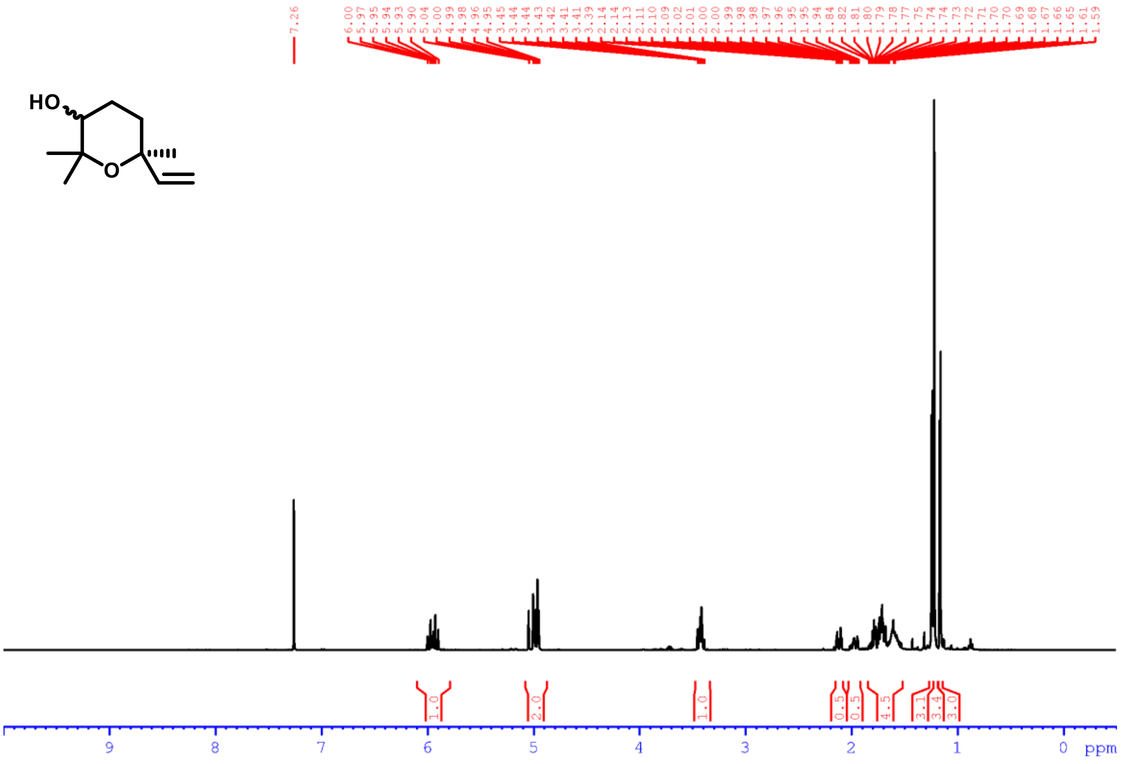


$^{1}$H NMR ($CDCl_3$, 400 MHz) *rac*-**S3**

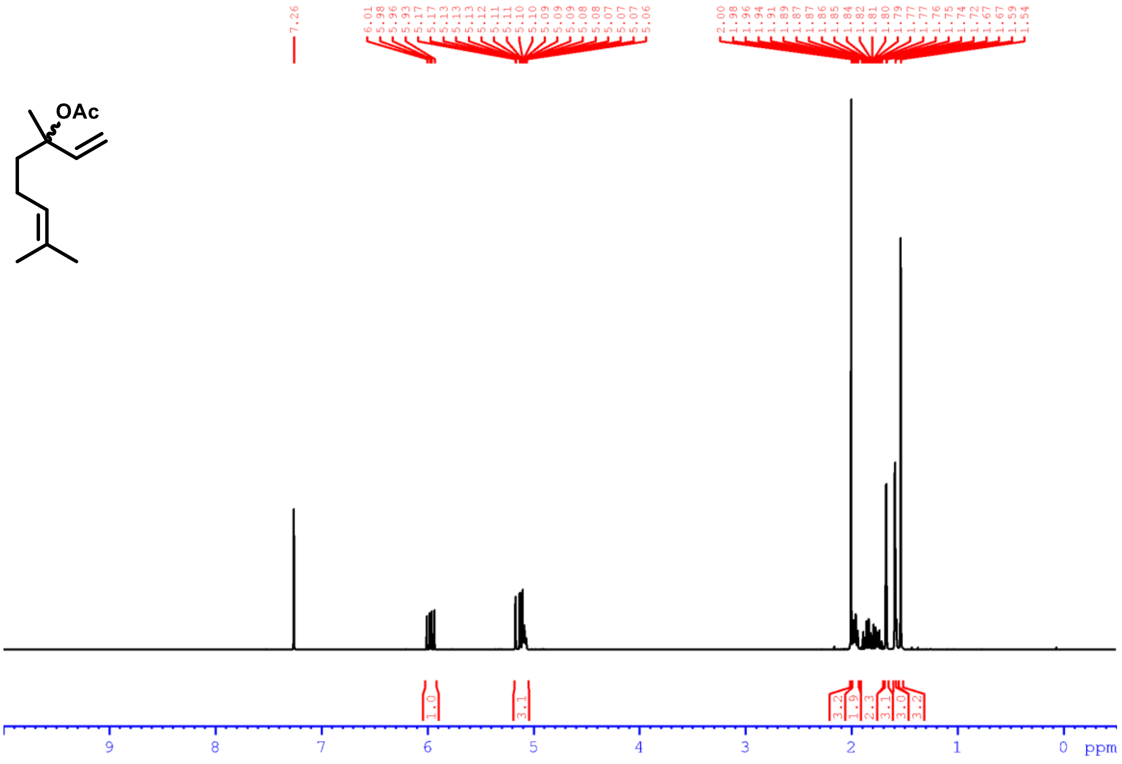

$^1$H NMR ($CDCl_3$, 400 MHz) *rac*-**S4**

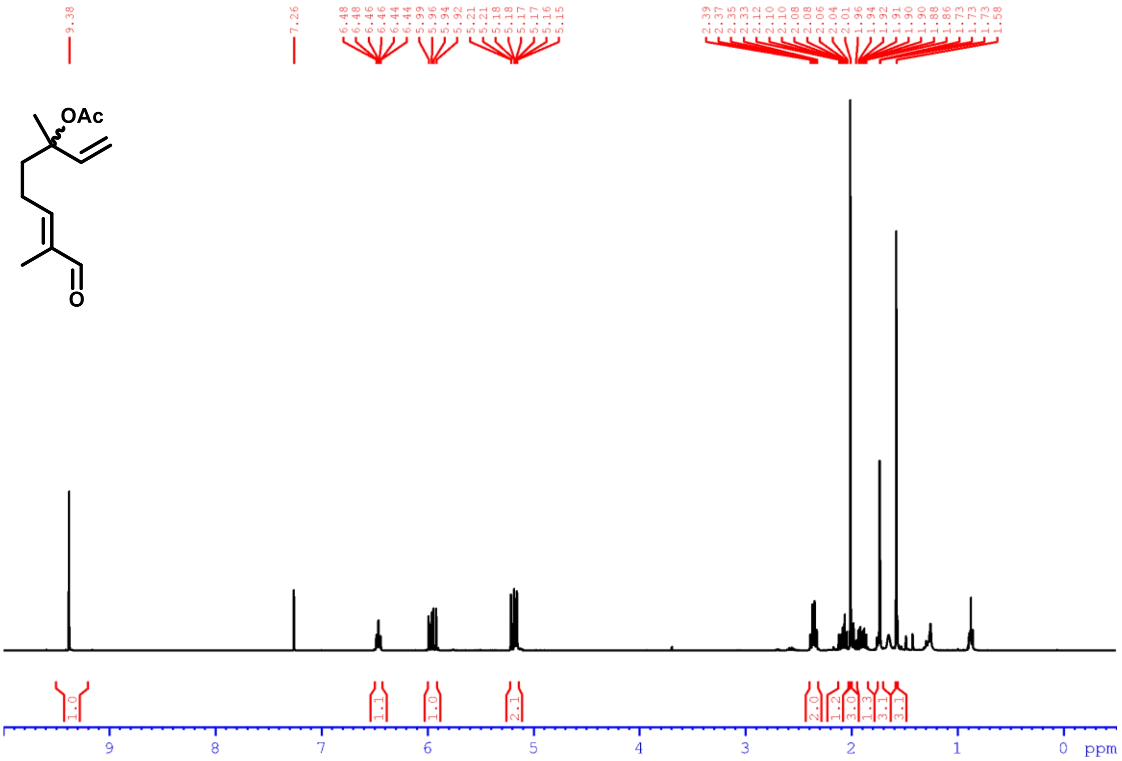


$^1$H NMR ($CDCl_3$, 400 MHz) *rac*-**8**

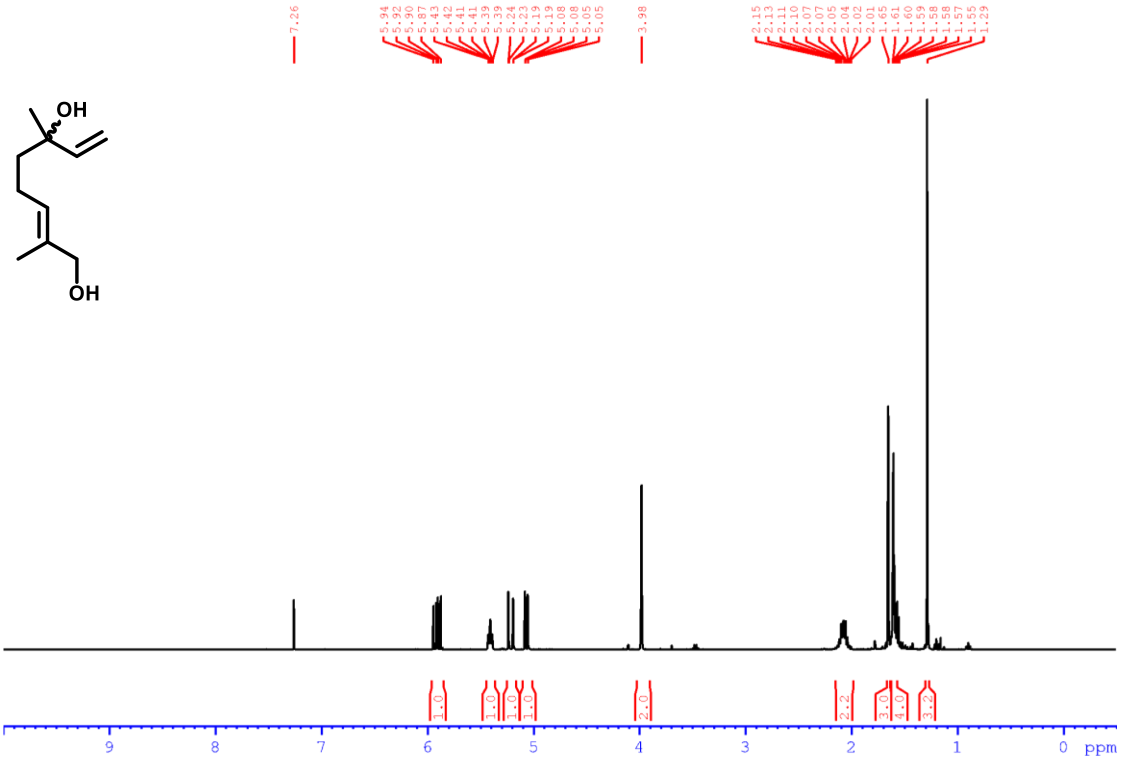

$^{1}$H NMR ($CDCl_3$, 400 MHz) *rac*-**7**

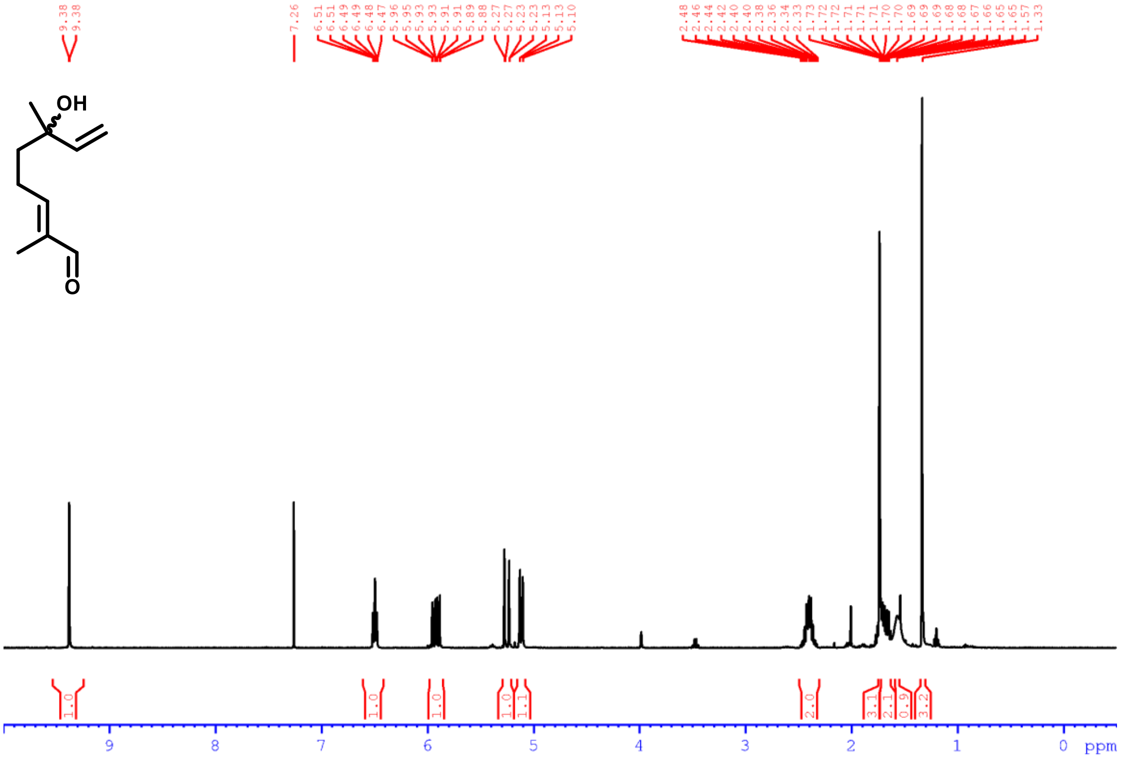


$^{1}$H NMR ($CDCl_3$, 400 MHz) **3**

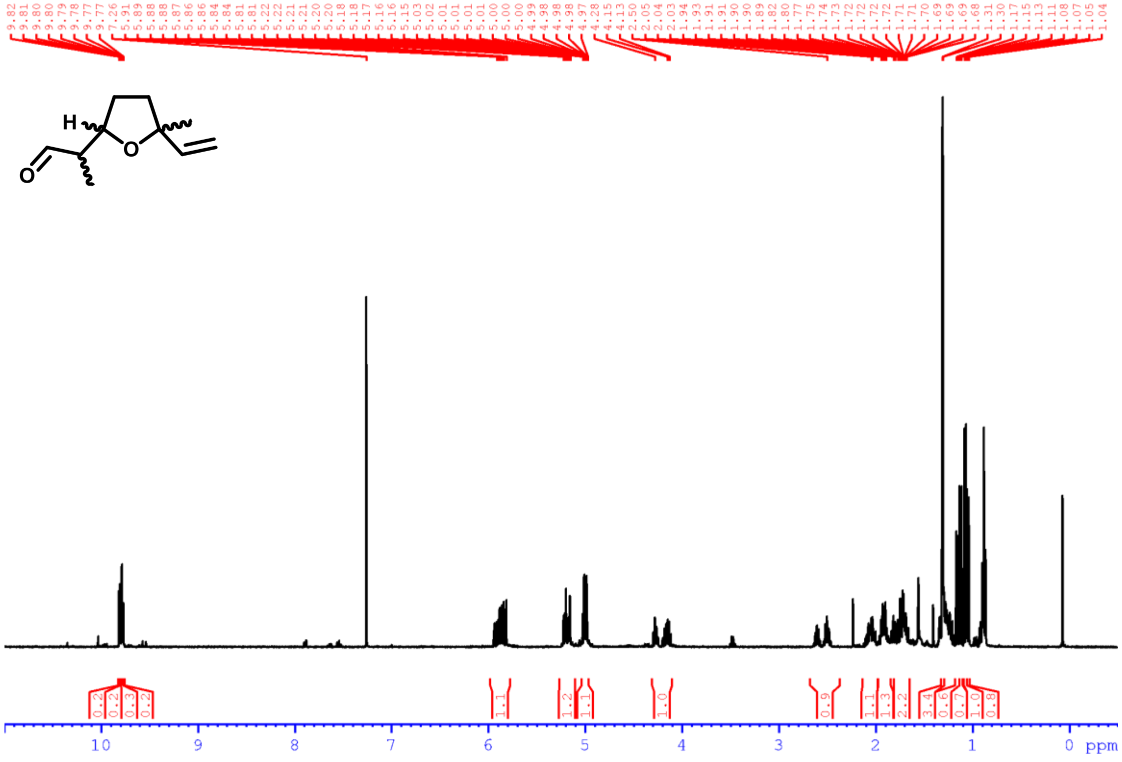

[1]H NMR ($CDCl_3$, 400 MHz) **6**

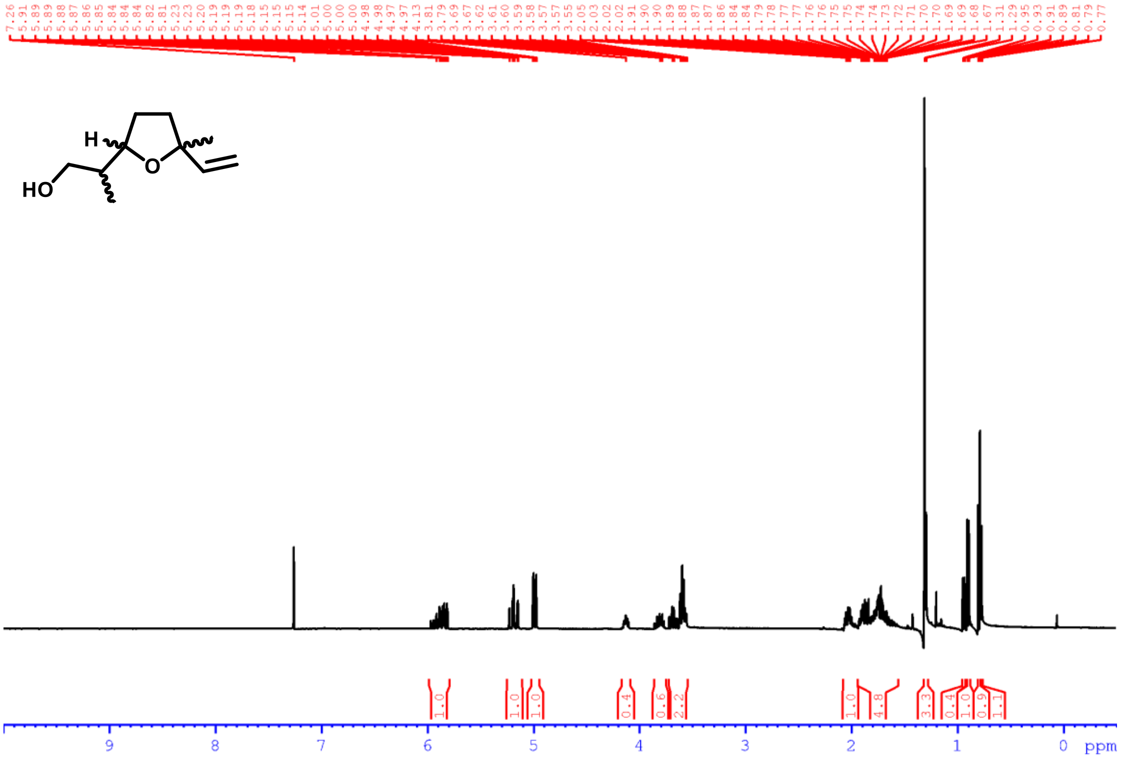

**Bee acquisition and maintenance**

*Osmia bicornis* were commercially obtained as individual pupae (Sven Behr Hummelvertrieb; Welle, Lower Saxony, Germany) and *Bombus terrestris* were acquired as a colony (Katz Biotech AG; Baruth/Mark, Brandenburg, Germany). Specimens of *Colletes similis* were collected in the field (Jena, Thuringia, Germany) with permission from Untere Naturschutzbehörde Jena and *Apis mellifera* were from a local beehive at the Max Planck Institute for Chemical Ecology (MPI-CE; Jena, Thuringia, Germany). Wild-caught bees were stored at 4 °C overnight prior to use in experiments. Animals were individually starved for 4 h in plastic vials containing a bed of moist tissue paper, directly before use in behavioral assays.

**Y-tube olfactometer dual-choice assays with bees**

A Y-tube olfactometer (see **Supplementary Fig. 21**) was used in dual-choice assays to assess the behavioral responses of all tested bee species (*O. bicornis*, *C. similis*, *B. terrestris*, and *A. mellifera*) to different odor stimuli. The apparatus consisted of a 3D-printed Y-tube (main arm: length, 12 cm; side-arms: length, 12 cm; internal diameter: 2.2 cm) connected *via* silicon tubing to two 250 mL glass bottles (bottles A1 and A2 and bottles B1 and B2) at the end of each side-arm. When testing bee preference for extracted triungula volatiles obtained by DHV collection, synthetic compounds, or blends thereof, $CH_2Cl_2$ was used as solvent. Treatment samples were filled into 2 mL glass vials and then placed in bottle A1 (see **Supplementary Fig. 21B**). To test extracts and synthetic compounds against more than just neat $CH_2Cl_2$ or nothing, *i.e.*, not just "scent versus *no* scent", a block of eight freshly sprouted wheat plants (*Triticum aestivum*) were placed in bottle A1 of each side-arm to create a standardized volatile background. Vials containing solutions of synthetic compounds, DHV collection extracts, or neat $CH_2Cl_2$ were placed in a holder that positioned it in the center of the glass bottle above the wheat. For choice assays with living triungula, animals were placed directly on the wheat in bottle A1. To maintain humidity in the testing arena and thus preserve volatile perception, in-house purified compressed air was first passed through bottle A2 or B2 which were filled with distilled water, prior to entering bottles A1 or B1 (containing wheat and a solution of synthetic compound in $CH_2Cl_2$ or neat $CH_2Cl_2$), respectively. This charcoal-filtered and humidified air was pumped through each side-arm of the Y-tube at a constant flow rate of 150 mL/min and was regulated by flow meters to ensure equal airflow. At the base of the main arm, air was extracted from the system at a rate of 420 mL/min, to prevent saturation of volatiles in the Y-tube and create a continuous air flow through both side-arms. Before each trial, the Y-tube was cleaned with 70% (v/v) EtOH and oven-dried at 60 °C to remove residual odors. The experiment was conducted under controlled environmental conditions (23 ± 1 °C, 65 ± 5% RH). A Planon Plus 30 x 60 cm light panel (22 W; 4000 K; 1600 lm) was used as a light source over the enclosed Y-tube to create a stable light intensity of 100 µmol/($m^2$s) on the surface of the arena. An individual insect was gently released at the base of the main arm and allowed up to 5 min to make a choice by walking to the end of one of the two side-arms. Individuals that did not make a choice within the allotted time were recorded as "no response" and excluded from statistical analyses. To avoid positional bias, the odor sources were alternated between the left and right arms after every insect, and the Y-tube apparatus was replaced or cleaned regularly. Between two test subjects and the exchange of the odor sources, air was allowed to flow through the setup for 5 min. During experiments, the arena was monitored from above using a webcam that transmitted the image to a laptop so as not to influence the animals' decision. Each treatment was tested with 18–22 replicates, using a new bee for each repetition. The proportion of bees that chose each side-arm was analyzed using a chi-square ($\chi^2$) test of independence (Microsoft® Excel® LTSC MSO 16.0) with a significance level of $p < 0.05$.

**Triungulin olfactometric choice assays**

An in-house setup, consisting of two equal-sized wooden toothpicks fixed to a sheet of circular filter paper on an inverted glass Petri dish positioned within a larger Petri dish containing water (see **Supplementary Fig. 25**) was used to assess triungulin preference for a synthetic blend of (*S*)-linalool-derived monoterpenoids. To remove light bias, all experiments were performed in a dark room under an LED light panel. Immediately before starting an experiment, one toothpick was dosed with 50 µL of the synthetic (*S*)-linalool-derived monoterpenoid blend dissolved in $CH_2Cl_2$ and 50 µL of neat $CH_2Cl_2$ was applied to the other. Approximately 20 triungula were directly released mid-way between these two toothpicks ($t_0$). The absolute number of triungula present on each toothpick was then recorded at 5 min intervals for 30 min, following release. Absolute numbers of triungula on toothpicks at each time point were recorded and analyzed using a paired t-test: paired two sample for means (Microsoft® Excel® LTSC MSO 16.0) to determine if there was a statistically significant preference for the monoterpenoid blend (see **Supplementary Table 1**).

**Transcriptome sequencing and gene identification**

Female *M. proscarabaeus* beetles (biological replicates, *n* = 3) were dissected in PBS into fat body, gut, reproductive organs, and head, and immediately transferred to ice-cold Trizol (Direct-zol, Zymo Research). Samples were homogenized in ceramic bead tubes using a TissueLyser LT (Qiagen) and total RNA was extracted from the different tissues using the Direct-zol RNA Miniprep kit (Zymo Research) according to the manufacturer's instructions. The concentration and integrity of the RNA were determined using an N60 nanophotometer (Implen) and a TapeStation System (Agilent), respectively. After RNA-seq library preparation (poly-A enrichment), short-read sequencing was performed on an Illumina HiSeq 2500 sequencer (Max Planck Genome Centre, Cologne, Germany) with ~30 Mio reads per library, 150 bp per read, paired end. In addition, RNA from the different tissues were pooled and subjected to PacBio long-read RNA sequencing on the Sequel II system (Max Planck Genome Centre), resulting in ~93 Gb sequence data. Total RNA from *M. proscarabaeus* larvae (3 samples, each containing approximately 50 larvae) was extracted using the RNeasy® Plant Mini Kit (Qiagen) according to the manufacturer's instructions, and was sequenced by using Illumina technology (poly-A enrichment, NovaSeq X Plus (Novogene, Munich, Germany), 150 bp per read, paired end, 60 Mio raw reads per sample). Aliquots of the individual larval RNA extractions were pooled and PacBio long-read RNA sequencing was performed on the Revio system (Max Planck Genome Centre), resulting in ~950 Gb sequence data. For generating a reference transcriptome as mapping template, the PacBio sequence reads from female beetles and larvae were combined and processed using the IsoSeq pipeline including the removal of cDNA primer, demultiplexing, polyA-tail trimming and clustering. Trimming of the Illumina reads and mapping to the PacBio reference transcriptome were performed with the software CLC Genomics Workbench (Qiagen Bioinformatics, mapping parameters: mismatch cost, 2; insertion cost, 3; deletion cost, 3; length fraction, 0.8; similarity fraction, 0.95; maximum number of hits, 5). Note, the reads from the four different tissues of *M. proscarabaeus* female beetles were combined prior mapping.

To identify cytochrome P450 genes potentially involved in volatile terpenoid biosynthesis, we first selected genes annotated as CYP450 or CYP450-like. We further refined this list by selecting genes expressed at least tenfold more in the larvae than in female beetles, with a minimum expression threshold of 10 RPKM. This approach identified 15 potential candidate genes.

**Gene cloning**

Complementary DNA (cDNA) was prepared from total RNA of *M. proscarabaeus* triungula, using SuperScript™ IV VILO™ Master Mix kit (Thermo Fisher Scientific) following manufacturer's instructions. Coding sequences were amplified with Platinum™ SuperFi II DNA polymerase kit (Thermo Fisher Scientific) using *M. proscarabaeus* triungulin cDNA as template and gene-specific primers listed in **Supplementary Table 2**. Amplified sequences are described in **Supplementary Table 3**. Constructs for heterologous expression in *S. cerevisiae* were prepared by cloning CYP345BZ1 or CYP347BT1 in combination with MpRed as sticky-end fragments (MpRed) or using In-Fusion® (CYP345BZ1 and CYP347BT1; TaKaRa) into the same pESC-Leu-2d vector employing two different cloning sites with [55]. Inserted sequences were confirmed by Sanger sequencing.

**Heterologous expression of CYP450 gene candidates in *Saccharomyces cerevisiae* and *in vivo* enzyme assays**

Following a previously described procedure for heterologous expression in yeast [56], constructs were transformed into the INVSc1 (Thermo Fisher Scientific) *S. cerevisiae* strain using the *S.c.* EasyComp™ Transformation Kit (Invitrogen) according to the manufacturer's instructions. Subsequently, 30 mL Yeast Synthetic Drop-out Medium Supplement without leucine (Sigma Aldrich Y1376-20g; containing 70 mg/L adenine and 20 g/L D-glucose) was inoculated with single yeast colonies and grown overnight at 30 °C and 200 rpm. For main cultures, 100 mL yeast peptone glucose agar (YPGA) (Glc) full medium (10 g/L yeast extract, 20 g/L bactopeptone, 74 mg/L adenine hemisulfate, 20 g/L D-glucose) was inoculated with one-unit OD600 of the overnight cultures and incubated under the same conditions for 30 to 35 h. After centrifugation (5,000 × g, 16 °C, 5 min), expression was induced by resuspension of the cells in 100 mL YPGA (Gal) medium (see above, but including 20 g/L D-galactose instead of D-glucose) and grown for another 2 h at 30 °C and 200 rpm. For *in vivo* enzyme assays, a 600 µL aliquot of culture was transferred to a septum-sealed 4 mL glass vial and then treated with 6 µL of a 1 mg/mL solution of (*S*)-linalool in MeOH. After approximately 24 h at 30 °C and 200 rpm, the culture was overlaid with EtOAc (200 µL), and vortexed for 1 min. Following phase separation, the organic layer was carefully removed and submitted for achiral GC-EI-MS analysis using separation method A.

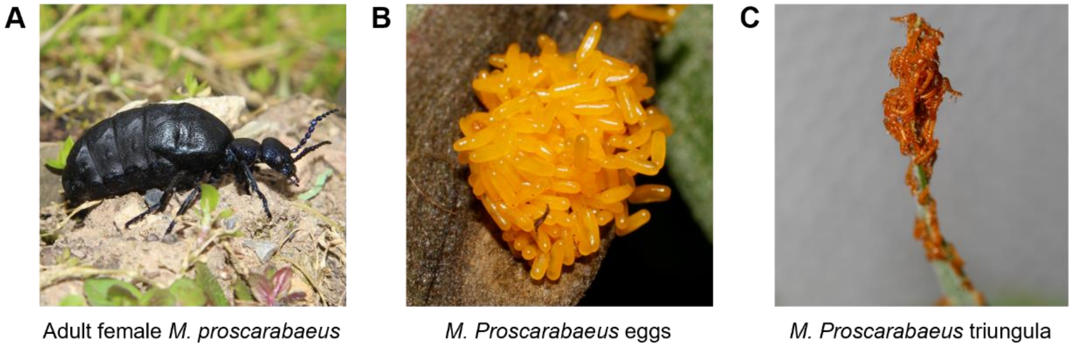


**Supplementary Fig. 1. Collection and cultivation of *M. proscarabaeus* triungula.**

(**A**) Wild adult female *M. proscarabaeus* collected in the field (Feb–Apr, 2024–25, Jena, Thuringia, Germany); (**B**) *M. proscarabaeus* eggs; (**C**) newly hatched *M. proscarabaeus* triungula aggregating on wheatgrass.

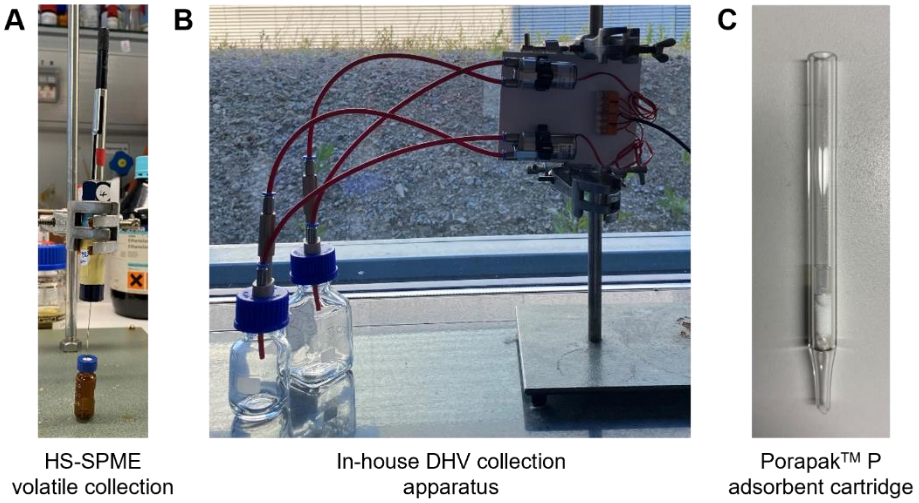


**Supplementary Fig. 2. Triungulin volatile collection.**

(**A**) Triungulin volatiles were collected using HS-SPME or (**B**) with an in-house dynamic headspace volatile (DHV) collection sampling apparatus fitted with a (**C**) removable Porapak$^{TM}$ P adsorbent cartridge.

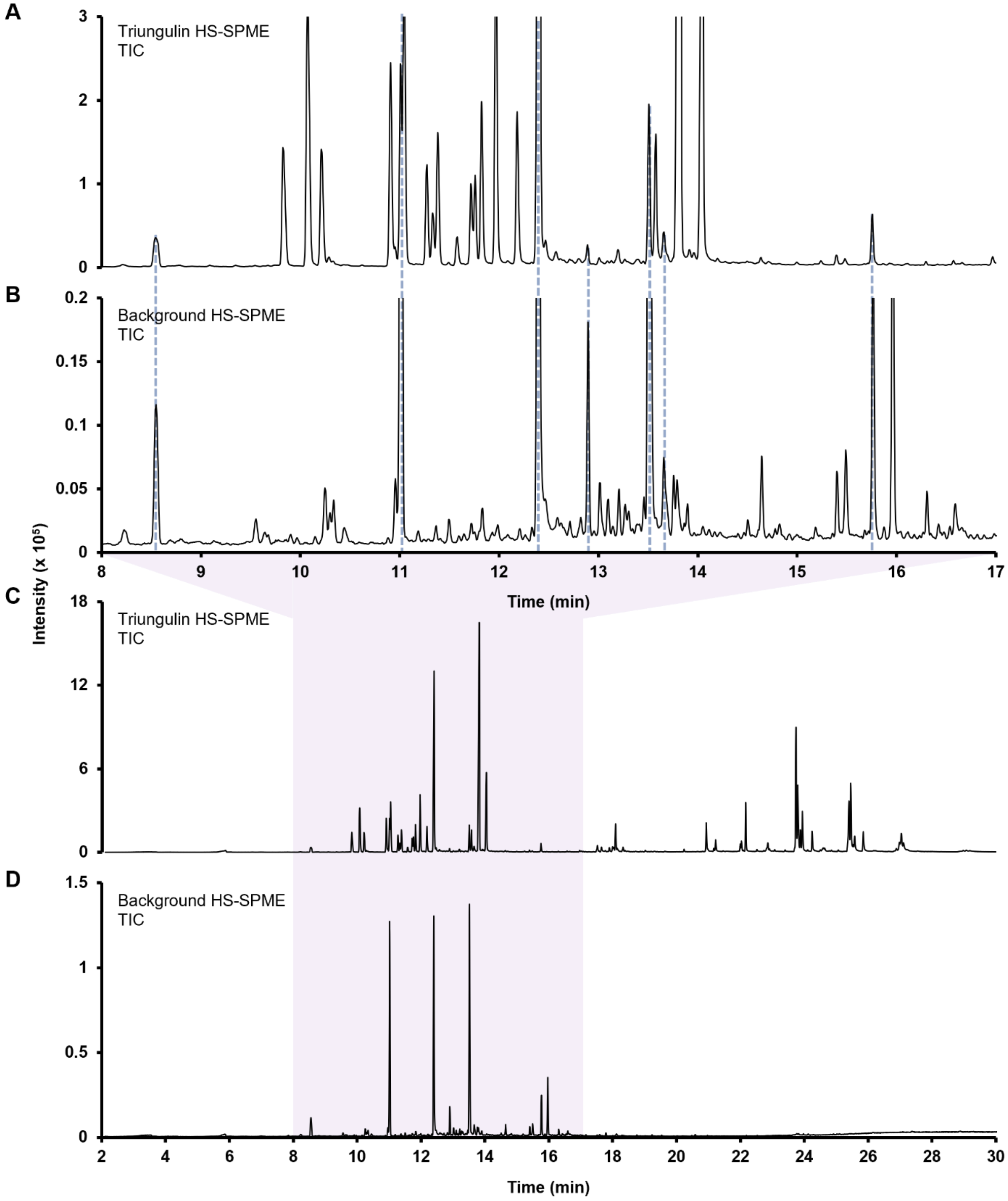


**Supplementary Fig. 3. Comparison of HS-SPME GC-EI-MS volatile profile of triungula with background.**

(**A**) Expansion of GC-EI-MS total ion current (TIC) chromatogram of triungulin volatile profile. (**B**) Expanded TIC chromatogram obtained for background. (**C**) Complete TIC chromatogram for triungula showing over thirty distinct $C_{19}$–$C_{27}$ saturated and unsaturated long-chain hydrocarbons that were tentatively identified using the National Institute of Standards and Technology (NIST) MS-Library v. 3.0 (2023). (**D**) Full TIC chromatogram for background. Separation method: A.

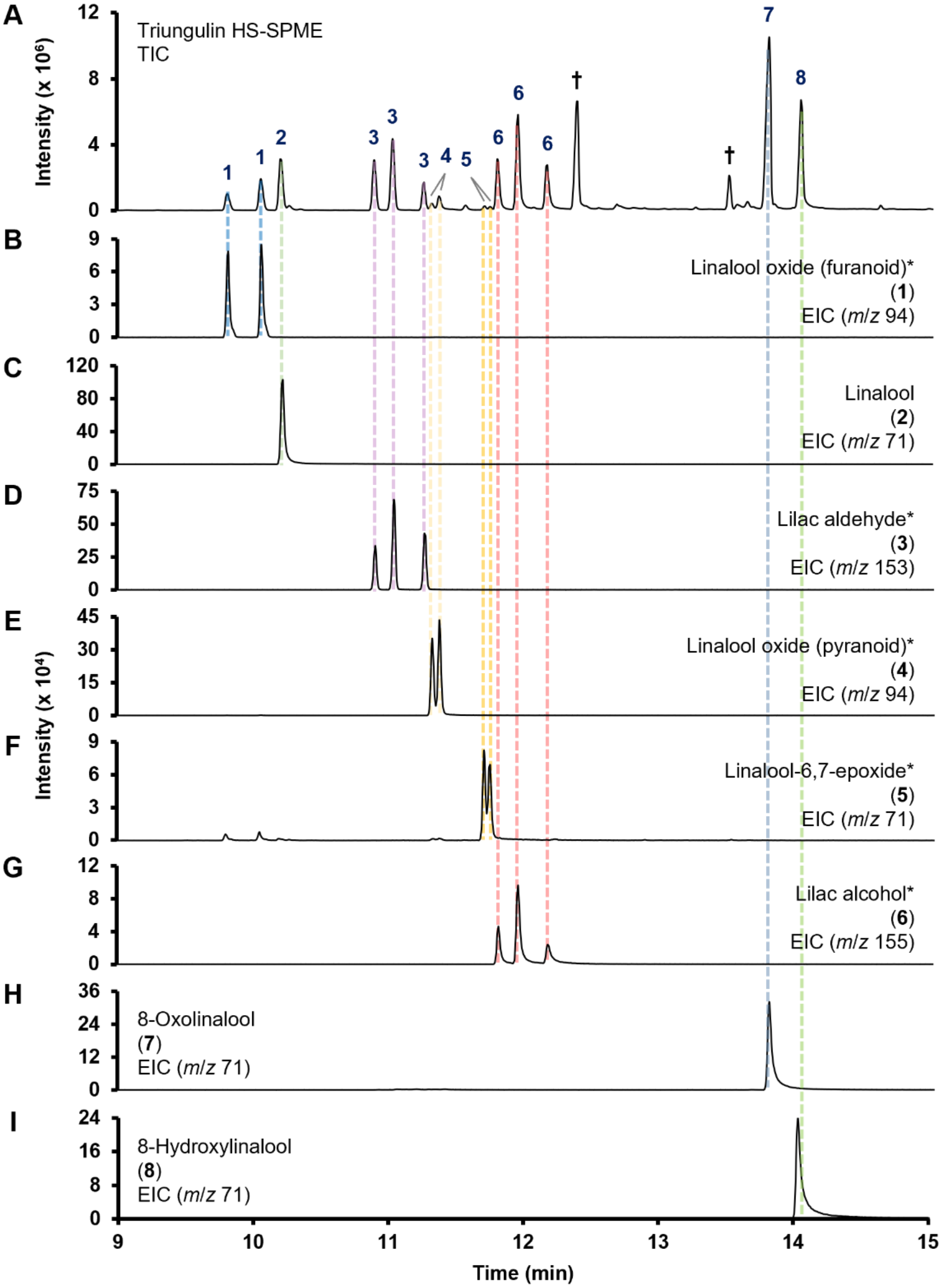


**Supplementary Fig. 4. Identification of triungulin monoterpenoid volatiles.**
Comparison of achiral GC-EI-MS chromatographic data obtained for (**A**) HS-SPME of triungula, (**B**) linalool oxide (furanoid) (**1**)*, (**C**) linalool (**2**), (**D**) lilac aldehyde (**3**)*, (**E**) linalool oxide (pyranoid) (**4**)*, (**F**) linalool-6,7-epoxide (**5**)*, (**G**) lilac alcohol (**6**)*, (**H**) 8-oxolinalool (**7**), and (**I**) 8-hydroxylinalool (**8**). Separation method: A. *Mixture of stereoisomers. †Background.

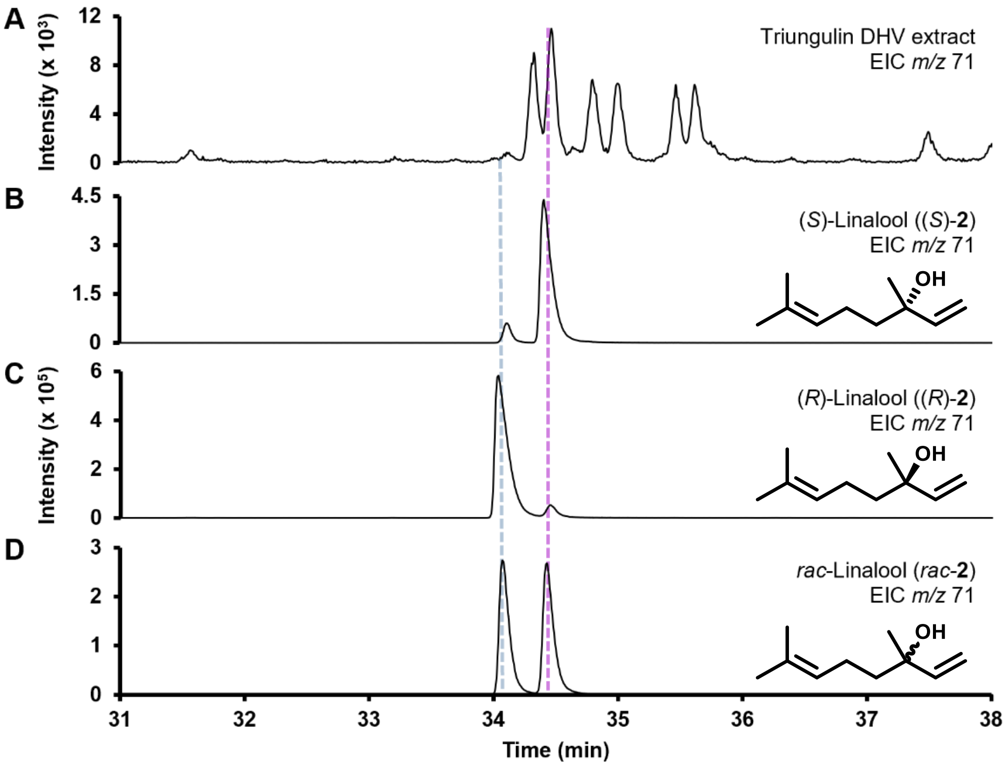


**Supplementary Fig. 5. Triungula exclusively emit (*S*)-linalool ((*S*)-2).**
Comparison of chiral GC-EI-MS chromatographic data obtained for (**A**) triungula DHV collection extract, (**B**) (*S*)-**2**, (**C**) (*R*)-**2**, and (**D**) *rac*-**2**. Separation method: B.

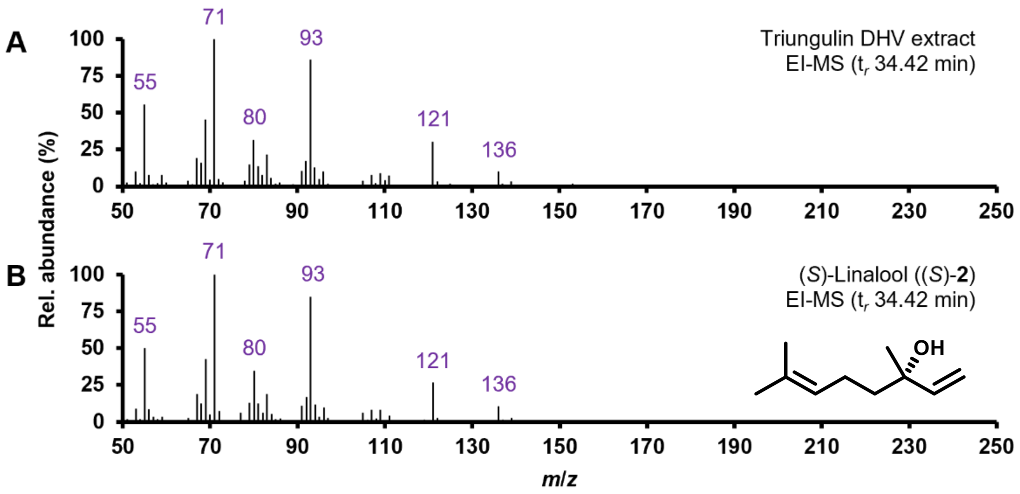


**Supplementary Fig. 6. Nominal EI-MS spectra confirm peak identity as (*S*)-linalool ((*S*)-2).** Comparison of nominal EI-MS data obtained for (**A**) triungulin DHV collection extract peak ($t_r$ 34.42 min, **Supplementary Fig. 5**) and (**B**) (*S*)-**2** reference standard.

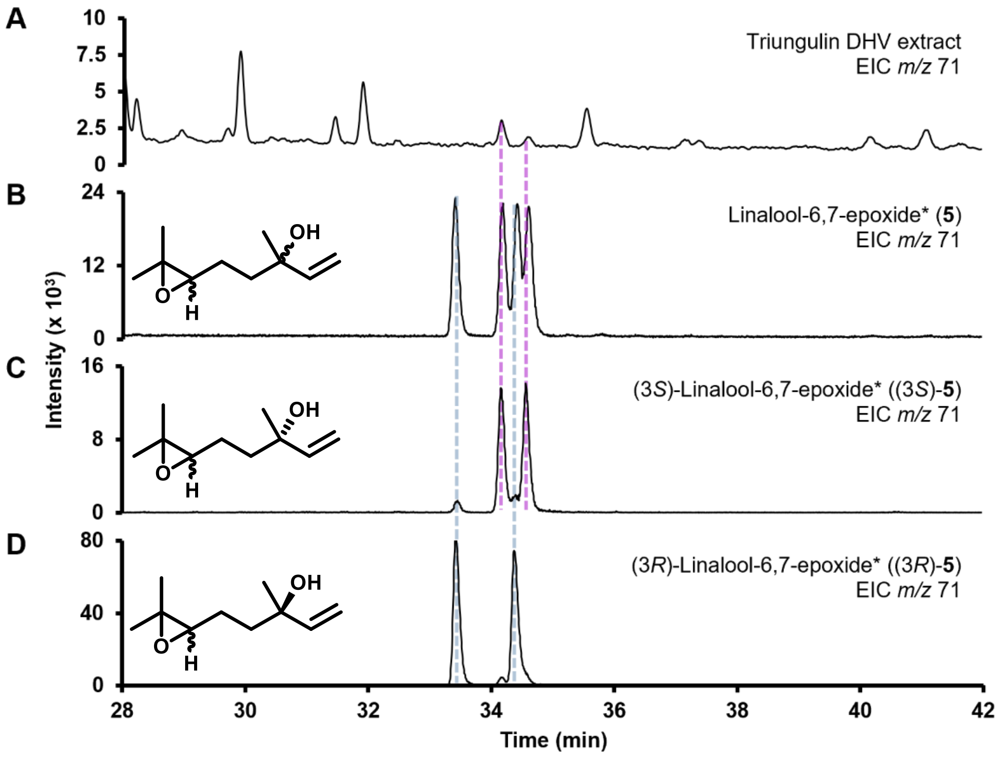


**Supplementary Fig. 7. Triungula emit both stereoisomers of (3*S*)-linalool-6,7-epoxide.** Comparison of chiral GC-EI-MS chromatographic data obtained for (**A**) triungula DHV collection extract, (**B**) linalool-6,7-epoxide (**5**), (**C**) (3*S*)-**5**, and (**D**) (3*R*)-**5**. Separation method: C. *Mixture of stereoisomers.

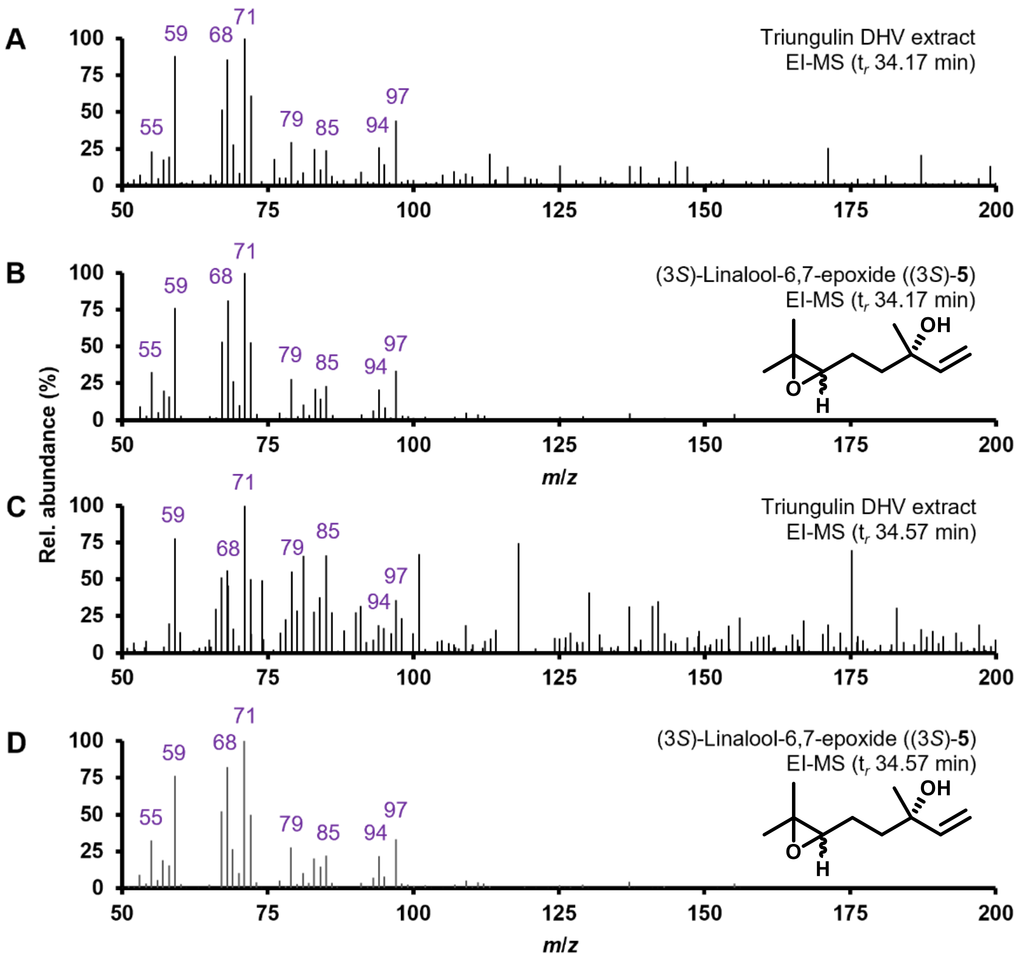


**Supplementary Fig. 8. Nominal EI-MS spectra confirm peak identities as both stereoisomers of (3*S*)-linalool-6,7-epoxide.**

Comparison of nominal EI-MS data obtained for (**A**) triungulin DHV collection extract peak ($t_r$ 34.17 min, **Supplementary Fig. 7**), (**B**) (3*S*)-**5** reference standard ($t_r$ 34.17 min, **Supplementary Fig. 7**), (**C**) triungulin DHV collection extract peak ($t_r$ 34.57 min, **Supplementary Fig. 7**), and (**D**) (3*S*)-**5** reference standard ($t_r$ 34.57 min, **Supplementary Fig. 7**).

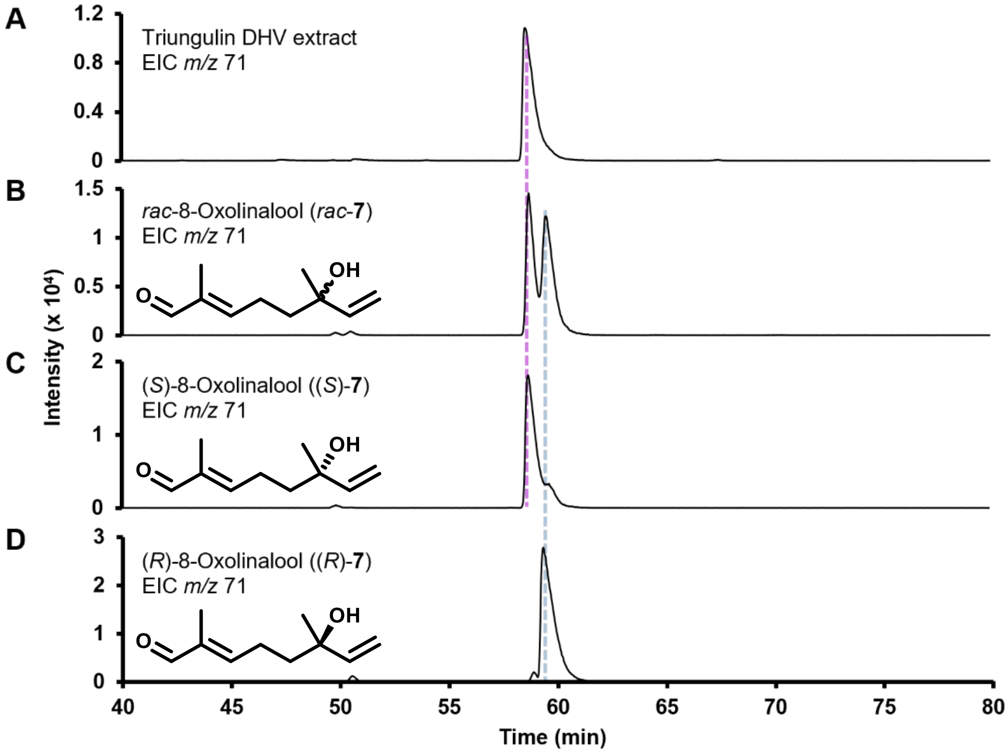


**Supplementary Fig. 9. Triungula emit (*S*)-8-oxolinalool ((*S*)-7).**

Comparison of chiral GC-EI-MS chromatographic data obtained for (**A**) triungula DHV collection extract, (**B**) *rac*-8-oxolinalool (*rac*-7), (**C**) (*S*)-**7**, and (**D**) (*R*)-**7**. Separation method: F.

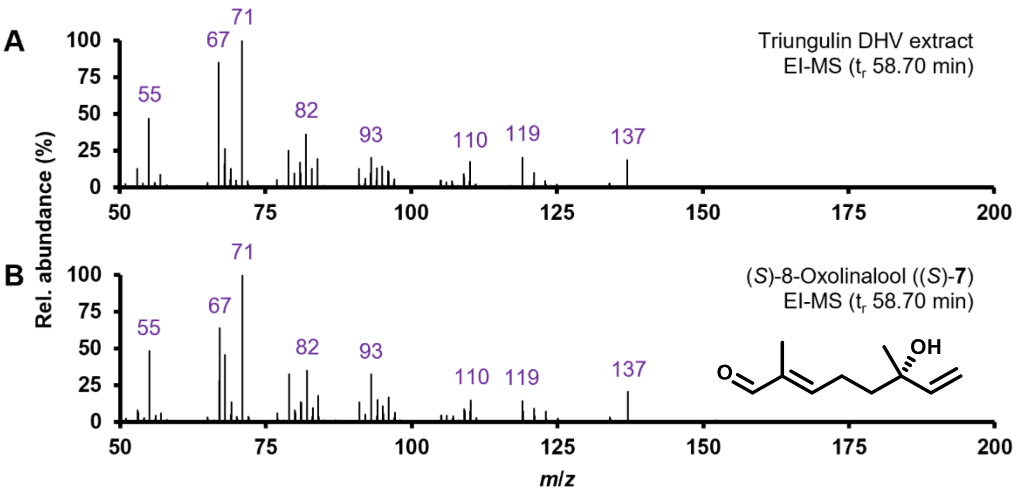


**Supplementary Fig. 10. Nominal EI-MS spectra confirm peak identity as (*S*)-8-oxolinalool ((*S*)-7).**

Comparison of nominal EI-MS data obtained for (**A**) triungulin DHV collection extract peak ($t_r$ 58.70 min, **Supplementary Fig. 9**) and (**B**) (*S*)-**7** reference standard.

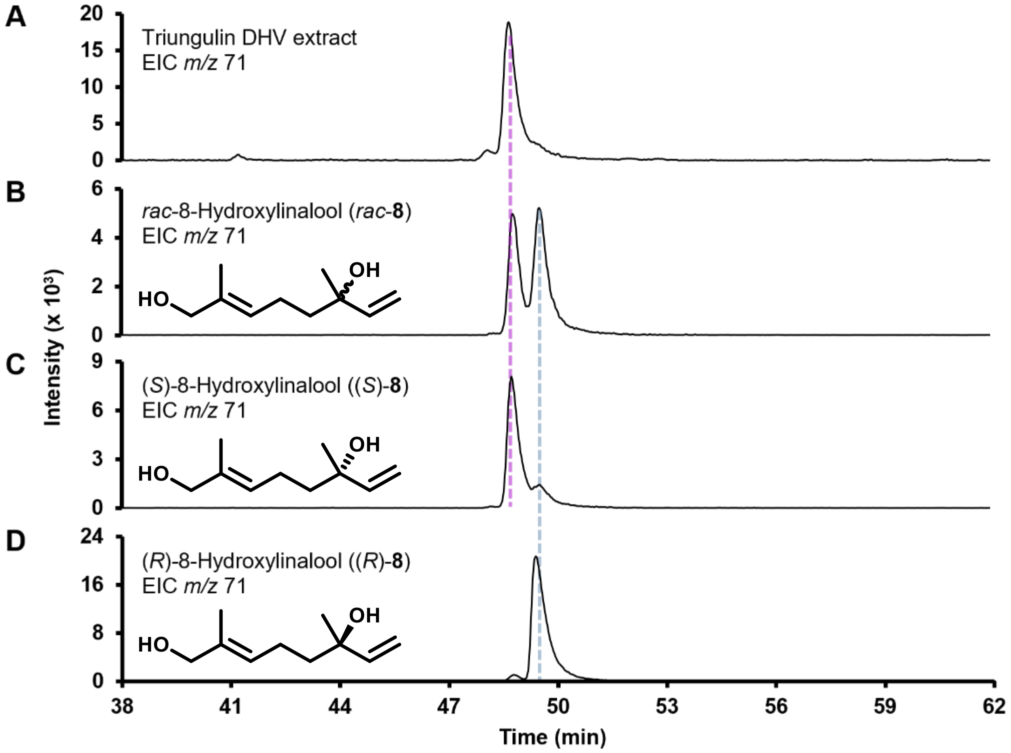


**Supplementary Fig. 11. Triungula emit (*S*)-8-hydroxylinalool ((*S*)-8).**

Comparison of chiral GC-EI-MS chromatographic data obtained for (**A**) triungula DHV collection extract, (**B**) *rac*-8-hydroxylinalool (*rac*-**8**), (**C**) (*S*)-**8**, and (**D**) (*R*)-**8**. Separation method: E.

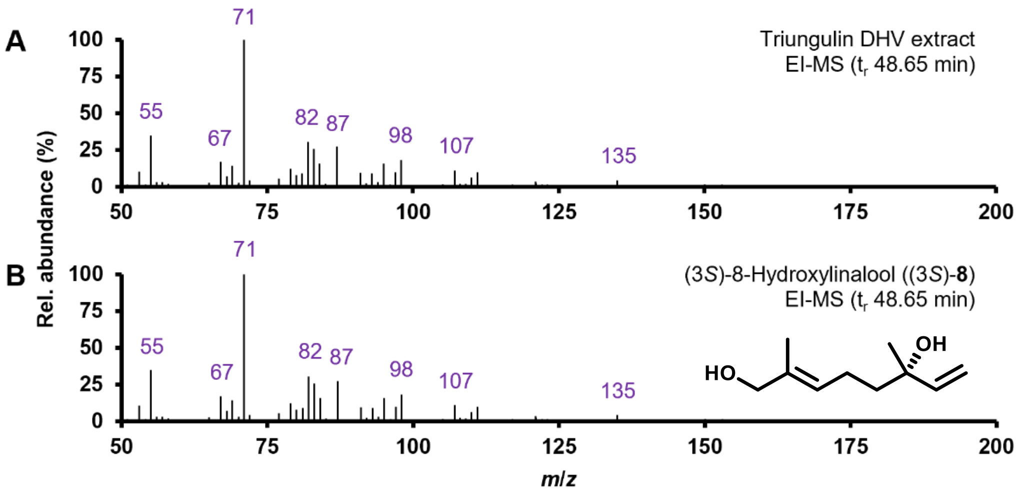


**Supplementary Fig. 12. Nominal EI-MS spectra confirm peak identity as (*S*)-8-hydroxylinalool ((*S*)-8).**

Comparison of nominal EI-MS data obtained for (**A**) triungulin DHV collection extract peak ($t_r$ 48.65 min, **Supplementary Fig. 11**) and (**B**) (*S*)-**8** reference standard.

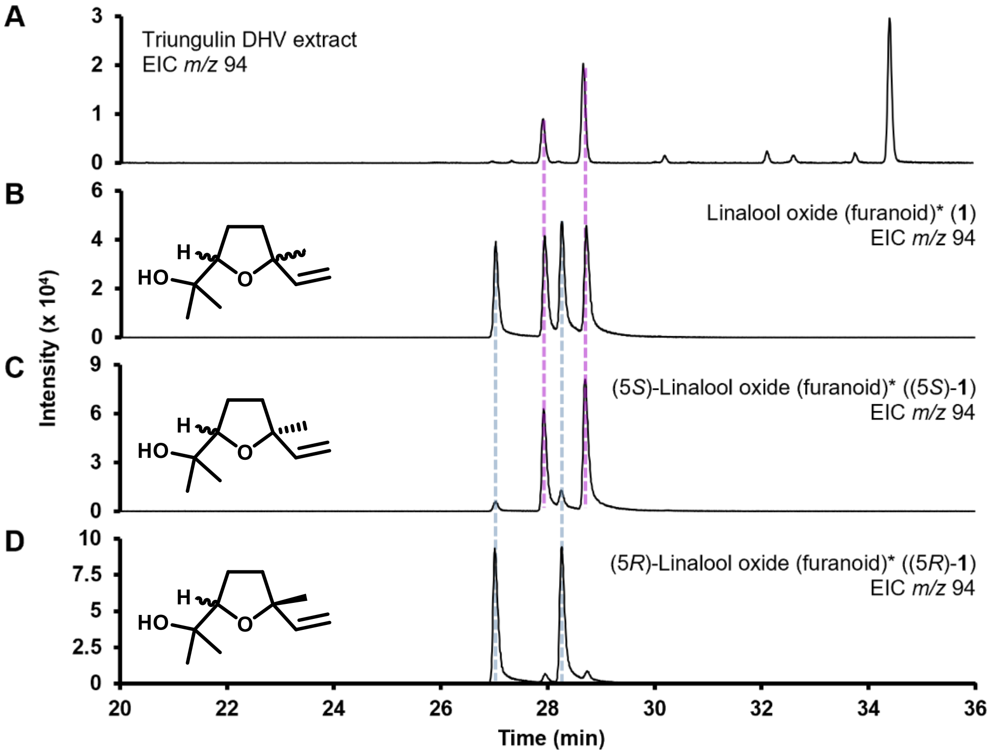


**Supplementary Fig. 13. Triungula emit (2*S*,5*S*)- and (2*R*,5*S*)-linalool oxide (furanoid) ((5*S*)-1).**

Comparison of chiral GC-EI-MS chromatographic data obtained for (**A**) triungula DHV collection extract, (**B**) linalool oxide (furanoid) (**1**), (**C**) (5*S*)-**1**, and (**D**) (5*R*)-**1**. Separation method: D. *Mixture of stereoisomers.

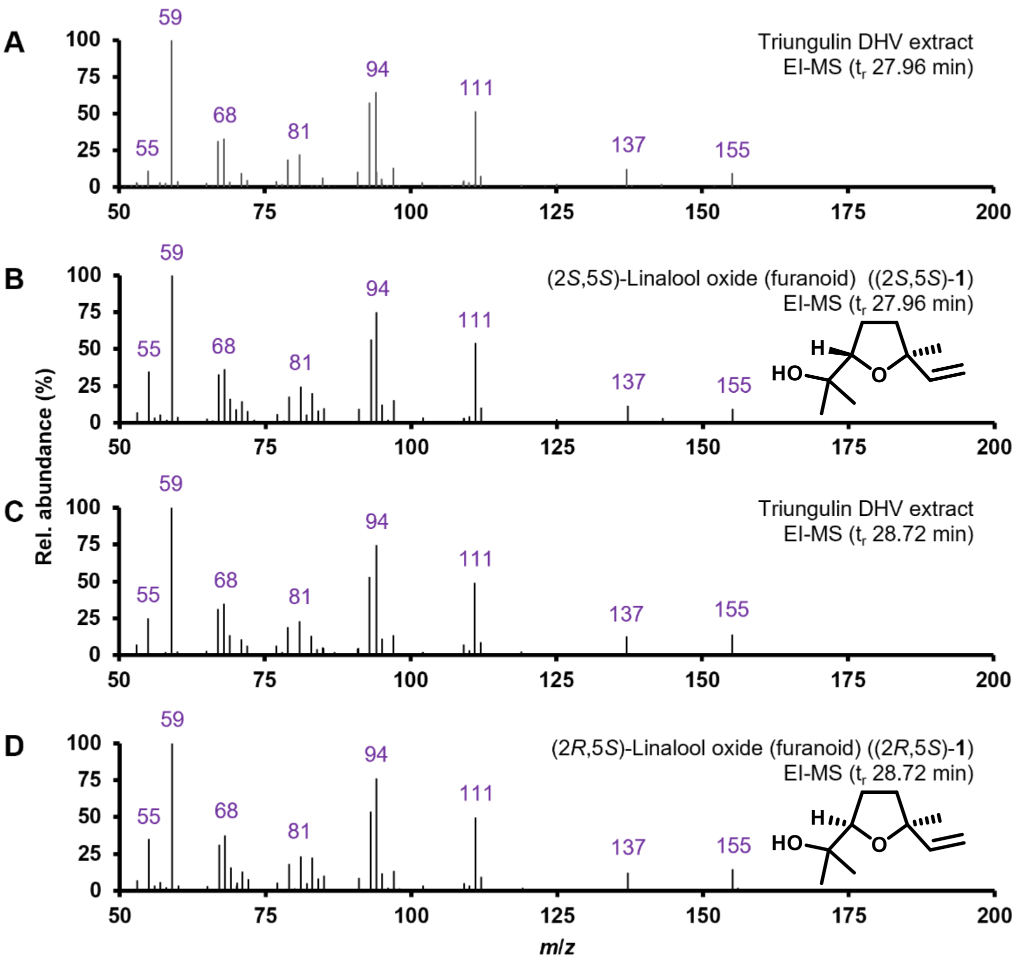


**Supplementary Fig. 14. Nominal EI-MS spectra confirm peak identities as both stereoisomers of (5*S*)-linalool oxide (furanoid) ((5*S*)-1).**

Comparison of nominal EI-MS data obtained for (**A**) triungulin DHV collection extract peak ($t_r$ 27.96 min, **Supplementary Fig. 13**), (**B**) (2*S*,5*S*)-**1** ($t_r$ 27.96 min, **Supplementary Fig. 13**), (**C**) triungulin DHV collection extract peak ($t_r$ 28.72 min, **Supplementary Fig. 13**), and (**D**) (2*R*,5*S*)-**1** ($t_r$ 28.72 min, **Supplementary Fig. 13**).

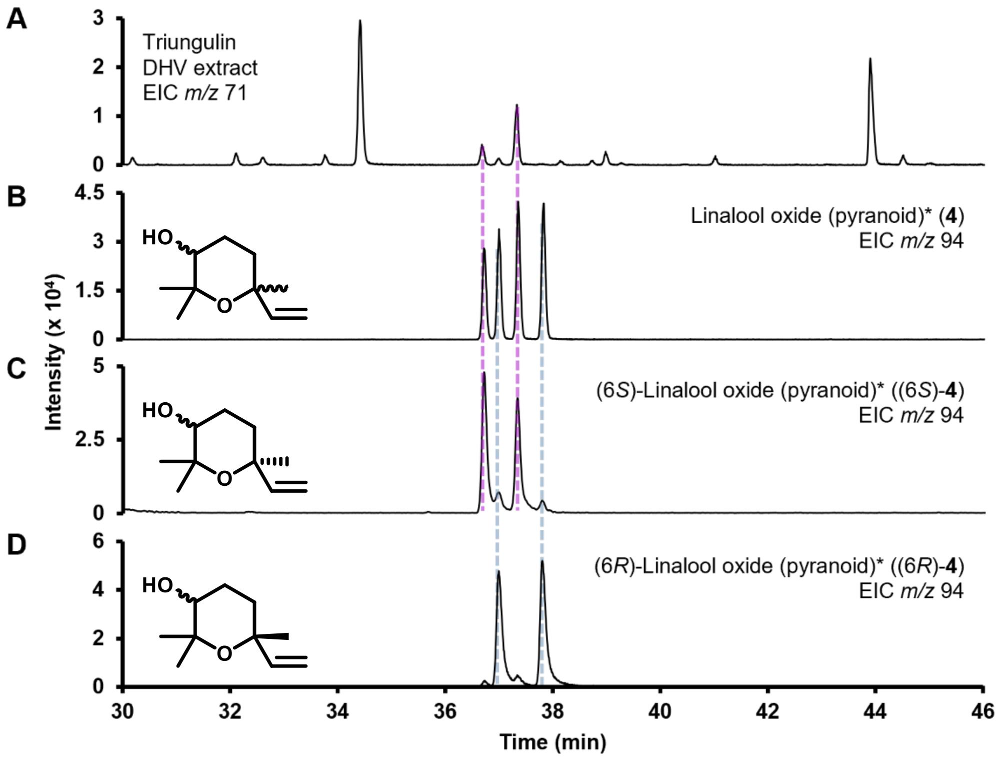


**Supplementary Fig. 15. Triungula emit (3*R*,6*S*)- and (3*S*,6*S*)-linalool oxide (pyranoid) ((6*S*)-4).**

Comparison of chiral GC-EI-MS chromatographic data obtained for (**A**) triungula DHV collection extract, (**B**) linalool oxide (pyranoid) (**4**), (**C**) (6*S*)-**4**, and (**D**) (6*R*)-**4**. Separation method: D. *Mixture of stereoisomers.

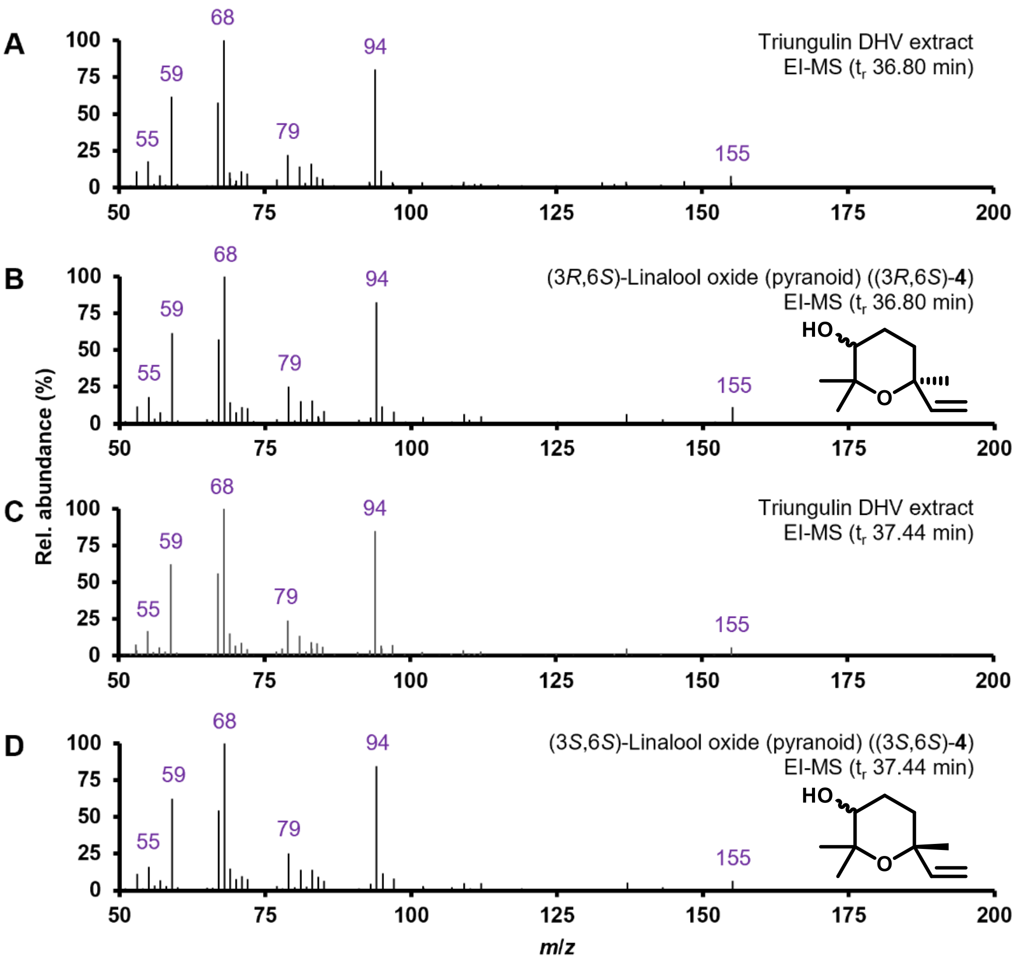


**Supplementary Fig. 16. Nominal EI-MS spectra confirm peak identities as both stereoisomers of (6*S*)-linalool oxide (pyranoid).**

Comparison of nominal EI-MS data obtained for (**A**) triungulin DHV collection extract peak (t$_r$ 36.80 min, **Supplementary Fig. 15**), (**B**) (3*R*,6*S*)-**4** (t$_r$ 36.80 min, **Supplementary Fig. 15**), (**C**) triungulin DHV collection extract peak (t$_r$ 37.44 min, **Supplementary Fig. 15**), and (**D**) (3*S*,6*S*)-**4** (t$_r$ 37.44 min, **Supplementary Fig. 15**).

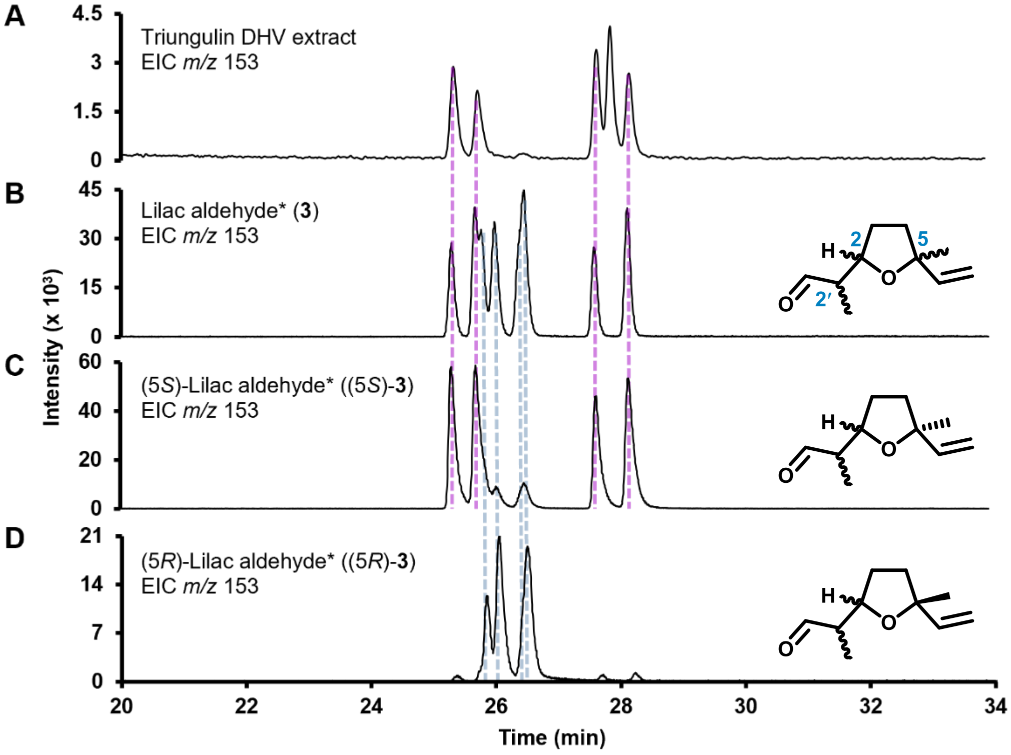


**Supplementary Fig. 17. Triungula emit (2´*S*,2*S*,5*S*)-, (2´*R*,2*S*,5*S*)-, (2´*S*,2*R*,5*S*)-, and (2´*R*,2*R*,5*S*)-lilac aldehyde ((5*S*)-3).**

Comparison of chiral GC-EI-MS chromatographic data obtained for (**A**) triungula DHV collection extract, (**B**) lilac aldehyde (**3**), (**C**) (5*S*)-**3**, and (**D**) (5*R*)-**3**. Separation method: C. *Mixture of stereoisomers.

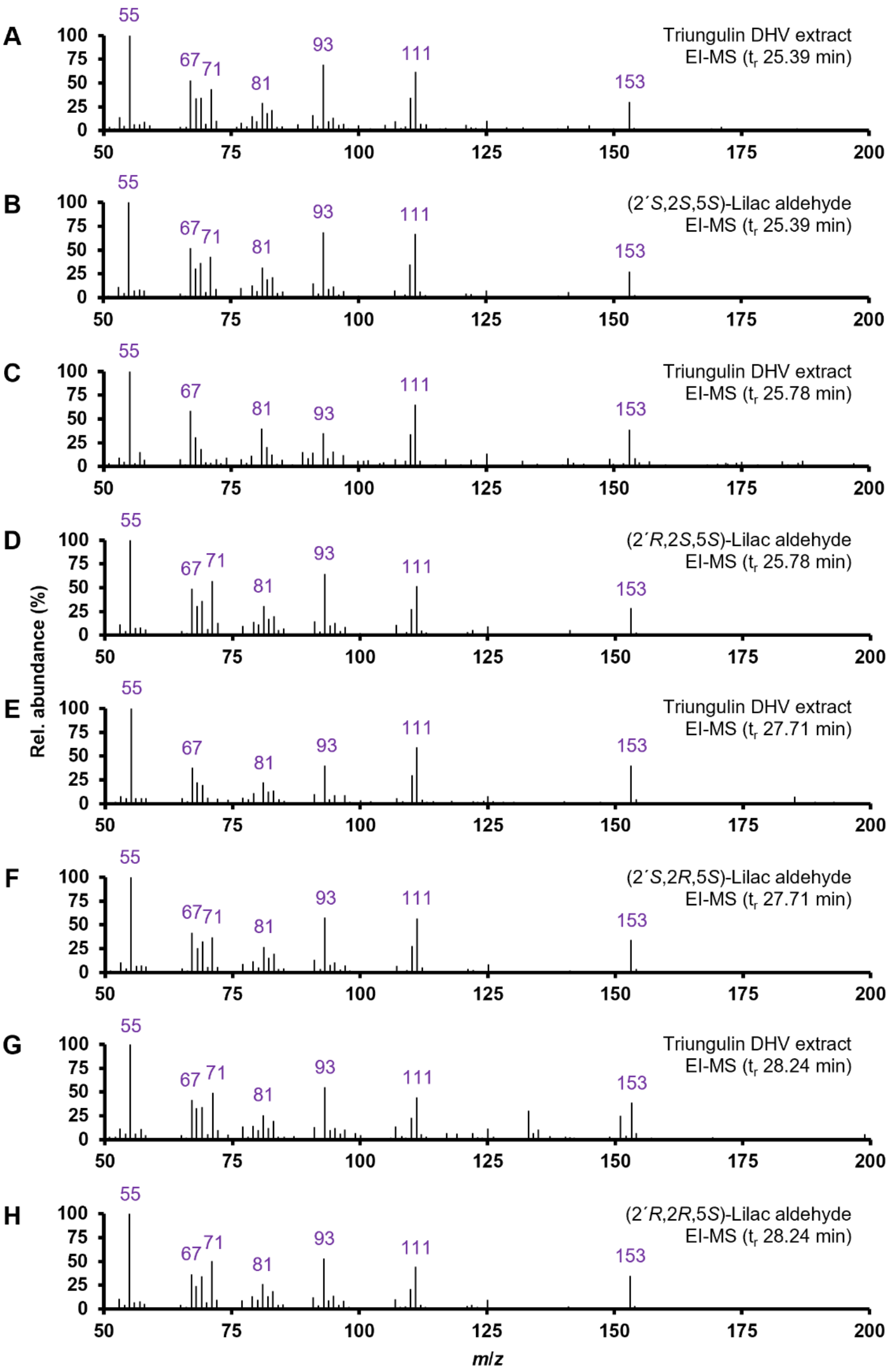


**Supplementary Fig. 18. Nominal EI-MS spectra confirm peak identities of four stereoisomers of (5*S*)-lilac aldehyde ((5*S*)-3).**

Comparison of nominal EI-MS data obtained for (**A**) triungulin DHV collection extract peak ($t_r$ 25.39 min, **Supplementary Fig. 17**), (**B**) (2´*S*,2*S*,5*S*)-**3** ($t_r$ 25.39 min, **Supplementary Fig. 17**),

(**C**) triungulin DHV collection extract peak ($t_r$ 25.78 min, **Supplementary Fig. 17**), (**D**) (2´*R*,2*S*,5*S*)-**3** ($t_r$ 25.78 min, **Supplementary Fig. 17**), (**E**) triungulin DHV collection extract peak ($t_r$ 27.71 min, **Supplementary Fig. 17**), (**F**) (2´*S*,2*R*,5*S*)-**3** ($t_r$ 27.71 min, **Supplementary Fig. 17**), (**G**) triungulin DHV collection extract peak ($t_r$ 28.24 min, **Supplementary Fig. 17**), and (**H**) (2´*R*,2*R*,5*S*)-**3** ($t_r$ 28.24 min, **Supplementary Fig. 17**).

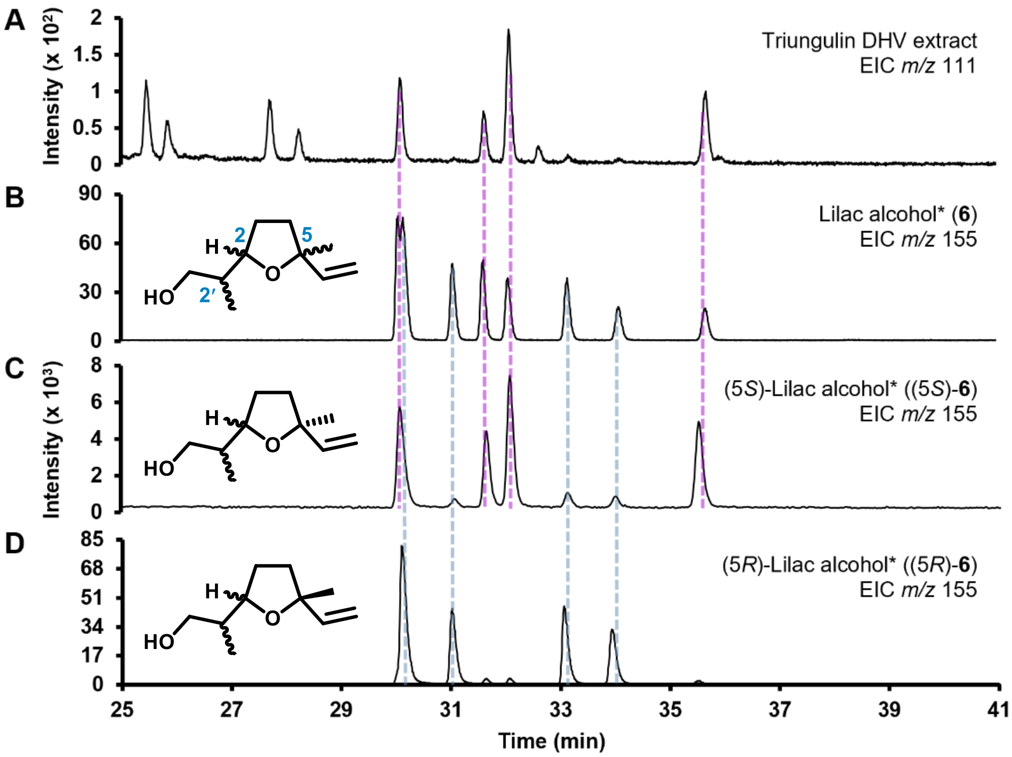


**Supplementary Fig. 19. Triungula emit (2´*S*,2*S*,5*S*)-, (2´*S*,2*R*,5*S*)-, (2´*R*,2*S*,5*S*)-, and (2´*R*,2*R*,5*S*)-lilac alcohol (((5*S*)-6).**

Comparison of chiral GC-EI-MS chromatographic data obtained for (**A**) triungula DHV collection extract, (**B**) lilac alcohol (**6**), (**C**) (5*S*)-**6**, and (**D**) (5*R*)-**6**. Separation method: C. *Mixture of stereoisomers.

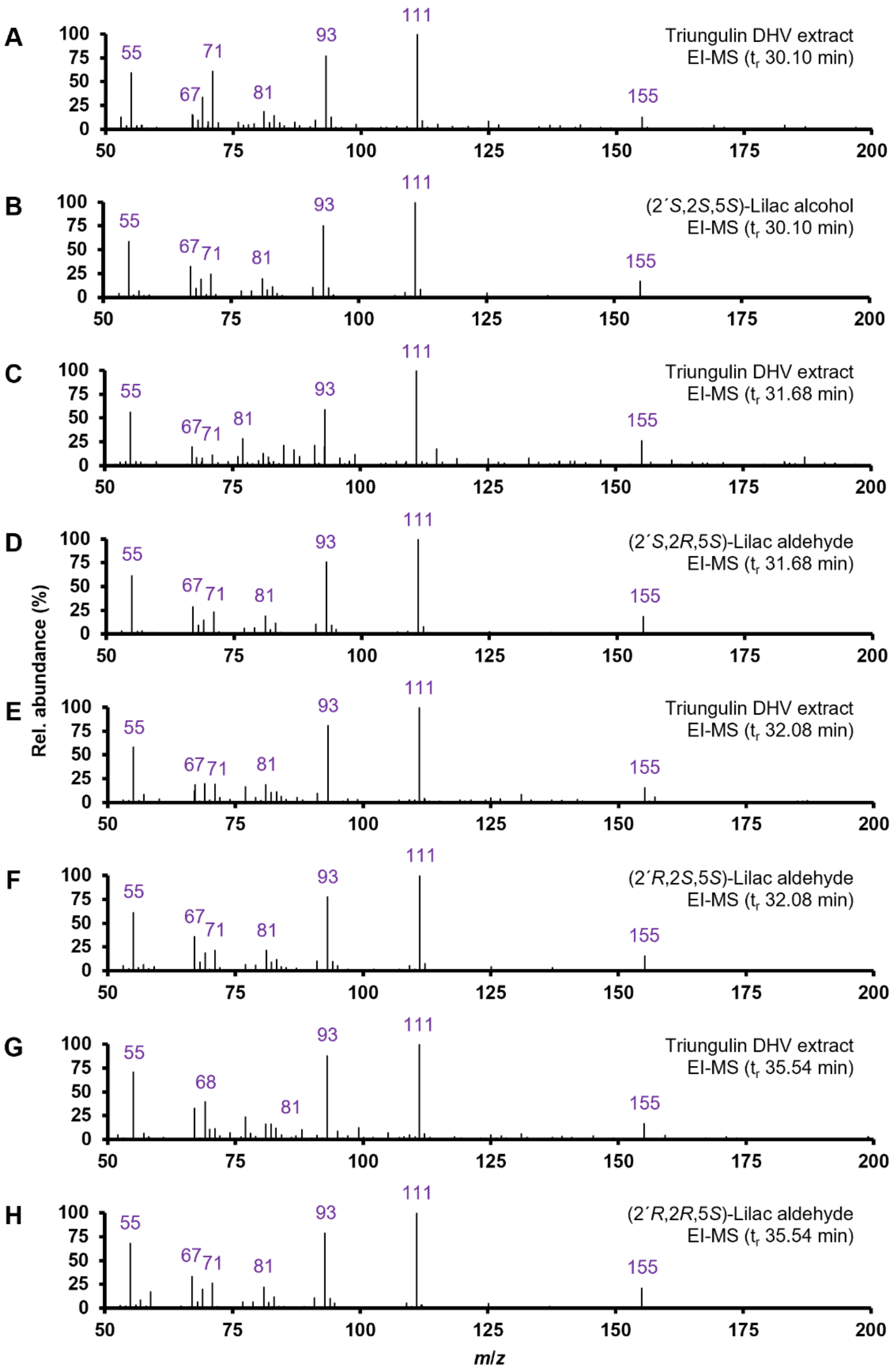


**Supplementary Fig. 20. Nominal EI-MS spectra confirm peak identities of four stereoisomers of (5*S*)-lilac alcohol ((5*S*)-6).**

Comparison of nominal EI-MS data obtained for (**A**) triungulin DHV collection extract peak ($t_r$ 30.10 min, **Supplementary Fig. 19**), (**B**) (2´*S*,2*S*,5*S*)-**6** ($t_r$ 30.10 min, **Supplementary Fig. 19**),

(**C**) triungulin DHV collection extract peak ($t_r$ 31.68 min, **Supplementary Fig. 19**), (**D**) (2´*S*,2*R*,5*S*)-**6** ($t_r$ 31.68 min, **Supplementary Fig. 19**), (**E**) triungulin DHV collection extract peak ($t_r$ 32.08 min, **Supplementary Fig. 19**), (**F**) (2´*R*,2*S*,5*S*)-**6** ($t_r$ 32.08 min, **Supplementary Fig. 19**), (**G**) triungulin DHV collection extract peak ($t_r$ 35.54 min, **Supplementary Fig. 19**), and (**H**) (2´*R*,2*R*,5*S*)-**3** ($t_r$ 35.54 min, **Supplementary Fig. 19**).

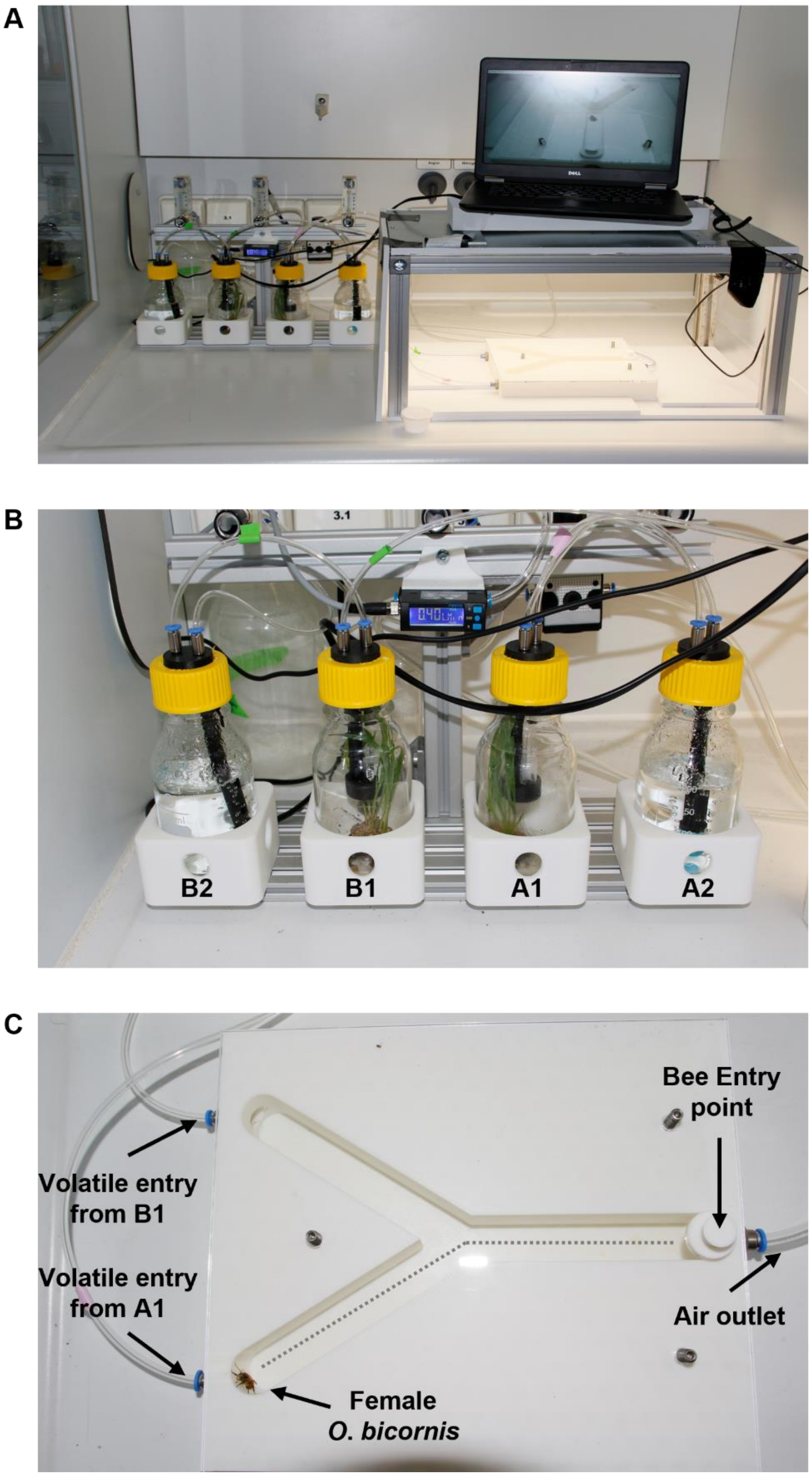


**Supplementary Fig. 21. Y-tube dual-choice olfactometric assay setup.**

(**A**) Dual-choice Y-tube assay setup. (**B**) Charcoal-filtered compressed air was first bubbled through distilled water (Bottles A2 or B2) and then passed over freshly sprouted *Triticum aestivum* and a glass vial containing a solution of synthetic (*S*)-**2**-derived monoterpenoids in $CH_2Cl_2$ (Bottle A1) or neat $CH_2Cl_2$ (Bottle B1) which then flowed into the Y-tube apparatus. Note: Volatile-rich air entering sidearms of the Y-tube olfactometer from Bottles A1 and B1 were switched between each experiment. (**C**) 3D-printed Y-tube apparatus showing entry ports for test bee and volatile-rich air from Bottles A1 and B1.

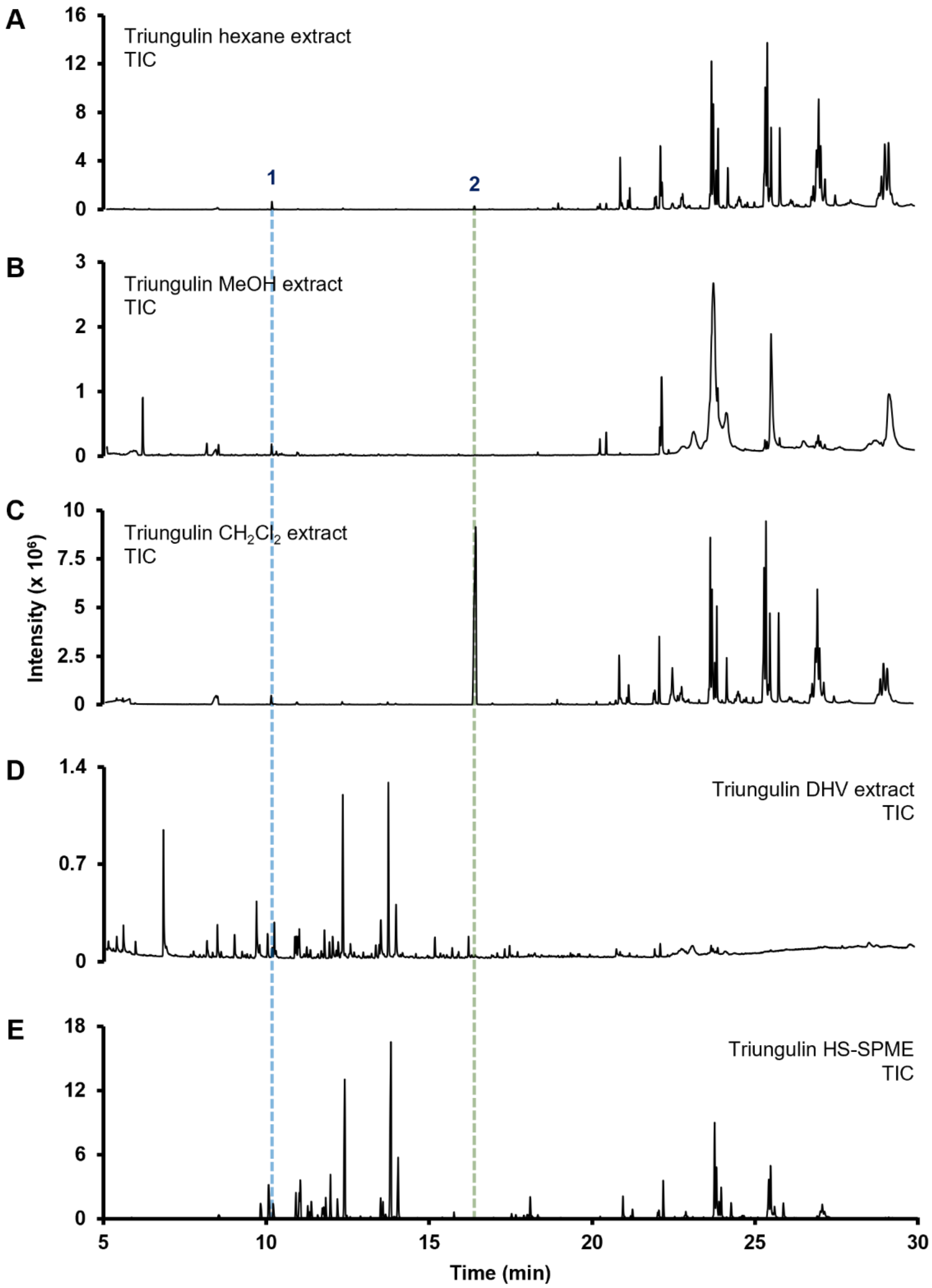


**Supplementary Fig. 22. Comparison of triungulin volatile extraction methods.**

Comparing GC-EI-MS TIC chromatograms obtained for whole-body triungula extracts, using (**A**) hexane, (**B**) MeOH, and (**C**) $CH_2Cl_2$, with those obtained for (**D**) DHV collection and (**E**) HS-SPME sampling, show that solvent-based extraction methods fail to recover the complete monoterpenoid profile. Peak identities: *1*) linalool, *2*) cantharidin*. Separation method: A. * tentatively identified using the National Institute of Standards and Technology (NIST) MS-Library v. 3.0 (2023).

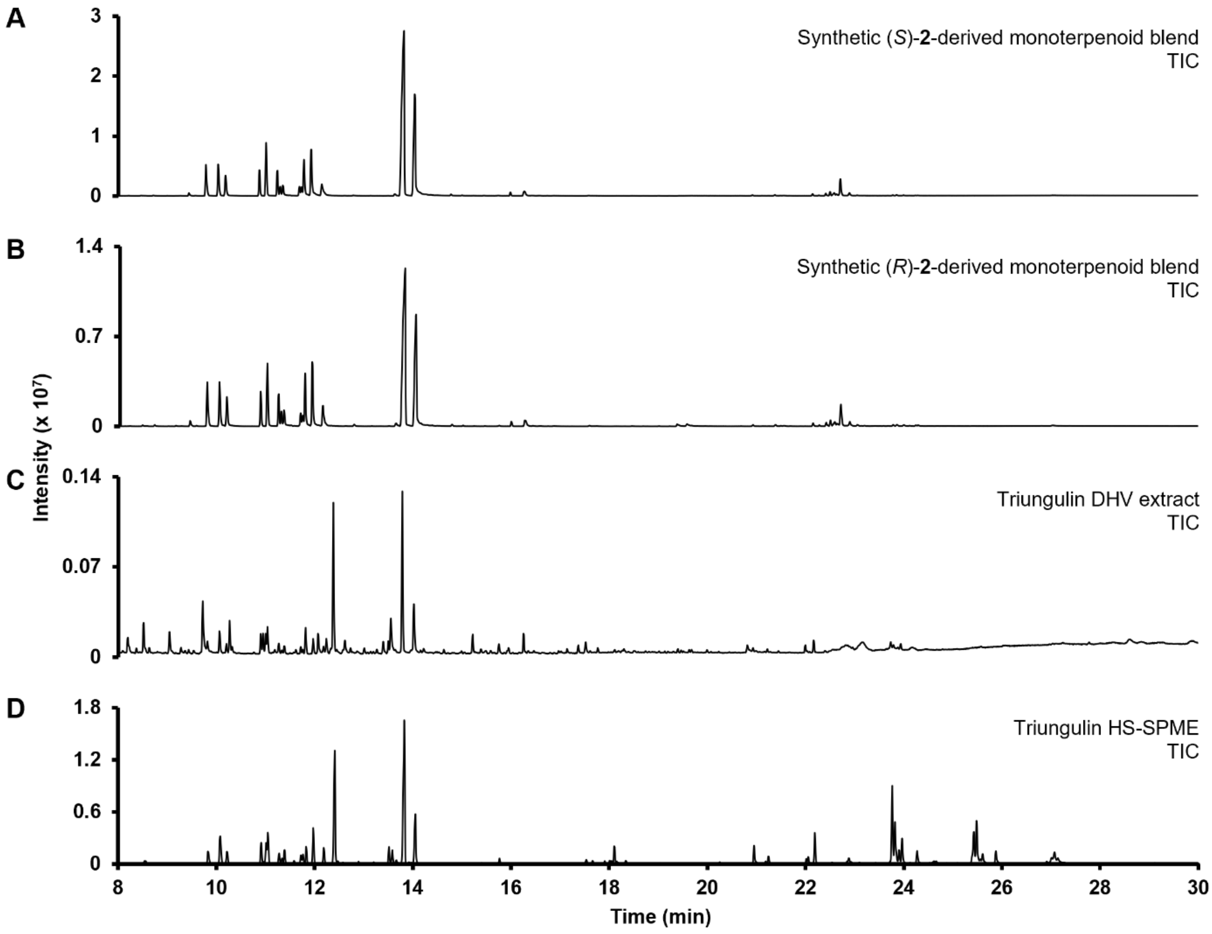


**Supplementary Fig. 23. Comparison of synthetic linalool (2)-derived monoterpenoid blends with the naturally-occurring triungulin volatile profile.**

Comparison of GC-EI-MS data obtained for (**A**) (*S*)-**2**- and (**B**) (*R*)-**2**-derived synthetic monoterpenoid blends lacking long-chain hydrocarbons present to varying degrees in triungulin volatile profiles obtained using (**C**) DHV collection and (**D**) HS-SPME ($t_r$ 20.95–29.22 min). Separation method: A.

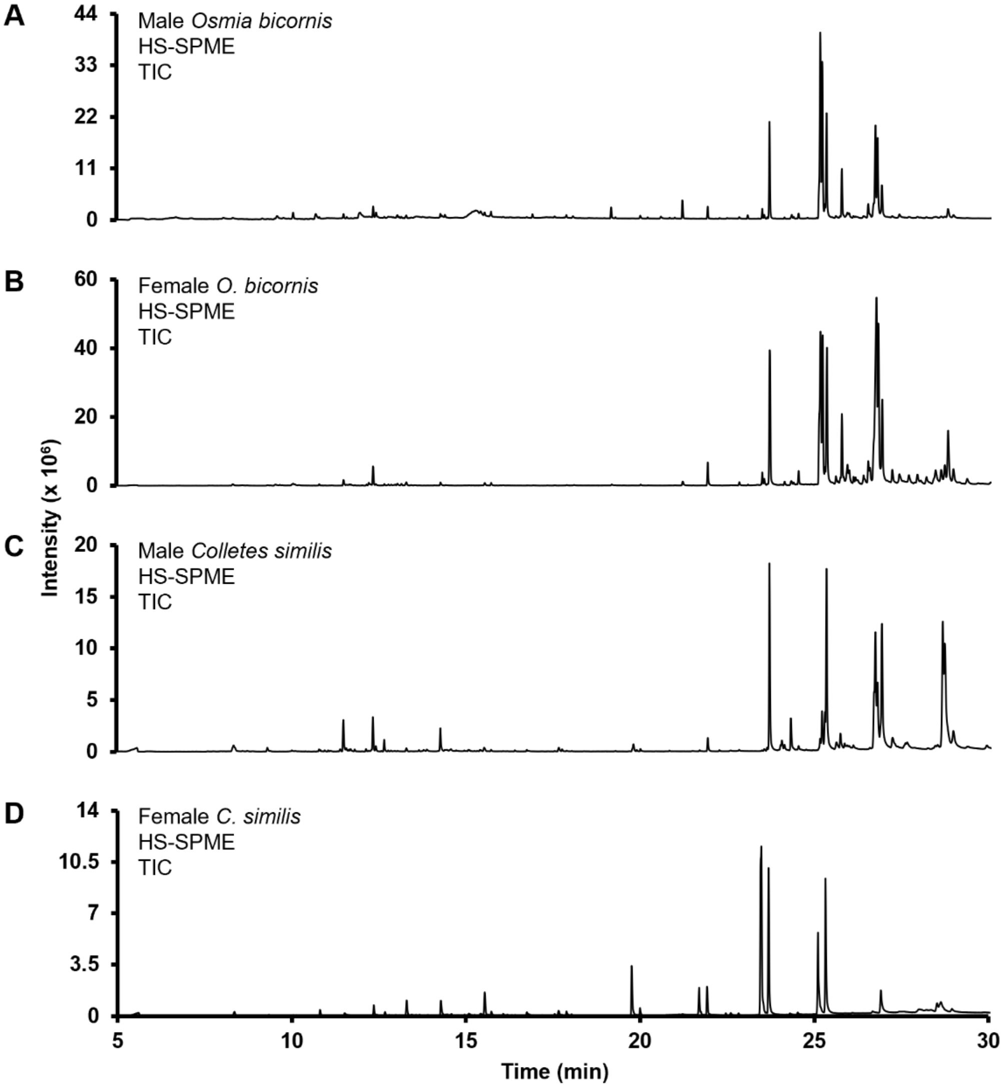


**Supplementary Fig. 24. Volatile profiles of male and female *O. bicornis* and *C. similis*.** HS-SPME GC-EI-MS analysis of (**A**) male *O. bicornis*, (**B**) female *O. bicornis*, (**C**) male *C. similis*, and (**D**) female *C. similis* failed to detect any monoterpenoids volatiles. Separation method: A.

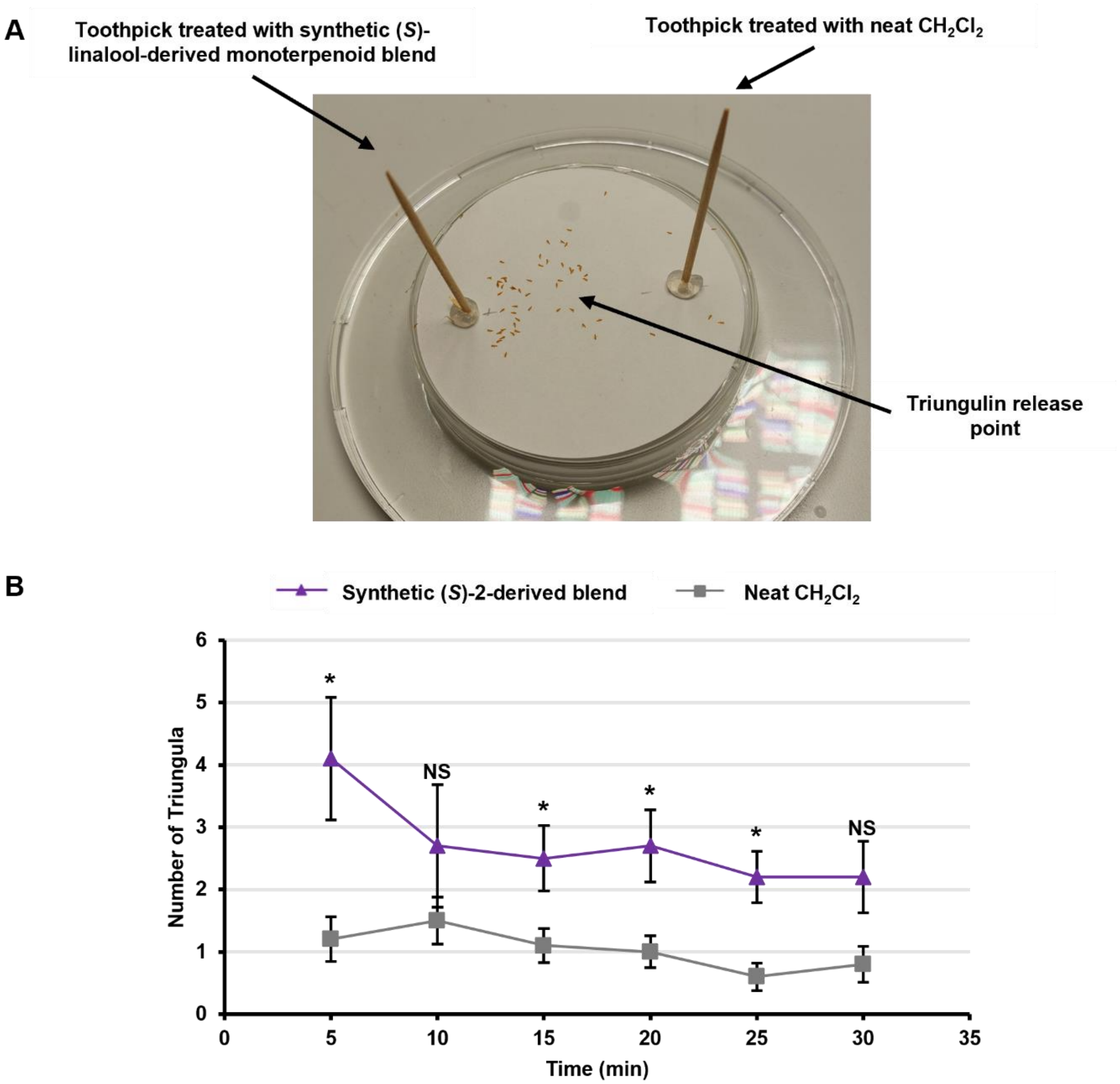


**Supplementary Fig. 25. Olfactometric assays assessing triungulin preference for (*S*)-linalool-derived volatiles.**

(**A**) An in-house setup was assembled to assess triungulin preference for a synthetic blend of (*S*)-linalool-derived monoterpenoids. (**B**) Line graph showing average number of triungula on a toothpick treated with a (*S*)-linalool-derived blend in $CH_2Cl_2$ versus a toothpick treated with neat $CH_2Cl_2$ after 5, 10, 15, 20, 25, and 30 min (error bars at each timepoint denote standard error of the mean [SEM]; $n$ = 10). Paired t-test: paired two sample for means: *$p < 0.05$, NS = not significant. For further data, see **Supplementary Table 1**.

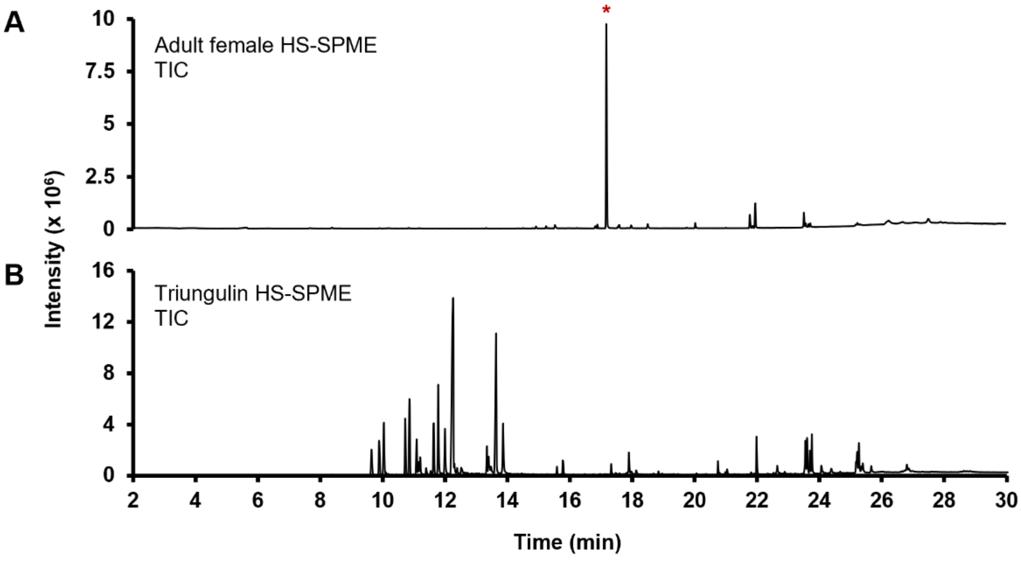


**Supplementary Fig. 26. Comparison of volatile profiles for adult female *M. proscarabaeus* and triungula show that adult females do not emit monoterpenoid volatiles.**

HS-SPME GC-EI-MS data obtained for (**A**) adult female *M. proscarabaeus* and (**B**) triungula. Separation method: A. *Unidentified sesquiterpenoid volatile.

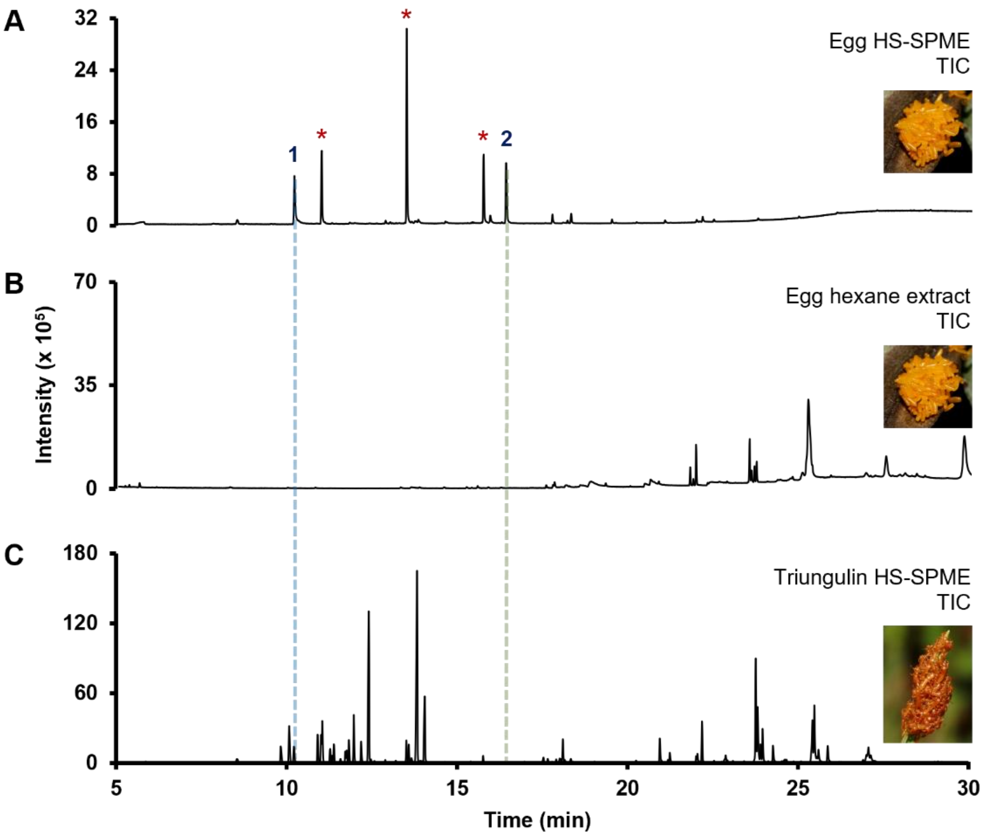


**Supplementary Fig. 27. *M. proscarabaeus* triungula emit monoterpenoid bouquet following eclosion.**

Comparison of GC-EI-MS data obtained following (**A**) HS-SPME or (**B**) hexane extraction of eggs with (**C**) the volatile profile of triungula indicate that the only monoterpenoid emitted by eggs is linalool. Peak identity: *1*) linalool and *2*) cantharidin. Separation method: A.*Background. Separation method: B.

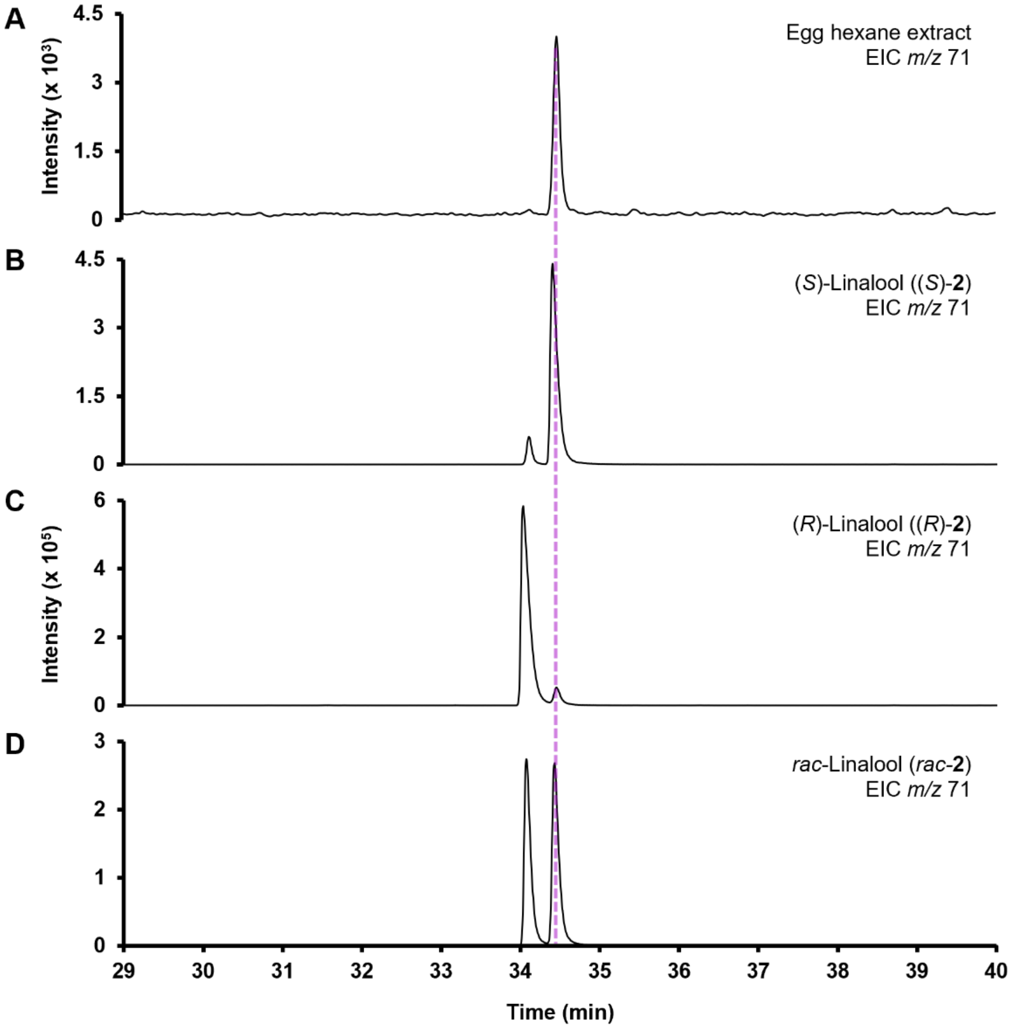


**Supplementary Fig. 28. *M. proscarabaeus* eggs emit (*S*)-linalool ((*S*)-2).**
Comparison of GC-EI-MS data obtained for (**A**) a hexane extract from eggs, (**B**) (*S*)-**2**, (**C**) (*R*)-**2**, and (**D**) *rac*-**2**, showed that eggs emit (*S*)-**2** exclusively. Separation method: B.

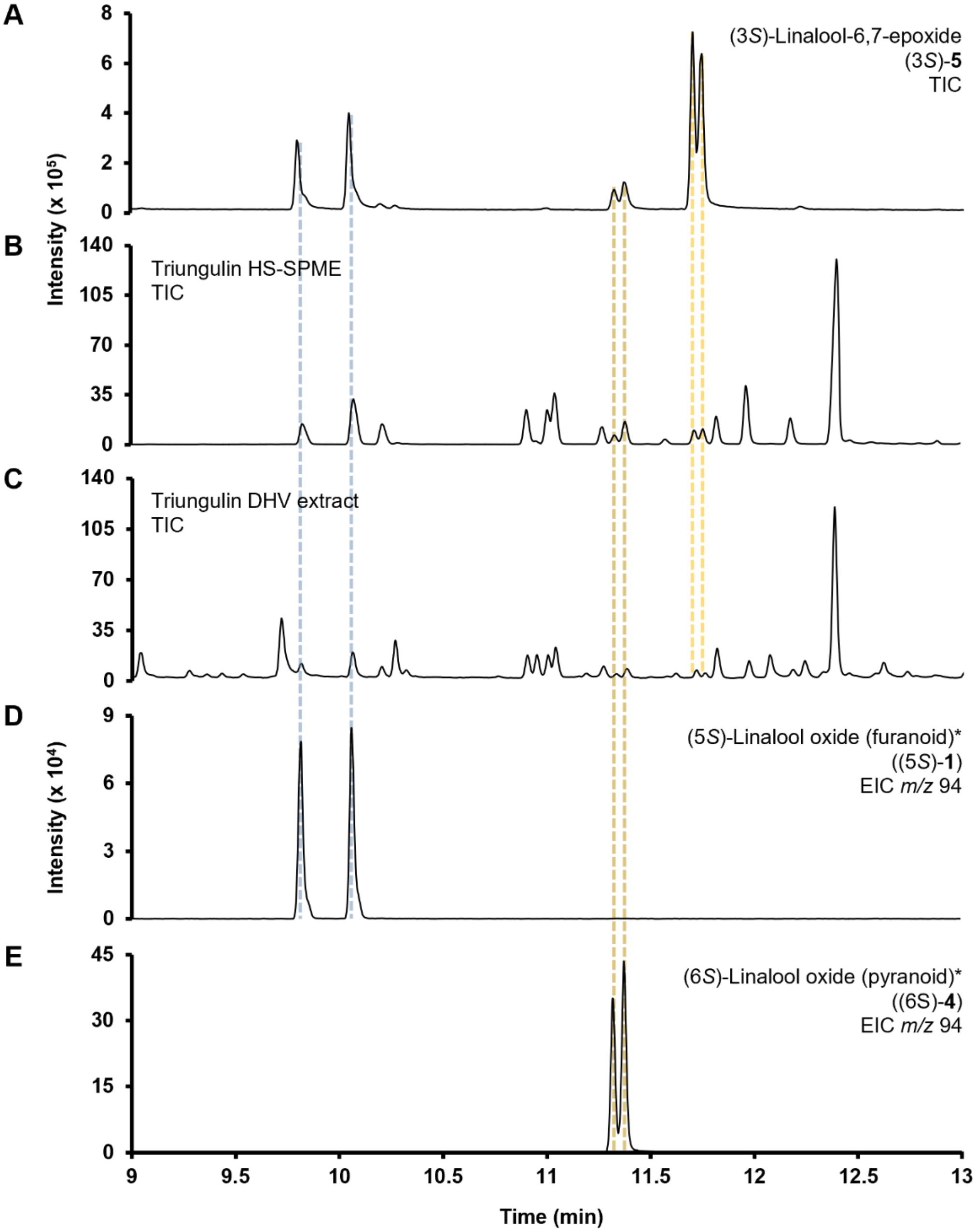


**Supplementary Fig. 29. GC-EI-MS data showing spontaneous cyclization of (3*S*)-linalool-6,7-epoxide ((3*S*)-5) to linalool oxides (2*S*)-1 and (6*S*)-4 during analysis.**

Comparison of GC-EI-MS data for (**A**) freshly prepared linalool-6,7-epoxide ((3*S*)-**5**), (**B**) triungulin volatiles collected by HS-SPME and (**C**) DHV collection, (**D**) (5*S*)-linalool oxide (furanoid)* ((5*S*)-**1**), and (**E**) (6*S*)-linalool oxide (pyranoid)* ((6*S*)-**4**). *Mixture of stereoisomers.

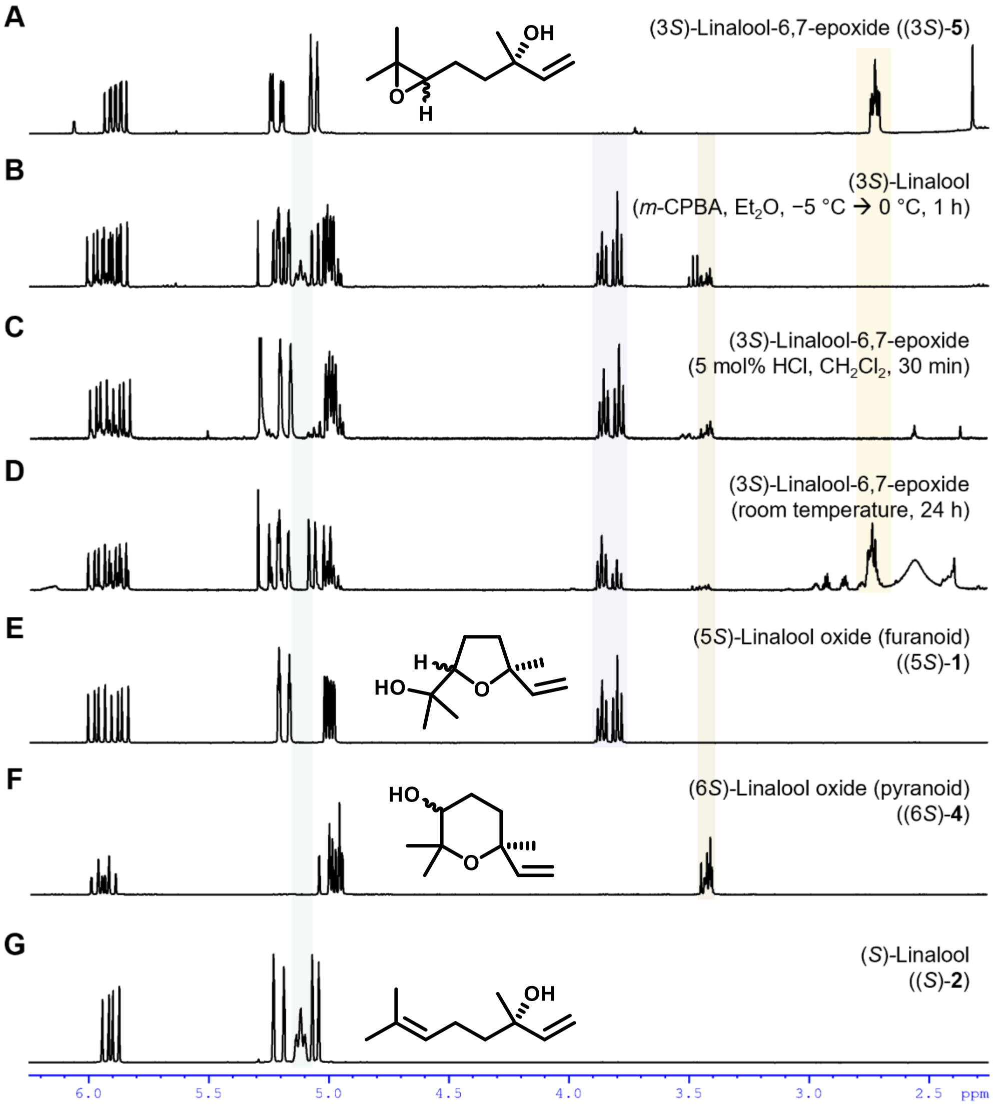


**Supplementary Fig. 30. [1]H NMR data showing spontaneous cyclization of (3*S*)-linalool-6,7-epoxide ((3*S*)-5) to linalool oxides (2*S*)-1 and (6*S*)-4 under various conditions.**

Comparison of $^1$H NMR ($CDCl_3$, 400 MHz) spectral data for (**A**) freshly prepared (3*S*)-linalool-6,7-epoxide ((3*S*)-**5**), (**B**) the crude reaction mixture following attempted epoxidation of (*S*)-linalool ((*S*)-**2**) with *m*-chloroperoxybenzoic acid (1 eq. *m*-CPBA, $CH_2Cl_2$, Ar, −5 °C → 0 °C, 2 h), (**C**) the crude residue arising from Brønsted acid-catalyzed intramolecular cyclization of epoxide (3*S*)-**5** (5 mol% HCl, $CH_2Cl_2$, Ar, rt, 30 min), (**D**) epoxide (3*S*)-**5** stored at room temperature for 24 h, (**E**) (5*S*)-linalool oxide (furanoid)* ((5*S*)-**1**), (F) (6*S*)-linalool oxide (pyranoid)* ((6*S*)-**4**), and (G) (*S*)-linalool ((*S*)-**2**).

**Supplementary Table 1. Triungulin olfactometric choice assay results.**

| | Time (min) | | | | | | | | | | | |
|---|---|---|---|---|---|---|---|---|---|---|---|---|
| | 5* | | $10^{NS}$ | | 15* | | 20* | | 25* | | $30^{NS}$ | |
| **Entry** | Treat. | Con. | Treat. | Con. | Treat. | Con. | Treat. | Con. | Treat. | Con. | Treat. | Con. |
| 1 | 0 | 3 | 0 | 3 | 0 | 2 | 0 | 2 | 0 | 1 | 0 | 2 |
| 2 | 0 | 0 | 0 | 0 | 3 | 0 | 4 | 0 | 4 | 0 | 4 | 0 |
| 3 | 5 | 0 | 2 | 0 | 2 | 0 | 2 | 0 | 2 | 0 | 2 | 0 |
| 4 | 1 | 0 | 2 | 0 | 1 | 1 | 1 | 1 | 1 | 0 | 1 | 2 |
| 5 | 6 | 3 | 4 | 2 | 3 | 2 | 3 | 1 | 3 | 1 | 3 | 0 |
| 6 | 10 | 1 | 2 | 2 | 2 | 0 | 2 | 1 | 2 | 0 | 1 | 0 |
| 7 | 6 | 1 | 5 | 1 | 6 | 2 | 5 | 2 | 4 | 1 | 6 | 1 |
| 8 | 4 | 1 | 5 | 3 | 2 | 1 | 2 | 0 | 2 | 0 | 1 | 0 |
| 9 | 5 | 1 | 5 | 2 | 2 | 1 | 2 | 1 | 1 | 1 | 1 | 1 |
| 10 | 4 | 2 | 2 | 2 | 4 | 2 | 6 | 2 | 3 | 2 | 3 | 2 |

Paired t-test: paired two sample for means: $*p < 0.05$, NS = not significant. Treat. = treatment ((*S*)-linalool-derived monoterpenoid blend in $CH_2Cl_2$); Con. = Control (neat $CH_2Cl_2$).

**Supplementary Table 2. Primers used for coding sequence amplification from triungulin cDNA.**

| Gene | Direction | Note | Sequence |
|---|---|---|---|
| MpRed | Forward | *Sal*I | ACCGTCGACATGGACGAAACAGAAGCG |
| | Reverse | *Nhe*I | GTCGCTAGCTTAACTCCAAACATCAGCTGA |
| CYP347BT1 | Forward | *Not*I | CACTAAAGGGCGGCCGCAATGTTAATAGTTTTGGTTAATGTCTG |
| | Reverse | *Sac*I | GAATTGTTAATTAAGAGCTCCTAAACTGCGTTGATTCTTAGAG |
| CYP345BZ1 | Forward | *Not*I | CACTAAAGGGCGGCCGCAATGATTTTGATAAGTCTAGTTCTAGTATT |
| | Reverse | *Sac*I | GAATTGTTAATTAAGAGCTCTTATAGTTCCCTTTCTTCAAAATTAATC |

**Supplementary Table 3. Sequences amplified from triungulin cDNA for this study.**

| Gene | Sequence |
|---|---|
| MpRed | ATGGACGAAACAGAAGCGAAAATTGGTGCCCAACAAATACCTGAAACTGTTGAAAG<br>TTTATTTACTACATTAGATTATATATTACTTGCCCTTTTAATTGGTGGTGTAAGTTATT<br>GGTATTTTAATCGGCAGAAAAAGAAAGAAGTAACAACTACGCGATCCTATACTATAC<br>AGCCAACTTCTATGACATTACAAGCAACAACAGAGAGTTCATTTATAAAAAAATTAA<br>AAGCATCAGGACGTTCATTGGTTGTATTTTATGGTAGTCAAACTGGTACTGGTGAAG<br>AATTTGCTGGACGTTTAGCTAAAGAAGGTTTACGTTATGGTATGAAAGGGATGATAG<br>CTGATCCAGAAGAATGCGATATGGAGGAATTAGTGAATTTAAAAGCAATACCAAATT<br>CATTAGCTGTATTTTGTATGGCAACATATGGTGAAGGTGACCCAACTGATAATGCAA<br>TGGAATTTTATGAATGGTTACAAAATGGTGATGCTGATTTAACTGGCTTAAATTATTC<br>GGTATTCGGCTTAGGTAATAAAACGTATGAACATTACAATGAAGTCGCAATCTACAT<br>AGATAAACGTTTAGAAGATTTAGGAGCAACTCGAGTGTTTGATTTAGGACTTGGCGA<br>TGATGATGCTAATATTGAAGACGATTTCATCACTTGGAAAGATAAGTTCTGGCCAGC<br>AGTTTGTGAATTCTTCGGTATCGAGTCAACTGGTGAAGATATCAGTATACGTCAATAT<br>CGCTTACAAGAATTCACTGATGAATTACCAGAACGTCTTTATACAGGTGAAATGGCA<br>CGATTGCATTCACTTAAAAATCAAAGACCACCTTATGATGCAAAGAATCCATTTTTG<br>GCGAAAATTTTAGTAAATCGTGAATTACATAAAAATGGTGATCGTTCATGTATGCAT<br>ATCGAATTTGATATTAGCGGTTCAAAGATGCGTTATGATTCTGGCGATCATTTAGCCG<br>TTTATCCAATTAATAATCATGAATTGGTTGAGAAAATTGGCAAATATACTGGTCAAG<br>ATTTAGATACTGTGTTCACTTTAGTAAATACTGATGAAGAATCAAGTAAAAAACATC<br>CATTCCCCTGTCCAACATCTTATCGTACAGCTTTAACACATTATCTTGATATCACAAT<br>GAATCCAAGAACGCACGTTTTAAAAGAATTATCGGAATATTGTAGTGATCCAAATGA<br>AAAAGAAAAACTAAAATTAATGGCCAGTATATGTCCAGAAGGTAAAGCATTATATC<br>AACAATGGATAAACGATGATAATCGTAATATTGCACAAATATTAGAAGATATGCCGT<br>CGTGTCGGCCAGCTTTAGATCATTTATGTGAATTATTACCACGTTTACAACCACGTTA<br>TTATTCAATATCATCATCTGCAAAACTATATCCAAATACTGTACATATAACAGCTGTT<br>GTTGTCGAATATAAAACACCAACAGGACGTATTAATAAAGGTGTTGCAACAACATGG<br>TTAGCAACTAAACAACCTAAACAAAATCAATTAGATTTACCAACTGCACCAATATTT<br>ATACGTAAATCACAATTTCGTTTACCAACAAAAACACAAACACCAATTATTATGATT<br>GGTCCTGGTACTGGTTTGGCACCGTTCCGTGGTTTTATACAAGAACGCAATCTAGCTA<br>ATGATGAAGGTAAAACTGTTGGCGAAACAGTTTTATATTTCGGTTGTAGAAAACGTA<br>CGGAAGATTTCTTATATGAAGACGAATTAGTGAAATATGAAAAGGATGGTATCATTA<br>AATTACATATTGCTTTCAGTCGTGATCAACCACAAAAAGTATATGTTACACATTTAGT<br>TGAACAAAATGCTGATGAAATTTGGAGGATTATCGGTGAAAATAATGGACATCTCTA<br>TATTTGCGGTGATGCTAAATCGATGGCATCAGATGTACGTAATATTGTTTTGAAAATA<br>TTTAAAGAAAAAGGTCAAATGACGGAAGAACAAGCTTTAGCGTATTTGAAGAAGAT<br>GGAAACACAAAAACGTCTATCAGCTGATGTTTGGAGTTAA |
| CYP347BT1 | ATGATTTTGATAAGTCTAGTTCTAGTATTGTTAATTACAGTTTATTTTCTGTTAAAACG<br>TAAATATGAATATTGGAGAAAACGTGGAATACCTGGCCCTCAACCACGTTTCATTAC<br>TGGTAATGTTGGATCAATGCTATTAATGAAACAATCATTAGCTGATATTATTTCAAAA<br>TTATACAACGAATATCACAATGTTTCTTTAGTGGGAATGTTTAAAGGCCTATCGCCAA<br>CATTACTTGTTCGTGATCCTGATCTAATTAAAGATATATTAATCAAAGATTTTAGTTA<br>TTTCCACGATAATGATTTAGACGTAAGTGAATCTGTTGATCCATTACTTGGTCGAAAT<br>CCATTTGTATTAAAAGGTGAGAAATGGAAAATTGTACGTACACTACTAACACCTTGC<br>TTCACCAGTGGAAAGATGAAATCTATTCATCCTTTATTAGAAGATATCAGTCAAAAA<br>CTTGTAAAATATATTAAACGACAATTAGATGGGAATCGACGGAATGGTCTTGAACTT<br>AAAGAATTATGTAAAAGTTATACATTAGAAACAGTAGCTAGTTGTGCATTTGGTTTA<br>GAAGGGAAATGTTTCGAAGATGAAAATTCTGAATTTCGACAAATTGCAAACAAATTT<br>TTTGTACCAGGTGGATGGACAAATTTTATTATGATTTTATCAAGTATAATACCATCAT<br>TACCTACATTATTTCGTGTAAAAGTGGTACTGAAAGAAGTTGAAGATCGTCTCATTGC<br>ATTAACAAGTGAAACAATAAAATATCGTAAAGAAAACAACATTGAACGAAATGATT<br>TTATTCAAATTGTTAGTAAATTAAAAGAAACATCAAAAAATTATGAATTTACTGATG<br>TTGATGTATGTGCCCAAGCAGCGTCTTTTTTCGTTGATGGTTTTGAAACGACAGCAGT |

| | |
|---|---|
| | TACAATGAGTTTCCTTCTATTTCACTTAGCTGATAATCCACATATTCAAGAAAAGCTT<br>CGTGAAGAGATTGATGAACAATTCCGACAAAATAATGGAAAATTAACTTATGATAGT<br>ATTCAACAAATGGCATATCTTGATGCTGCACTTAATGAATCGCAGCGAATTCATCCA<br>GCTGCACAAATATTGTTGAAACGTTGCACACGAAATTATAAATATATTCCGAAAGCG<br>AATGATGGATTTGATAAACCACTTGAAATCGAGGCGGGAACTAGCCTGATAATATCA<br>ATAAGGAATTTACACAATGATCCCAAATACTTCAATGATCCAAGTGTATACAATCCG<br>GATAGATTTTTGGGTAAAAACAAAGAAAACTTGAATAAAAATGTATTCTTTCCATTT<br>GGAGATGGCCAACGGGTTTGTTTAGGTCAAAGATTTGCTCTTCTTCAAATAAAAATA<br>GGAATTGCTTATATTATACGTAATTTCAAACTGTCTGTGAATAGCAAAACACAGAAA<br>CCTATAAAGCTTGATCCTCTCTATCTGATGCCAGCACCTATTGGGGGATTTTGGATTA<br>ATTTTGAAGAAAGGGAACTATAA |
| CYP345BZ1 | ATGTTAATAGTTTTGGTTAATGTCTGTATTGTGTTGTCACTTATTTATTATCTTTTGAA<br>GAGGAATAATAACTATTGGAAATCACGAGGTGTTCCTCAAGATAAGCCATTTCTATT<br>GTTTGGAAGTTTCTACAATGTAGTTGCAGGAAAACAAAGTATACTTCAACGCATCAC<br>TGAAATCTACCAAAATTACAGAGATTTACCATATTTTGGTTTTTATACATTCACAAAA<br>CCAGTATTAATGATAAACAATCCTGATATTATCAAACGTATATTAGTGAAAGATTTTG<br>ATCAATTTGCTGATCGTAATGTTCATACAAATGAAAAAAGTGATCCCATTGGACATC<br>ATTCAATGTTTACCACACGCGGTAAGACATGGCGTAGTCTACGTACGAAAATATCAC<br>CGATTTTCACATCCGGCAAGATGAAGATGATATTGCCATTAATGATGGAATGTGGTG<br>AAAATATGTATCGAATTTTGAAAAATTCGGAACGCTCAATAGTTGATGTGCAAGATT<br>TAATAAAACGATATTCAGTTGATGTCATATCATCATGTGCTTTTGGTATAAATGCAAA<br>TAGTTTAAATAGTCAAAATGGTGGAATATTAGGTGCTGCTGAAAAATTAATGGATCA<br>AAATTCATTCAAACGCAGTTTTTCATTGTTTTGCTTCTTTTTCGTACCGAAATTGGCTA<br>ATATTCTACATTTGAAAATGTTCGATATGACGGCTGCCAATTTTCTAAGAAATATCTT<br>CACAACAACATTAGATGAAAGGGAGAAATTGAATATTCAACGGACTGATTTGATTGA<br>TTTATTGAATCAATTGAAGAAAACTCAATCGATTTCTGATGAATATAAATTTGATGAT<br>GATAAATTAGCAGCACAAGCTATGACTTTTTTCAGTGCTGGTAATGAAACAACATCA<br>ACAACTATTGCTTCTACTTTACATGAATTAGCTTTAGATAAAGCTGTACAAAGTCGAT<br>TAAGAGATGAAATTCGGGAATCTTATGAAACGAATAAAGGCTTCACTTATGAAGGAA<br>TCCAAGAAATGAAATACCTAGATATGGTCTTTAATGAAACCCTCAGACGGTACCCAG<br>TGACAATATTTCTAACACGAGAAGCTATTGACGATTATGTTATTGAGGAGACTGGAT<br>TTAAAATTGAAAAAGGTACATCGATCATGATACCGGTGTGTGGTTTACATTTCGATCC<br>TGAATATTTCCCAAATCCACAAAAATTCGATCCGGAACGTTATAGTGATGAAAACAA<br>AATTAATATTAAACCGTATACGTATATGCCTTTCGGAGAAGGTCCACGTAACTGTATT<br>GGTCGACGATTTGGTTTATTATCAGCAAAAATTGCACTTGTTCATCTACTGAAAGACT<br>TTGAATATGATGTGGCTGAAGACACACCAGTTCCATTAAAACTCAAACCTAAAGCTA<br>TTTTGTTACAACACGATGGTGGAATTCCTCTAAGAATCAACGCAGTTTAG |

**Supplementary Table 4. Comparison of absolute gene expression values (RPKM) for larvae and adult females.**

| Gene ID | Gene annotation | Absolute Expression (RPKM) | |
|---|---|---|---|
| | | Larvae | Adult |
| 490872 | P450 | 965.0 | 16.7 |
| 417025 | P450 | 575.8 | 0.9 |
| 1078326 | P450 | 154.9 | 11.0 |
| 727319 | P450-like | 99.9 | 0.0 |
| 983480 | P450 (**CYP347BT1**) | 76.4 | 2.7 |
| 566517 | P450 | 76.1 | 4.3 |
| 1547317 | P450 | 57.0 | 7.9 |
| 352995 | P450 | 51.5 | 0.6 |
| 634655 | P450 | 50.4 | 3.6 |
| 622392 | P450-like | 46.2 | 6.3 |
| 364323 | P450-like | 42.8 | 5.2 |
| 775472 | P450-like | 20.1 | 0.5 |
| 325667 | P450 (**CYP345BZ1**) | 18.8 | 0.1 |
| 561969 | P450 | 18.2 | 0.1 |
| 54557 | P450-like | 16.8 | 0.1 |
| 240516 | P450 reductase (**MpRed**) | 15.6 | 0.6 |
